# Activity-based profiling of bacterial RNA-modifying enzymes reveals species-specific 5-methyluridine modification in *Bacillus subtilis* 23S rRNA

**DOI:** 10.64898/2026.04.17.719261

**Authors:** Qais. Z. Jaber, Nathan J. Yu, Isao Masuda, Ya-Ming Hou, Ralph E. Kleiner

## Abstract

RNA modifications play an important role in biological processes. Mapping the diversity of RNA chemistry and studying the biological function of individual modifications remains an outstanding challenge in many organisms. In particular, RNA modifications remain poorly studied across most bacterial systems. Our group previously developed RNA-mediated activity-based protein profiling (RNABPP), a reactivity-based strategy employing metabolic labeling and quantitative proteomics to profile RNA modification writer enzymes in human cells. Here we adapt this approach to characterize RNA-modifying enzymes in bacteria. We apply metabolic labeling with 5-fluoropyrimidine nucleosides and phase separation-based enrichment of RNA-protein complexes (RNABPP-PS) to profile RNA pyrimidine modifying enzymes in *E. coli* and *B. subtilis*. We identify known and putative bacterial pyrimidine C5 methyltransferases, pseudouridine synthases, and dihydrouridine synthases, demonstrating the utility of our approach. Further, we find the carboxymethylaminomethyluridine (cnmn^5^U)-forming enzyme MnmG (GidA), supporting the existence of a covalent protein-RNA intermediate during the catalytic cycle. Finally, we use RNABPP-PS in *B. subtilis* to identify YfjO, an uncharacterized protein that is homologous to 5-methyluridine (m^5^U) methyltransferases. We use nucleoside and oligonucleotide mass spectrometry to establish that YfjO (which we rename as RlmS) installs m^5^U620 in the 23S rRNA (U576 in *E. coli*), a modification specific to the *B. subtilis* ribosome. We characterize Δ*yfjO B. subtilis* which has impaired cell proliferation and protein translation rate compared to WT. Taken together, our study establishes a versatile platform for RNA modifying enzyme discovery and characterization in bacteria and illuminates species-specific rRNA modification chemistry in *B. subtilis*.

## INTRODUCTION

Post-transcriptional or epitranscriptomic modifications on RNA play an important role in biological processes (1, 2). RNA modifications are found throughout all kingdoms of life and on all known RNA species, underscoring the importance of post-transcriptional regulation of RNA chemistry for RNA function (3–5). Despite decades of study, there are still major gaps in our understanding of the chemical identity, transcriptome-wide distribution, and cellular function of RNA modifications across diverse biological systems.

The best studied RNA modifications are those that occur on rRNA and tRNA. Studies of tRNA modifications in *E. coli* and *S. cerevisiae*, which date back to the 1960s (6), are thought to have revealed the entire complement of tRNA modifications and corresponding tRNA modifying enzymes in these model organisms (7, 8). More recent work using nanopore-based sequencing (9, 10) and cryogenic electron microscopy (cryoEM) (11, 12) have globally mapped rRNA modifications and their dynamics. Nevertheless, even in well-studied systems such as *E. coli*, new insights into tRNA (13) and rRNA modifications (14) and their associated writer enzymes continue to emerge. Moreover, their remain major gaps in our understanding of the biological and biochemical function of well-established tRNA and rRNA modifications, as well as the enzymatic mechanisms used by diverse RNA-modifying enzymes.

Bacteria are extremely versatile organisms that can grow in different environments and rapidly adapt to their surroundings. The diversity of RNA modifications in bacteria exceeds that found in eukaryotes (7), and in addition to roles of tRNA and rRNA modifications in translation fidelity (15, 16) and ribosome assembly (17–19), respectively, a number of bacterial RNA modifications show dynamic regulation in response to environmental stress and nutrient availability (20–22). In pathogenic bacteria, RNA modifications have been implicated in antibiotic resistance (23, 24), growth (25), and virulence (26–28). Comprehensive surveys of bacterial RNA modifications using advanced liquid chromatography-tandem mass spectrometry (LC-MS/MS) and modification-specific sequencing platforms are only beginning to emerge across diverse bacterial species (29–31), and further work in this area should provide new antibacterial drug targets and better understanding of mechanisms regulating bacterial growth, adaptation and pathogenicity.

Our lab previously developed RNA-mediated activity-based protein profiling (RNABPP) (32–34), a reaction-mechanism-based approach to discover and characterize RNA-modifying enzymes using chemoproteomics and modified nucleoside probes. Our initial applications of this method were in human cells, where we used oligo(dT)-pulldown-based enrichment and C5-modified pyrimidine nucleosides (32, 33) to capture diverse classes of RNA-modifying enzymes. Here we adapt our strategy to provide the first comprehensive profiles of 5-fluoropyrimidine-reactive RNA modifying enzymes in Gram-negative and Gram-positive bacteria. Our approach captures known and putative C5-pyrimidine methyltransfererases, dihydrouridine synthases (DUS), and pseudouridine synthases (PUS) in *E. coli* and *B. subtilis*, and provides evidence for a novel 5-fluorouridine (5-FUrd)-mediated crosslink between the carboxymethylaminomethyluridine (cmnm^5^U)-forming enzyme MnmG and its RNA substrate. Finally, we identify a species-specific 5-methyluridine (m^5^U) modification on *B. subtilis* 23S rRNA installed by the previously uncharacterized methyltransferase enzyme YfjO and we investigate the role of the *yfjO* gene in *B. subtilis* physiology. Taken together, our work provides a general reactivity-based approach to study prokaryotic RNA modifying enzymes and reveals new insights into the enzymology and biological function of epitranscriptomic regulators.

## RESULTS

### Metabolic labeling with 5-fluoropyrimidines in bacteria

To profile RNA-modifying enzymes in bacteria using activity-based nucleoside probes and mass spectrometry-based proteomics, we sought to apply our recently developed RNABPP-PS (Figure 1a) (34) method with 5-fluoropyrimidines. We previously established RNABPP-PS in human cells using 5-halopyrimidines and showed that aqueous-organic extraction with the OOPS method (35) (i.e. “phase separation”) can efficiently enrich mechanism-based RNA-protein crosslinks from intact cells. In contrast to our original RNABPP method that relied upon oligo(dT)-enrichment (32, 33) and is therefore biased towards polyadenylated RNAs that are primarily found in eukaryotes, RNABPP-PS can be applied broadly across different RNA classes and should therefore be suitable for activity-based profiling in prokaryotes.

**Figure 1.**
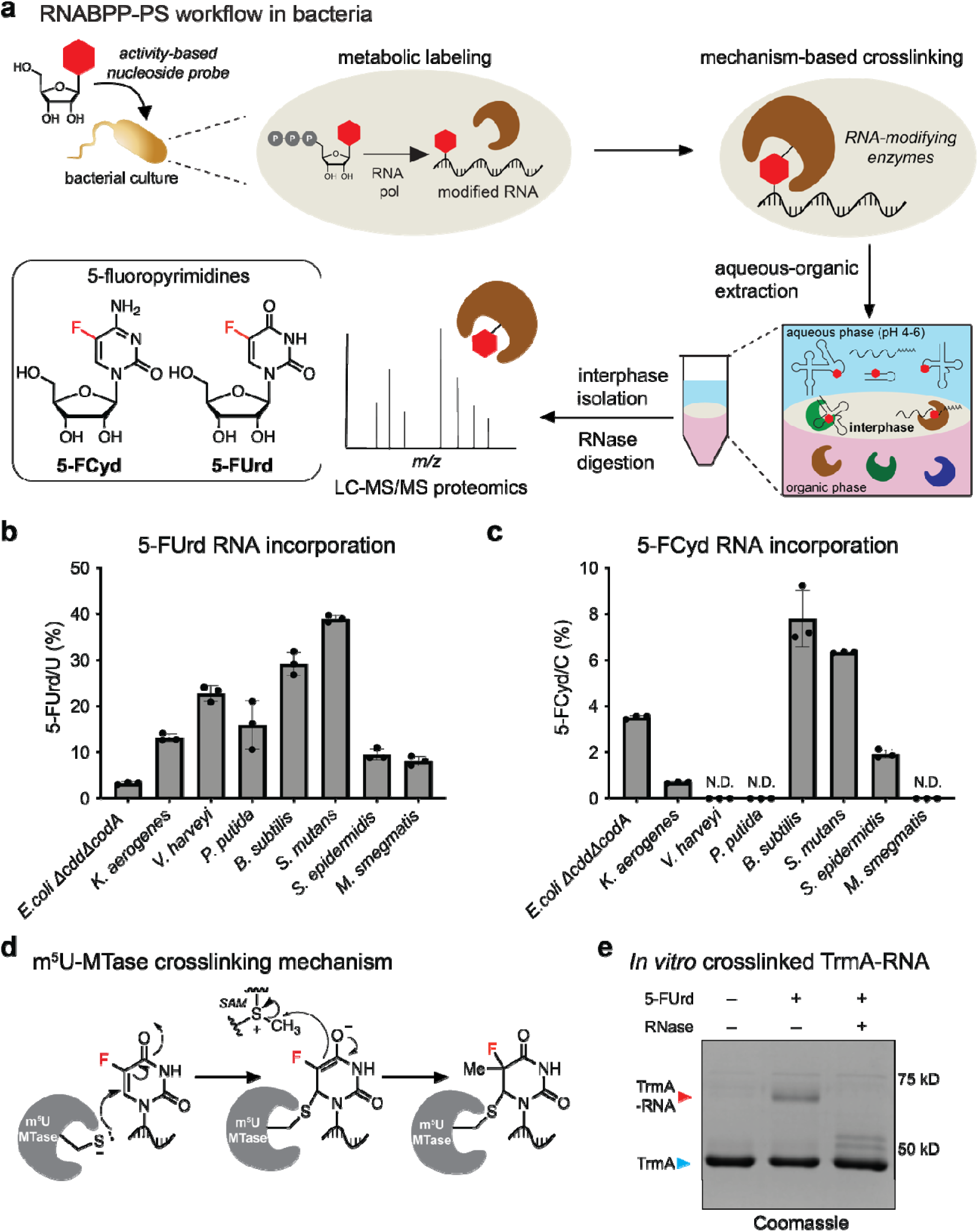
RNABPP-PS enables the capture of RNA-modifying enzymes in bacteria. **(a)** Schematic representation of RNABPP-PS and 5-fluoropyrimidine structures used in this study. Metabolic labeling of bacterial cells with an activity-based nucleoside probe facilities mechanism-based crosslinking to form covalent RNA-protein complexes. Crosslinked RNA-modifying enzymes are enriched through aqueous-organic extraction, followed by RNase digestion and identification by mass spectrometry-based proteomics. **(b)**, **(c)** Quantitative LC-QQQ-MS analysis of 5-FUrd and 5-FCyd incorporation levels in total RNA after metabolic labelling across a panel of bacteria. Values represent the percentage of 5-FUrd or 5-FCyd relative to total Urd or Cyd, respectively. Cells were treated with 5-FUrd or 5-FCyd at mid logarithmic phase OD = 0.5 and incubated until reaching stationary phase (OD ∼ 2). Three independent biological replicates were analyzed; data represent mean values ± s.e.m.; N.D., not detected **(d)** RNA-protein crosslinking mechanism between 5-FUrd-labeled RNA and m^5^U RNA methyltransferase (MTase). **(e)** Coomassie-stained SDS-PAGE analysis of RNA-protein crosslinking after *in vitro* methylation reaction between recombinant TrmA and isolated small RNA from 5-FUrd-treated *E. coli* Δ*trmA* cells.

We first explored metabolic labeling of bacterial RNA with 5-fluoropyrimidine nucleosides due to their compatibility with nucleotide salvage pathways (36, 37) and ability to form mechanism-based crosslinks with multiple classes of RNA modifying enzymes (32, 38–40). We evaluated the effect of 5-FUrd and 5-fluorocytidine (5-FCyd) on the growth of eight bacterial species from different genera, including Gram-positive, Gram-negative, and a mycobacterial strain (Table S1). All but one of the tested species, *P. putida*, contain a uridine kinase (Udk) homologue (41), which should mediate salvage of modified ribopyrimidines. In *P. putida*, uridine salvage may proceed through pyrimidine/purine nucleoside phosphorylase (*ppnP*) (42) and uracil phosphoribosyltransferase (*upp*) (43), however such a pathway is unlikely to enable direct metabolism of cytidine ribonucleosides. For experiments with *E. coli*, we generated a strain lacking known cytidine and cytosine deaminase enzymes (Δ*cdd*Δ*codA*) (44, 45) in order to prevent metabolic conversion of 5-FCyd to 5-FUrd. Addition of 5-fluoropyrimidines to bacterial cultures at concentrations ranging from 10 to 500 µM generally resulted in growth inhibition (Figure S1, S2), however the specific sensitivity varied among the evaluated bacterial species. Most strains showed comparable sensitivity to 5-FCyd and 5-FUrd with the exception of *E. coli* Δ*cdd*Δ*codA* and *P. putida,* which were both more sensitive to 5-FUrd.

We selected the maximum nucleoside concentration that exerted minimal effect (∼30% inhibition or lower) on bacterial growth and assessed metabolic RNA labeling in total cellular RNA using nucleoside LC-QQQ-MS (Figure 1b, 1c; Figure S3, S4, S5; Table S2, S3, S4). After treatment of bacteria with 5-FUrd, we measured RNA incorporation rates ranging from 3.3% to 38.9% (normalized to U) (Figure 1b), comparable or greater than incorporation levels achieved in our previous RNABPP experiments in human cells (32, 34). Low-level (0.02-0.3%) metabolic conversion between 5-FUrd and 5-FCyd (i.e. the presence of 5-FCyd in the RNA after 5-FUrd feeding) was detectable in *B. subtilis*, *S. epidermidis*, *V. harveyi* and *S. mutans* strains (Figure S5). This likely occurs at the nucleoside or nucleotide level and does not prevent efficient accumulation of 5-FUrd in the RNA. Metabolic labeling with 5-FCyd was generally less efficient than for 5-FUrd, although measurable incorporation between 0.7% to 7.8% (normalized to C) (Figure 1c) was observed in all but three strains (*V. harveyi*, *P. putida*, and *M. smegmatis*). In *V. harveyi* and *M. smegmatis*, this is likely due to cytidine deaminase activity as indicated by incorporation of 5-FUrd after 5-FCyd feeding (Figure S5). Indeed, most strains accumulated substantial levels of 5-FUrd (ranging between 4.2% to 23.5%) after 5-FCyd feeding with the exception of *P. putida*, suggesting that 5-FCyd is incompatible with its metabolism. In the *E. coli* Δ*cdd*Δ*codA* mutant strain we observed 3.5% incorporation of 5-FCyd with no deamination to 5-FUrd, and 3.3% incorporation of 5-FUrd, indicating success of the double knockout strategy. Overall, our LC-MS measurements of metabolic RNA labeling in diverse bacteria suggest that RNABPP-PS with 5-fluoropyrimidine nucleoside probes is likely to be a promising strategy for profiling RNA pyrimidine modifying enzyme activity in prokaryotes.

### RNABPP-PS with 5-fluoropyrimidines in *E. coli*

After validating metabolic labeling in multiple bacterial species, we piloted enrichment of a known 5-FUrd-reactive *E. coli* enzyme, the ubiquitous tRNA m^5^U54 methyltransferase TrmA (Figure 1d) (46), using phase separation. This enzyme is homologous to the human TRMT2A, previously isolated in our RNABPP studies utilizing 5-FUrd in human cells (32, 34). We performed an *in vitro* methylation reaction between purified TrmA enzyme and small RNA isolated from 5-FUrd-treated Δ*trmA E. coli* and detected a crosslinked band that remained stable under denaturing conditions and was susceptible to RNase treatment (Figure 1e; Figure S6). When formed in cells treated with 5-FUrd, this complex could be enriched in the interphase during aqueous-organic extraction (Figure S6), validating our ability to enrich bacterial 5-fluoropyrimidine-reactive proteins using RNABPP-PS.

Next, we prepared RNABPP-PS samples (Figure 1a) from *E. coli* Δ*cdd*Δ*codA* treated with either 10 µM 5-FUrd, 50 µM 5-FCyd, or left untreated as control. Three independent biological replicates were analyzed for each condition using label-free proteomics and data-independent acquisition (47, 48). In the *E. coli* 5-FCyd RNABPP-PS experiment (Figure 2a; Table S5; Supplementary Datafile 1), we found all three known 5-methylcytidine (m^5^C) methyltransferases (49) – RlmI, RsmB, and RsmF – significantly enriched between 11.3- to 97.0-fold. No other protein demonstrated a similar level of enrichment and only one other protein showed >4-fold enrichment – nirB, a nitrite reductase (50) which lacks homology to RNA modifying enzymes. As anticipated based on our experiments in human cells (32, 34), we found a broader range of enzymes enriched in the 5-FUrd *E. coli* RNABPP-PS experiment including all three known m^5^U methyltransferases (46, 51) RlmD, RlmC, and TrmA, all three known dihydrouridine synthases (DUS) (52) DusA, DusB, and DusC, and nine different pseudouridine synthase enzymes (PUS) (53) – TruA, TruB, RluA, RluB, RluC, RluD, RluE, RluF, and RsuA (Figure 2b; Table S5; Supplementary Datafile 1). Only two reported *E. coli* PUS enzymes, TruC and TruD, were not enriched in our study (Table S5). Notably, previous RNABPP experiments with 5-fluoropyrimidines in human cells did not capture PUS enzymes (32, 34). The literature provides evidence of high-affinity RNA-protein complex formation between bacterial PUS enzymes and 5-FUrd-modified RNA (54), however mechanistic studies have shown that this complex is either non-covalent or non-mechanism-based (55, 56). Although our RNABPP-PS experiment does not provide direct evidence of PUS-5-Urd crosslinking, we speculate that a sufficiently high-affinity RNA-protein species could survive the aqueous-organic workup. Finally, we also observed enrichment of MnmG, which is involved in the deposition of tRNA wobble uridine (U34) modifications (57). To our knowledge, mechanism-based adducts between MnmG-type enzymes and 5-FUrd-modified RNA have not been reported previously and are explored in more detail below. Overall, application of RNABPP-PS in *E. coli*, an organism where tRNA and rRNA modifying enzymes have been extensively studied, captured most of the expected enzymes and establishes a strong precedent for the use of this method in prokaryotes.

**Figure 2.**
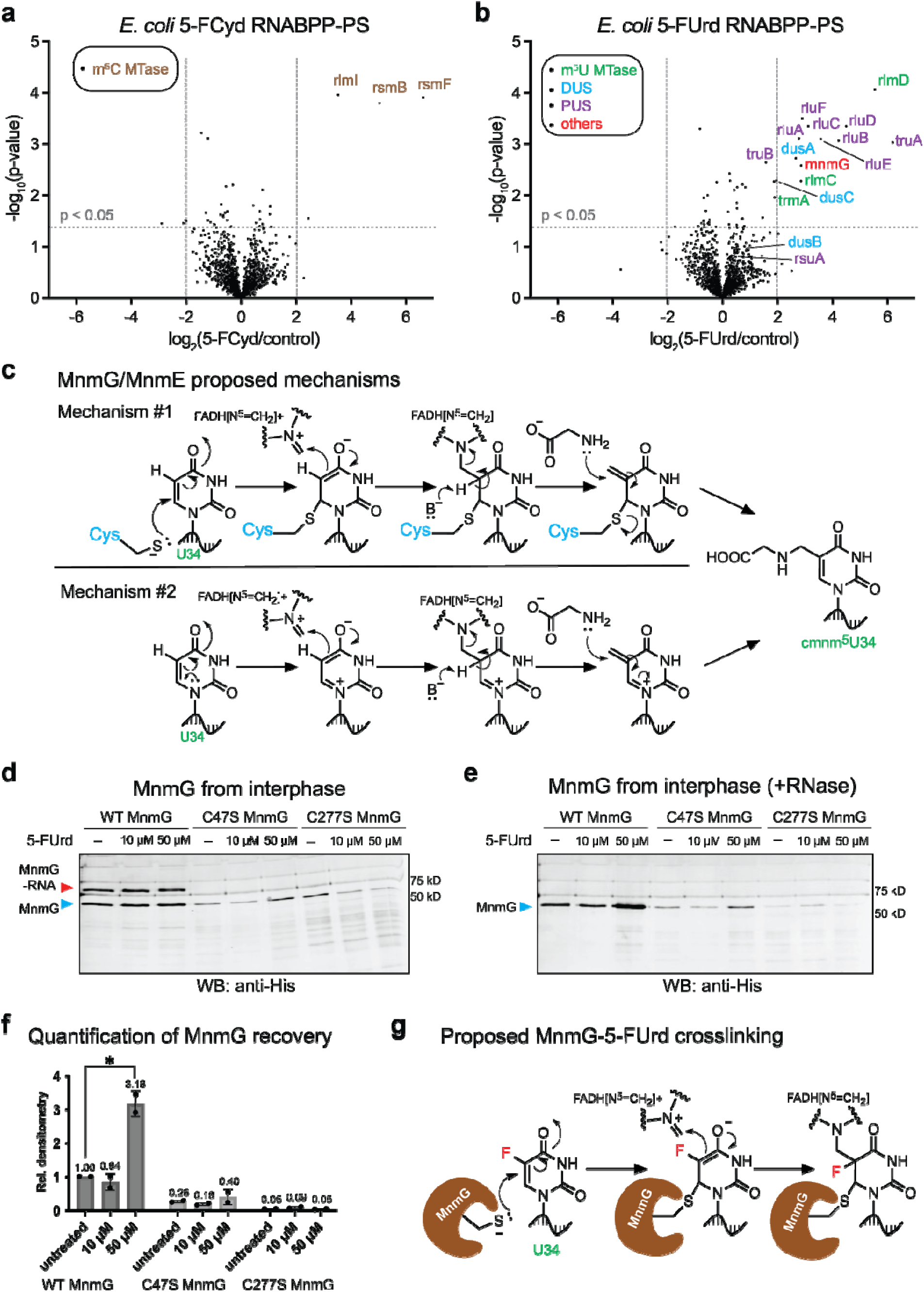
RNABPP-PS with 5-fluoropyrimidines in *E. coli* and characterization of a MnmG-RNA adduct. **(a), (b)** Proteomic profiling of 5-FCyd- and 5-FUrd-reactive proteins using RNABPP-PS in *E. coli*. Volcano plots are presented; expected and unexpected RNA-modifying enzymes are annotated and categorized. **(c)** Two proposed mechanisms for uridine activation during MnmG/MnmE reaction to install cmnm^5^U at position U34 in tRNA. **(d)** Western blot analysis of MnmG-RNA complex enriched in interphase samples following metabolic labeling with 5-FUrd. *E. coli* Δ*mnmG* cells expressing MnmG(WT)-6xHis, MnmG(C47S)-6xHis or MnmG(C277S)-6xHis were treated with 10 µM or 50 µM 5-FUrd for 6 hours or left untreated, followed by three rounds of interphase extraction. Two interphase extraction replicates were analyzed by Western blot using anti-His antibody. **(e)** Western blot analysis of MnmG recovery after metabolic labeling with 5-FUrd, interphase extraction, and RNase digestion. **(f)** Quantification of the relative densitometry of the recovered MnmG bands. Data are representative of mean values ± s.e.m.; unpaired *t*-test (two-tailed) were used to evaluate statistical significance. *P*-values: WT MnmG untreated vs. WT MnmG 50 µM 5-FUrd, 0.015486. **(g)** Proposed RNA-protein crosslinking mechanism between 5-FUrd-labeled RNA and MnmG enzyme.

### Characterization of a mechanism-based MnmG-RNA adduct

MnmG, also known as GidA (57), together with MnmE is responsible for the addition of a carboxymethylaminomethyl (cmnm^5^U) group to the wobble uridine (U34) residue of a subset of tRNAs. MnmG/MnmE collaborate to facilitate tRNA modification, utilizing glycine as a substrate and several cofactors, including GTP, FAD, NADH, and methylenetetrahydrofolate (5,10-CH_2_-THF). Recent studies by Bommisetti *et al.* (58, 59) have demonstrated that FADH_2_ is employed to transfer a methylene from 5,10-CH2-THF to the activated C5 uridine atom through a covalent FADH-iminium intermediate; subsequent nucleophilic attack by glycine results in the formation of the cmnm^5^U nucleotide (Figure 2c). Two mechanistic hypotheses were presented for uridine activation prior to methylation: 1) attack of a Cys residue from MnmG/MnmE at the C6 position, or 2) internal activation via the N1 nitrogen without Cys involvement. To better characterize a putative MnmG-RNA covalent adduct, we expressed recombinant MnmG-6xHis in Δ*mnmG E. coli* and performed Western blot following 5-FUrd treatment and aqueous-organic interphase isolation. We observed an RNase sensitive-band with higher mobility in the interphase fraction of samples treated and untreated with 5-Urd, suggesting the formation of a covalent MnmG-RNA intermediate (Figure 2d, 2e; Figure S7). In samples treated with 50 µM 5-FUrd, recovery of MnmG after phase separation and RNase digestion of the interphase was ∼3-fold higher than in the untreated sample (Figure 2f), suggesting that 5-FUrd can stabilize the MnmG-RNA complex. We further assayed the mechanism of complex formation by mutation of two conserved cysteine residues known to be essential for the activity of MnmG – C47 and C277 (60, 61). Neither C47S MnmG nor C277S MnmG mutants expressed in Δ*mnmG E. coli* generated bands consistent with crosslinking and consequently recovery of these mutant proteins after interphase isolation and RNase digestion was minimal in comparison to wildtype (WT) MnmG protein (Figure 2d, 2e; Figure S7). Taken together our data supports the formation of a covalent intermediate between MnmG and substrate RNA (Figure 2g). We were unable to identify the protein residue responsible for RNA-protein adduct formation as mutation of either catalytically essential Cys residue abolished crosslinking. However, we speculate that only one Cys residue performs nucleophilic attack on the Urd C6 position, while the other may be important for stabilizing the covalent intermediate or for earlier catalytic steps.

### RNABPP-PS with 5-fluoropyrimidines in *B. subtilis*

To broaden the application of RNABPP-PS beyond the extensively studied *E. coli*, we targeted the model Gram-positive bacteria *B. subtilis*. In contrast to *E. coli*, there exist major gaps in the characterization of *B. subtilis* RNA modifications and corresponding RNA modifying enzymes. 5-FCyd and 5-FUrd RNABPP-PS experiments in *B. subtilis* (Figure 3a; Figure S8; Table S5; Supplementary Datafile 2) identified a highly similar set of proteins, likely due to metabolic interconversion between 5-FCyd and 5-FUrd as shown above (Figure S5). Although this introduces some ambiguity in enzyme-substrate assignment, in practice it enables identification of Cyd and Urd modifying enzymes in a single proteomic experiment. For the sake of simplicity we will primarily discuss the results of the 5-FUrd RNABPP-PS experiment in *B. subtilis* (Figure 3a; Table S5). Similar to our profiling experiments in *E. coli*, we identified enzymes from multiple enzyme classes including m^5^U methyltransferases TrmFO (62) and RlmCD (YefA) (63), the only known *B. subtilis* m^5^C methyltransferase RsmB, both known DUS enzymes (64), Dus1 and Dus2, and seven PUS enzymes. Although only TruA, TruB and RluB have been well-characterized (65, 66), based on homology to *E. coli* these seven PUS genes represent all known PUS enzymes in *B. subtilis*. Similar to our findings in *E.* coli, we also enriched the *B. subtilis* MnmG homologue. Its identification with 5-FUrd further supports the occurrence of a covalent RNA-protein intermediate during MnmG catalysis as described in the previous section (Figure 2d, 2e).

**Figure 3.**
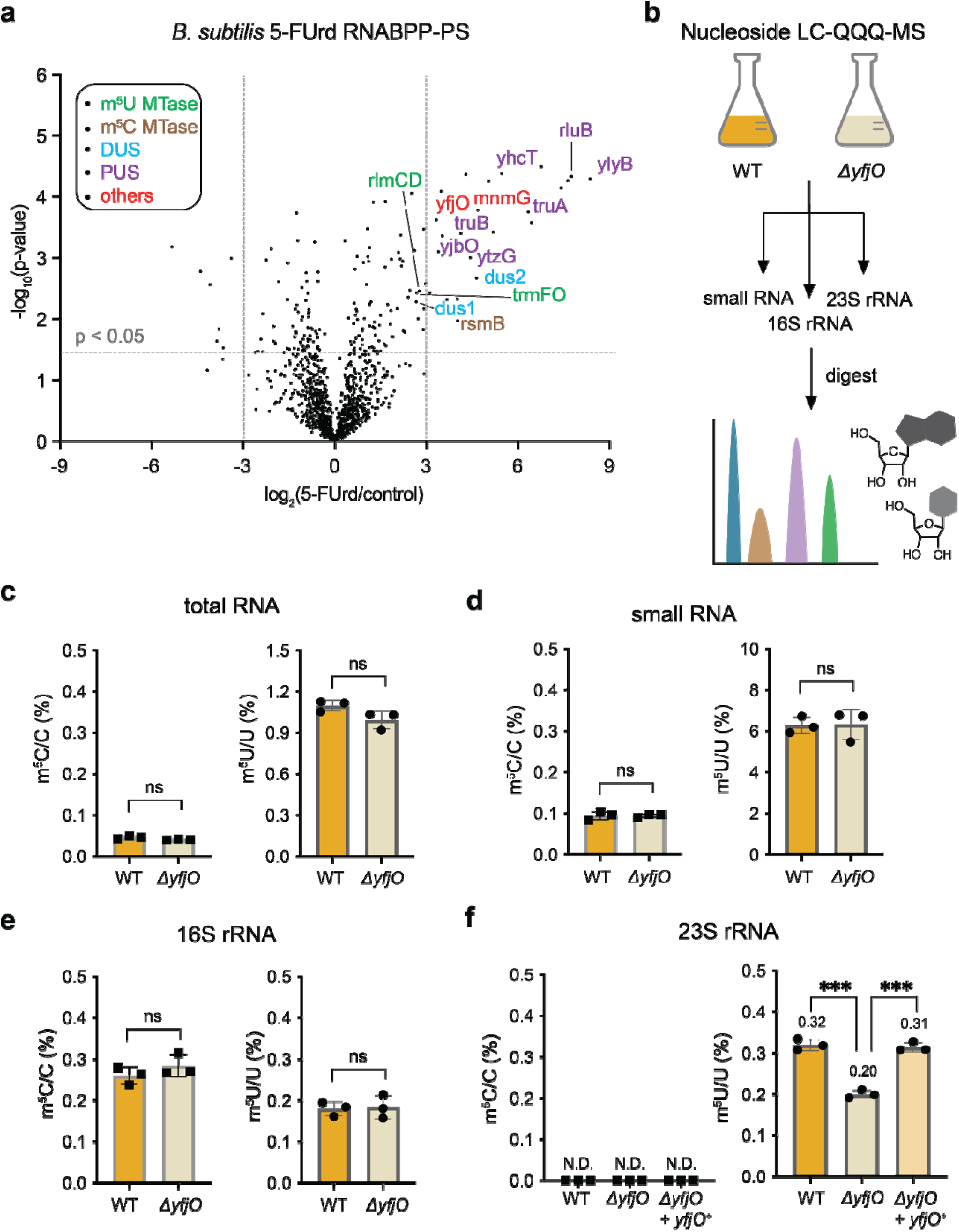
RNABPP-PS in *B. subtilis* and characterization of 23S rRNA m^5^U methyltransferase YfjO. **(a)** Proteomic profiling of 5-FUrd-reactive proteins using RNABPP-PS in *B. subtilis*. A volcano plot is shown; expected and unexpected RNA-modifying enzymes are annotated and classified. **(b)** Illustration of nucleoside LC-QQQ-MS workflow. Total RNA isolated from *B. subtilis* WT and *yfjO* knockout (Δ*yfjO*) strains was fractionated into distinct RNA species, digested to nucleosides, and analyzed by LC-QQQ-MS. **(c)-(f)** Nucleoside LC-QQQ-MS analysis of m^5^C and m^5^U levels across different RNA fractions: **(c)** total RNA; **(d)** small RNA; **(e)** 16S rRNA; **(f)** 23S rRNA. m^5^C and m^5^U levels in the *yfjO* add-back strain (Δ*yfjO+yfjO^+^*) were assessed in 23S rRNA. Three independent biological replicates were analyzed; data are representative of mean values ± s.e.m. Statistical significance was determined using unpaired *t*-test (two-tailed). *P*-values: WT vs. Δ*yfjO* in 23S rRNA, 0.000280; Δ*yfjO+yfjO^+^* vs. Δ*yfjO* in 23S rRNA, 0.000222.

In addition to characterized RNA modifying enzymes and homologues to known *E. coli* genes, our profiling experiment identified proteins without known function or lacking known association with RNA. For example, YbfG, and FadG, uncharacterized proteins with predicted peptidoglycan-binding domains were enriched. Multiple phage-like element PBXS proteins (67) were also identified. These proteins are known to be induced after the SOS response in *B. subtilis* (68). In addition, the N-acetylmuramoyl-L-alanine amidases XlyA/B, which are peptidoglycan-binding proteins and are associated with phage-like element PBXS, were similarly enriched (69). Although we cannot exclude physical associations between these proteins and 5-FUrd-modified RNA, we hypothesize that enrichment may be due to changes in gene expression upon 5-FUrd treatment combined with partitioning into the interphase layer due to their physicochemical properties or non-RNA-dependent interactions. Finally, YfjO, enriched 10-fold, is a predicted RNA methyltransferase, however its substrate is unknown and our RNABPP-PS experiment captured the two known RNA m^5^U methyltransferases in *B. subtilis* – TrmFO (62), which installs tRNA m^5^U54, and RlmCD (63), which installs m^5^U794 and m^5^U1968 on 23S rRNA, and the single known m^5^C methyltransferase, RsmB. We provide the first characterization of the biochemical and biological activity of YfjO in the sections below.

### Quantification of RNA m C and m U levels in *B. subtilis* Δ*yfjO*

A previous homology search (63) in *B. subtilis* using the *E. coli* 23S rRNA m^5^U methyltransferases RlmC and RlmD identified two *B. subtilis* orthologues – YefA, which has been renamed RlmCD, and YfjO. Desmolaize *et al.* (63) showed that RlmCD installs m^5^U at positions 794 and 1968 in the 23S rRNA (the homologous positions to those modified by RlmC and RlmD in *E. coli*), whereas YfjO does not participate in tRNA m^5^U formation or in m^5^U formation at the RlmCD 23S rRNA sites, leaving YfjO uncharacterized to date. The enrichment of YfjO in our 5-FUrd/5-FCyd RNABPP-PS experiments (Figure 3a; Figure S8) suggests either m^5^U or m^5^C modification activity. To investigate the role of YfjO enzyme in *B. subtilis* RNA modification and cellular physiology, we generated a *yfjO* knockout (Δ*yfjO*) using the double recombination strategy (70) in *B. subtilis* WT 168 (71) as the genetic background. We validated two independent knockout clones by targeted and whole genome sequencing to confirm *yfjO* deletion and any off-target genomic mutations (Table S6). In addition, we made an add-back strain (Δ*yfjO+yfjO^+^*) by introducing *yfjO* gene into the Δ*yfjO* strain at the *amyE* locus using an integrative plasmid (72).

We measured m^5^U and m^5^C levels in various cellular RNA species isolated from WT or Δ*yfjO* using nucleoside LC-QQQ-MS (Figure 3b). Our results showed no significant difference in the levels of m^5^C or m^5^U in total cellular RNA or the cellular small RNA fraction (RNAs < 200 nt) between WT and knockout strains (Figure 3c, 3d; Table S7). We measured ∼6% m^5^U in small RNA, consistent with one known tRNA m^5^U position (m^5^U54) and 14-22 uridines per tRNA. The m^5^C levels in small RNA were ∼0.1%. Considering there is no reported m^5^C modification in *B. subtilis* tRNA and that tRNA contains 18-27 cytidines, this suggests that measured m^5^C levels are likely due to contamination from 16S rRNA or occur on isolated tRNA species. Next, we purified 16S and 23S rRNA species and measured m^5^C and m^5^U content. In 16S rRNA, there is one known m^5^C modification (73) and no reported m^5^U modifications. We measured ∼0.25% m^5^C, consistent with stoichiometric modification at one of ∼365 cytidines, and 0.15% m^5^U, which equates to less than one m^5^U modification per ∼315 uridines in 16S rRNA (Figure 3e; Table S7). Importantly, both measured m^5^U and m^5^C levels in 16S rRNA were unchanged between WT and Δ*yfjO*, indicating they are not Δ*yfjO* substrates. In contrast, we measured a ∼30% decrease in 23S rRNA m^5^U levels in the knockout strain (Figure 3f; Table S7), and YfjO add-back restored m^5^U levels to those of WT. 23S rRNA contains ∼573 uridines and our measured m^5^U level in WT (0.32%) corresponds to modification at ∼3 uridines in the WT, with loss of a single m^5^U modification in the Δ*yfjO* strain. Taken together, these results suggest that YfjO is responsible for a single m^5^U modification within the 23S rRNA.

### Double-cleavage LC-MS mapping of YfjO target in *B. subtilis*

A recent LC-MS study of RNA modifications in the *B. subtilis* ribosome (73) identified a previously unreported m^5^U modification at position 620 in the 23S rRNA, in addition to the known m^5^U794 and m^5^U1968 sites. Notably, this m^5^U620 modification appears to be specific to *B. subtilis* rRNA and is absent from *E. coli* (Figure 4a). We therefore investigated whether YfjO is responsible for installing m^5^U620 in the 23S rRNA using a double-cleavage oligonucleotide LC-MS workflow (Figure 4b). Briefly, rRNA isolated from *B. subtilis* cultures was subjected to sequence-guided RNase H cleavage using a 2’-deoxy-2’-O-methyl chimeric oligonucleotide, and the resulting RNA oligonucleotide fragment spanning U620 was analyzed by LC-MS/MS. In WT *B. subtilis*, we detected an oligonucleotide mass consistent with methylation at U620: pAGAAm^5^UGAA (*m/z* = 1335.687, z = -2) (Figure 4c; Figure S9, S10, S11, S12). The corresponding unmodified oligo, pAGAAUGAA (*m/z* = 1328.687; z = -2), was not detected in WT 23S rRNA, consistent with the novel m^5^U620 site reported by Williamson and co-workers (73) and indicating that methylation at this position is stoichiometric. In contrast, RNA isolated from the Δ*yfjO* strain showed the opposite trend – we detected exclusively unmodified pAGAAUGAA oligonucleotide and none of the methylated species. Further, we recapitulated methylation at m^5^U620 in the genetic add-back strain. Finally, to evaluate the biochemical activity of YfjO directly, we purified recombinant protein by heterologous expression in *E. coli* and set up an *in vitro* methylation reaction containing recombinant YfjO, *S-*adenosyl methionine (SAM), and purified rRNA isolated from the Δ*yfjO* strain. Production of m^5^U620 was measured using double-cleavage oligonucleotide LC-MS as described above. As expected, a control reaction lacking YfjO yielded only unmodified oligo pAGAAUGAA (Figure 4d; Figure S13, S14) whereas addition of YfjO generated primarily the m^5^U-modified oligo (Figure 4d; Figure S13, S14). Together, these results demonstrate that YfjO is a SAM-dependent m^5^U methyltransferase responsible for installing m^5^U at position 620 of *B. subtilis* 23S rRNA.

**Figure 4.**
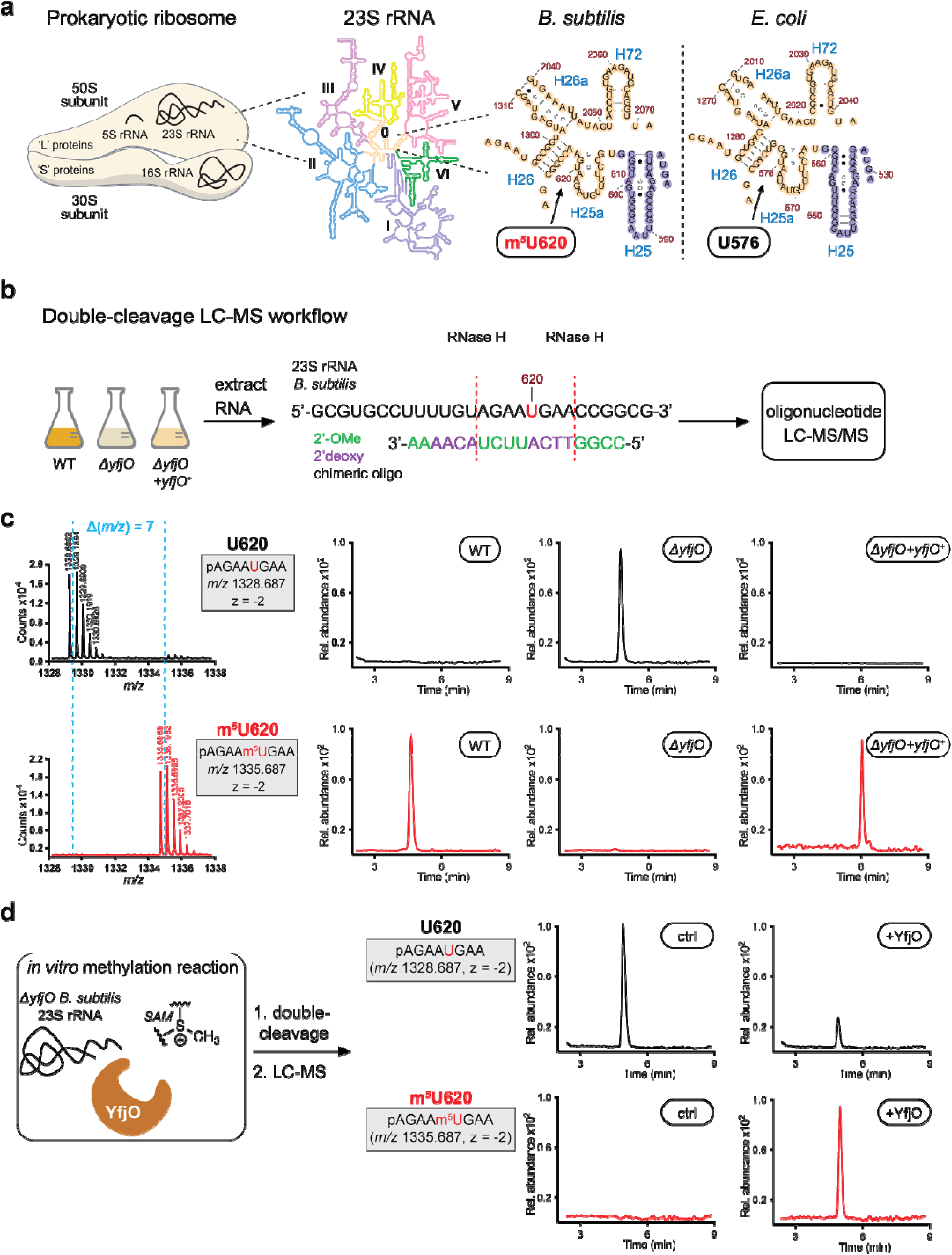
Double cleavage LC-MS pinpoints m^5^U620 in 23S rRNA as the YfjO target site. **(a)** Potential YfjO target within *B. subtilis* 23S rRNA. Shown are schematic of the prokaryotic ribosome and the secondary structure of 23S rRNA. The zoomed region highlights domain 0 and Helix 25a, including position U620 in *B. subtilis* 23S rRNA and the corresponding U576 in *E. coli* numbering. “—” indicates canonical base pair interactions (A-U and G-C), “•” indicates G-U interactions, and “ο” indicates A-G, U-C, U-U and A-A interactions. **(b)** Workflow for double-cleavage oligonucleotide LC-MS to map modification sites. 23S rRNA was isolated from *B. subtilis* cultures, subjected to sequence-guided RNase H cleavage using a 2’-deoxy-2’-O-methyl chimeric oligonucleotide, and the resulting oligonucleotide fragment was analyzed by LC-MS/MS. **(c)** Double-cleavage oligonucleotide LC-MS analysis of 23S rRNA from *B. subtilis* WT, Δ*yfjO,* and Δ*yfjO+yfjO^+^*strains. Shown are mass spectra with the sequence, *m/z* value, and charge state of each fragment (left), and extracted ion chromatograms of m^5^U-modified and corresponding unmodified oligonucleotide fragments produced from sequence-guided RNase H cleavage. **(d)** Recombinant YfjO reinstalls m^5^U620 *in vitro*. Double-cleavage LC-MS analysis of 23S rRNA after *in vitro* methylation reaction using recombinant YfjO and isolated 23S rRNA from *B. subtilis* Δ*yfjO*. Extracted ion chromatograms of m^5^U-modified and corresponding unmodified oligonucleotide fragments are shown. The sequence, *m/z* value and charge state of each oligonucleotide are shown on the left.

### Heterologous expression of YfjO in *E. coli*

We performed a BLAST search using the *yfjO* sequence to determine whether close homologues are present outside of *B. subtilis*. This analysis identified 1400 hits with >80% query coverage and >80% identity, all belonging to the *Bacillus* genus. Although the secondary structure surrounding domain 0 is conserved between *E. coli* and *B. subtilis*, the primary sequence in this region shows modest divergence (Figure 4a). Alignment of the cryo-EM 3D structures of the two ribosomes (74, 75) reveals an almost identical fold and close spatial correspondence in the region of interest (Figure 5a).

**Figure 5.**
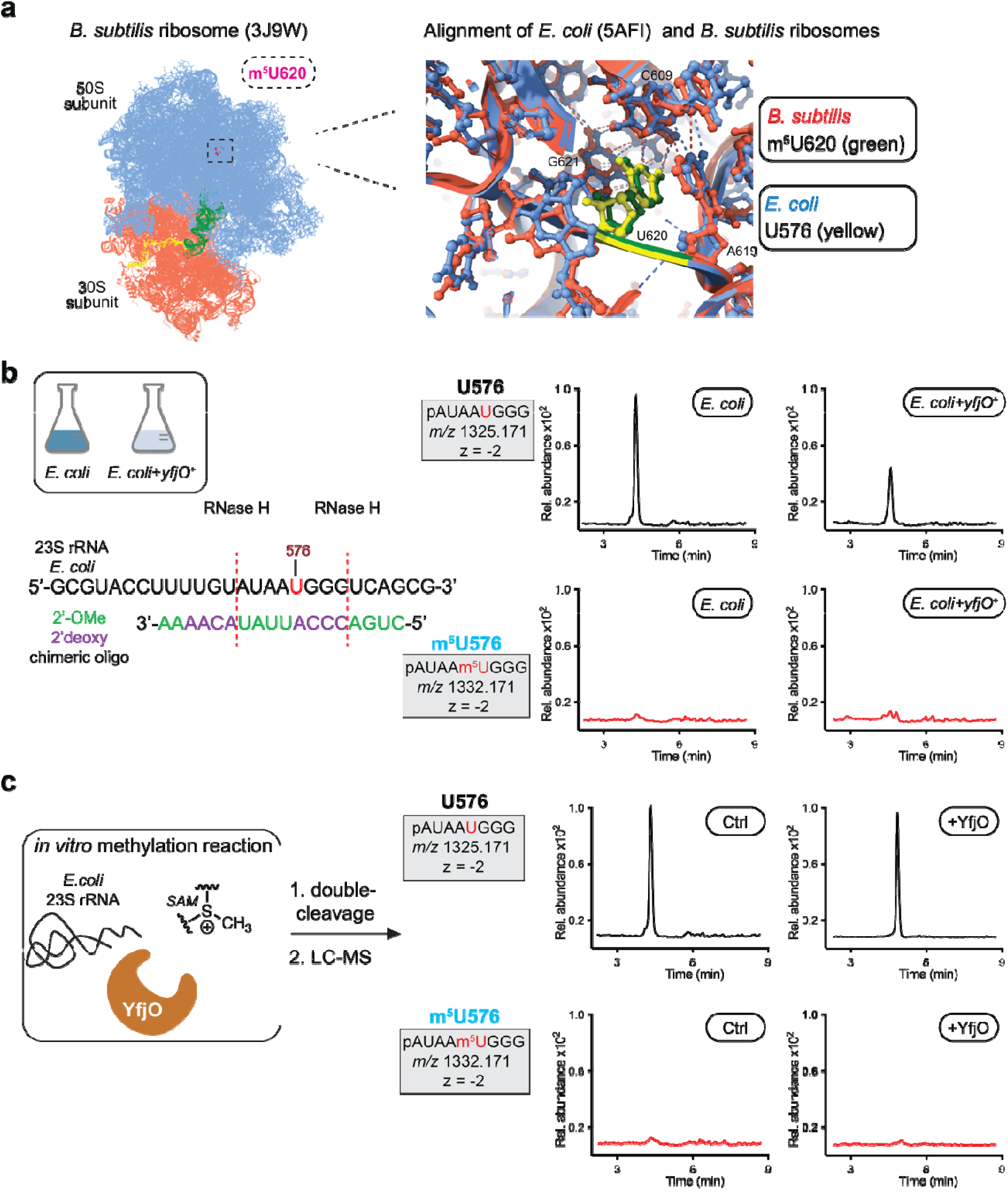
YfjO is unable to modify U576 in *E. coli* 23S rRNA. **(a)** Structural comparison of the YfjO target region in *B. subtilis* and *E. coli* ribosomes. Left: cryo-EM structure of *B. subtilis* ribosome, with the large subunit in blue, small subunit in red, tRNA in green, mRNA in yellow, and m^5^U620 modification highlighted in magenta. Right: zoomed-in view of the modification site after structural alignment of *B. subtilis* (Protein Data Bank (PDB) ID: 3J9W), and *E. coli* (PDB ID: 5AFI) 23S rRNA. The *B. subtilis* ribosome is shown in red, the *E. coli* ribosome in blue, with m^5^U620 depicted in green and the corresponding U576 in *E. coli* shown in yellow. Root Mean Square Deviation (RMSD) between 2206 pruned atom pairs is 0.954 Å (RMSD across all 2841 pairs: 5.904 Å) calculated using ChimeraX. **(b)** Double-cleavage oligonucleotide LC-MS analysis of *E. coli* 23S rRNA. Extracted ion chromatograms of m^5^U-modified and corresponding unmodified oligonucleotide fragments generated by sequence-guided RNase H cleavage from control *E. coli* BL21 and YfjO-expressing *E. coli* (*E. coli +yfjO^+^*). The sequence, *m/z* value and charge state of each oligonucleotide are shown on the left. **(c)** Double-cleavage LC-MS analysis following *in vitro* methylation reaction of isolated *E. coli* 23S rRNA with recombinant YfjO. Extracted ion chromatograms of m^5^U-modified and corresponding unmodified oligonucleotide fragments are shown, with oligonucleotide sequences, *m/z* values, and charge states indicated (left).

To further probe the substrate specificity of YfjO and assist in elucidating biological phenotypes, we evaluated whether YfjO can modify U576 in the *E. coli* 23S rRNA (the homologous position to U620 in *B. subtilis*) (Figure 4a and Figure 5a). First, we expressed YfjO in *E. coli* strain BL21 (DE3) and measured m^5^U modification on 23S rRNA by LC-MS. In both control and the YfjO- expressing *E. coli*, only the unmodified oligo pAUAAUGGG (*m/z* 1325.171, z = -2) was detected (Figure 5b; Figure S15, S16, S17). We then performed an *in vitro* methylation assay using purified YfjO enzyme and rRNA isolated from *E. coli*. Both the control and YfjO reactions yielded exclusively the unmodified oligo pAUAAUGGG (*m/z* 1325.171, z =-2) (Figure 5c; Figure S15, S16). These results indicate that YfjO is unable to install m^5^U modification at U576 in *E. coli* 23S rRNA, likely due to a requirement for local sequence context.

### Probing the biological role of YfjO in growth and protein translation

To investigate the role of YfjO in *B. subtilis* physiology, we first examined previously reported phenotypes for Δ*yfjO* strains from the Gross laboratory *B. subtilis* knockout collection (70): Δ*yfjO::kan* (BKK08020) and Δ*yfjO::erm* (BKE08020). Both strains were reported to exhibit modestly reduced growth, with relative fitness values of ∼0.9 at 37 °C in LB media in a genome-wide growth screen. In addition, both strains were reported to be cold sensitive, with reduced fitness values of 0.83 and 0.58 at 16 °C for Δ*yfjO::kan* (BKK08020) and Δ*yfjO::erm* (BKE08020), respectively, and reported to have reduced sporulation efficiency (70). We therefore obtained these strains from the *Bacillus* Genetic Stock Center and compared their growth to the WT 168 strain. Both knockout strains displayed growth defects relative to WT strain, but with distinct phenotypic patterns (Figure S18). The Δ*yfjO::kan* (BKK08020) strain exhibited normal exponential growth with a rapid die off of 20-25% of the population after entering stationary phase (Figure S18). In contrast, the Δ*yfjO::erm* (BKE08020) strain showed slower exponential growth but behaved similarly to WT upon reaching stationary phase. We also tested sporulation efficiency for Δ*yfjO::kan* (BKK08020) and found a significant ∼2/3 reduction compared to WT (Figure S19; Table S8). Because these two knockout strains exhibited different phenotypes despite targeting the same gene, we performed whole-genome sequencing on Δ*yfjO::kan* (BKK08020) strain. This analysis revealed additional genomic mutations, including changes in sporulation-associated factors (*prpC*, *bmrD*), ribosomal proteins (*rplW*), and motility-related genes (*swrAA*), that likely contribute to the observed phenotypes (Table S9).

We next characterized the Δ*yfjO* strains generated herein that were free of off-target genomic mutations (Table S6). We first measured growth at 37 °C (Figure 6a), 30 °C (Figure 6b), and 25 °C (Figure S20) in LB media. In contrast to the Δ*yfjO* strains from the *Bacillus* knockout collection, under these conditions both Δ*yfjO* clones exhibited indistinguishable growth from WT (Figure 6a, 6b; Figure S20). Likewise, sporulation efficiency measurements revealed no significant differences between our Δ*yfjO* strains and WT (Figure S19).

**Figure 6.**
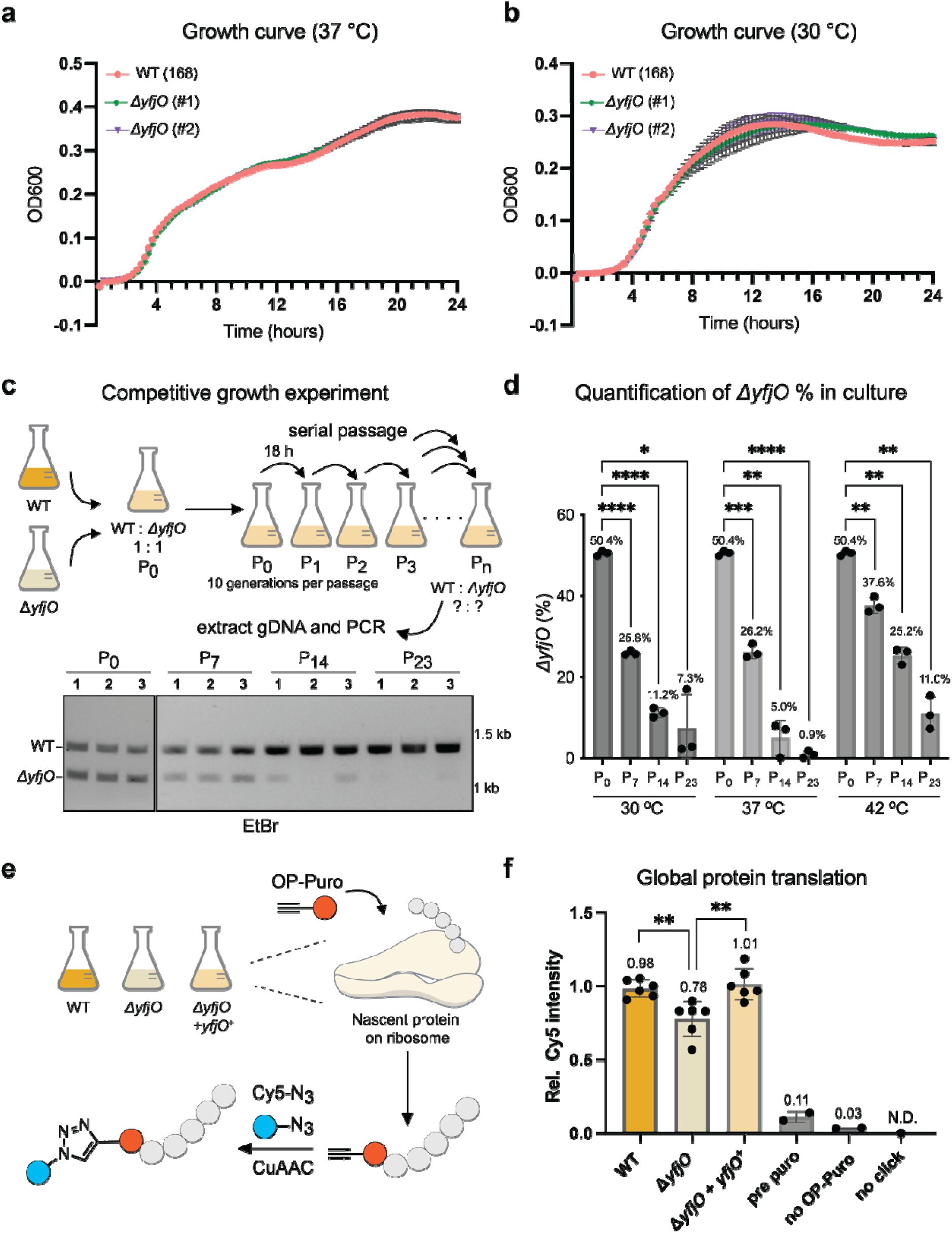
YfjO contributes to optimal cellular fitness and global protein translation capacity in *B. subtilis*. **(a), (b)** Growth curves of *B. subtilis* WT and two independent Δ*yfjO* clones at 37 °C or 30 °C. Cultures were monitored by OD600 measurement every 15 minutes for 24 hours using a plate reader. Two independent biological replicates with three technical replicates each were analyzed; data represent mean ± s.e.m. **(c)** Competitive growth assay of WT versus Δ*yfjO*. A 1:1 mixed culture was grown for 18 hours at 37 °C, followed by serial passaging (10 generations per passage). Genomic DNA was extracted, and the *yfjO* locus was amplified by PCR. WT cells producing a 1464 bp (upper band) band and the Δ*yfjO* strain producing a 1135 bp (lower band). **(d)** Strain abundance by quantification of PCR product densitometry from Passage 7, 14, and 23 at 30 °C, 37 °C, and 42 °C. Three independent biological replicates were analyzed; data are representative of mean values ± s.e.m. Unpaired *t*-test (two-tailed) were used to evaluate statistical significance. *P*-values: P_0_ vs. 30 °C P_7_, 0.0000008; P_0_ vs. 30 °C P_14_, 0.000017; P_0_ vs. 30 °C P_23_, 0.012094; P_0_ vs. 37 °C P_7_, 0.000612; P_0_ vs. 37 °C P_14_, 0.002629; P_0_ vs. 37 °C P_23_, 0. 0000009; P_0_ vs. 42 °C P_7_, 0.004521; P_0_ vs. 42 °C P_14_, 0.001069; P_0_ vs. 42 °C P_23_, 0.003123. **(e)** Illustration for global protein translation analysis. *B. subtilis* WT, Δ*yfjO*, and Δ*yfjO+yfjO^+^* cultures were labeled with OP-Puro at 37 °C for 30 minutes, followed by cell lysis and CuAAC click chemistry with Cy5-azide. Cy5 signal in nascent proteins was measured by fluorescent gel imaging. **(f)** Quantification of global protein translation. The median of relative Cy5 intensity and s.e.m. from six independent biological replicates are shown. No OP-Puro treatment, pretreatment with puromycin (Pre Puro), and no click controls were performed. Statistical significance was determined by unpaired *t*-test (two-tailed). *P*-values: WT vs. Δ*yfjO*, 0.003767; Δ*yfjO+yfjO^+^* vs. Δ*yfjO*, 0.004857.

Next, we performed competitive growth experiments between WT and Δ*yfjO* strains. A 1:1 mixture of WT and Δ*yfjO* cells was grown at the indicated temperature for 18 hours, followed by serial passaging. Genomic DNA was extracted at defined passages, and PCR amplification of the *yfjO* locus was used to quantify strain abundance, with WT cells producing a 1464 bp band and the Δ*yfjO* strain producing a 1135 bp band generated from the kanamycin resistance cassette incorporated during gene knockout (Figure 6c: Figure S21). The initial culture (P_0_) contained 50.4% Δ*yfjO* as measured by optical density and gel densitometry of genomic PCR products. By passage 7, the fraction of Δ*yfjO* cells decreased to 25.8–37.6%, and by passage 14 it further declined to 5.0-25.2%. After 23 passages, the Δ*yfjO* population was reduced to 0.9-11.0%. The strongest effect was observed at 37 °C, the optimal growth temperature for *B. subtilis*, where Δ*yfjO* cells were nearly eliminated by passage 23, leaving almost exclusively WT cells (Figure 6c, 6d; Figure S21; Table S10). Calculation of relative fitness at 37 °C revealed a relative fitness value of 0.98 for the Δ*yfjO* strain at 37 °C, indicating a subtle but measurable disadvantage during long-term competition. To assess whether loss of *yfjO* affects global protein synthesis, we measured O-propargyl-puromycin (OP-puro) (76) incorporation into nascent polypeptides in Δ*yfjO*, Δ*yfjO* + *yfjO* add-back, and WT (Figure 6e). The Δ*yfjO* strain displayed a ∼20% reduction in OP-puro incorporation compared to WT and this defect was rescued by genetic *yfjO* add back (Figure 6f; Figure S22; Table S11). Overall, our results suggest that *yfjO* contributes to optimal cellular fitness and global protein translation capacity in *B. subtilis*.

## DISCUSSION

In this manuscript, we develop RNABPP-PS for proteomic profiling of bacterial RNA-modifying enzymes. Application of this method to *E. coli* and *B. subtilis* with 5-fluoropyrimidine nucleoside activity-based probes demonstrates its ability to reliably identify known and novel RNA pyrimidine modifying enzymes, and to illuminate catalytic mechanism. In particular, we identify the *B. subtilis* enzyme YfjO as a novel m^5^U writer that modifies U620 in the 23S rRNA. We therefore rename *yfjO* as *rlmS*. We also show the compatibility of 5-fluoropyrimidine nucleoside metabolic labeling with a diverse range of bacterial species, underscoring its potential application to systems with largely uncharacterized RNA modifications and RNA modifying enzymes.

Two m^5^U sites in 23S rRNA are conserved to varying degrees across bacteria. These modifications, m^5^U747 and m^5^U1939 (*E. coli* numbering) are installed by RlmC/RlmD homologues (51) – in *B. subtilis* a single enzyme, RlmCD (YefA) (63), installs both modification sites. Here we show that *B. subtilis* contains a third m^5^U site in the 23S rRNA at U620 (U576 in *E. coli*) installed by the methyltransferase RlmS (formerly YfjO). Although analysis of cryoEM structures between *E. coli* and *B. subtilis* ribosomes (74, 75) does not show a clear structural effect of the methylation, we demonstrate that *B. subtilis* Δ*rlmS* mutant strains show reduced protein translation efficiency and fitness defects compared to WT.

While rRNA modifications have been well-characterized in *E. coli*, they remain less studied in *B. subtilis*. The dispensable nature of individual rRNA modifications has made it challenging to directly connect a specific modification to a defined functional defect (18, 77). The prevailing view is that rRNA modifications fine-tune the structure and function of the ribosome. To date, no single rRNA modification has been shown to be essential for bacterial viability (18, 77), although combined deletions of multiple rRNA modifying enzymes can lead to substantial growth defects and impairment in ribosome assembly (18, 78). Consistent with this, most single-gene knockouts of rRNA modifying enzymes display either mild or undetectable growth phenotypes, although some exhibit reduced competitive fitness during long-term growth (18, 77, 79, 80). A notable exception is RlmE, which installs Um2552, an essential base in the A-site tRNA binding region in the peptidyl transferase center (PTC) (81). Loss of RlmE results in severe growth defects and delayed 50S assembly (17).

The newly identified RlmS substrate site, m^5^U620 (U576 according to *E. coli* numbering), is located in Helix 25a in Domain 0 of the 23S rRNA (Figure 4a). According to the revised domain model, Domain 0 forms the structural core of the large subunit (82). It is composed of six helices that are tightly interconnected with one another yet relatively insulated from the surrounding rRNA. Domain 0 appears to play a primarily structural role, forming a cleft that encloses the A-and P- sites of the peptidyl transferase center (PTC) and holds them in proximity (82). Proximity of Domain 0 to the PTC, combined with the observed protein translation rate defects in *rlmS* knockout cells, suggests a role for m^5^U620 modification in optimizing ribosome translocation, although we cannot rule out effects on ribosome stability and assembly.

Analysis of 5-FUrd-reactive proteins in *E. coli* and *B. subtilis* identified MnmG (GidA). Previous studies of *E. coli* MnmE/MnmG have speculated on the formation of a covalent complex during catalysis (58, 60), but the existence of this species has not been confirmed. Interestingly, Bandarian and co-workers identified the formation of an MnmE/MnmG-tRNA complex resulting from an *in vitro* enzymatic reaction that migrated on urea-PAGE and agarose gel, but ultimately concluded that this complex was non-covalent (59). Importantly, in their hands mutation of either Cys47 or Cys277 in MnmG to Ala did not abrogate complex formation. We demonstrate that MnmG forms an RNA-protein complex that can be enriched by aqueous-organic extraction and that is resistant to denaturing PAGE. We propose that this complex is distinct from that identified previously as it is formed in cells (rather than *in vitro*), migrates by SDS-PAGE with apparent molecular weight corresponding to a single MnmG-tRNA conjugate, and requires Cys47 and Cys277. The formation of this complex can be observed during catalysis with native RNA substrates and is stabilized by 5-FUrd modification. Our data argues for activation of the uridine substrate for methylation through a covalent RNA-protein adduct, likely involving a nucleophilic Cys residue in MnmG. We also find broad enrichment of bacterial PUS enzymes in our 5-FUrd RNABPP-PS experiment. Our previous RNABPP studies in human cells with 5-FUrd did not capture human PUS enzymes and whether 5-FUrd forms a covalent mechanism-based adduct with PUS enzymes has been controversial (54–56). Although we cannot exclude the formation of covalent 5-FUrd-PUS adducts, it is also possible that high-affinity, but non-covalent RNA-PUS adducts induced by 5-FUrd modification can be reliably enriched by aqueous-organic extraction. Capture of these species may be mediated by the relatively high frequency of metabolic 5-FUrd incorporation in bacteria as compared to mammalian cells.

Finally, the efficiency of metabolic RNA labeling with 5-fluoropyrimidines in bacteria suggests that our strategy is a promising approach for RNA modifying enzyme discovery and characterization in prokaryotes. Application of RNABPP-PS in additional bacterial species and with an expanded set of modified nucleosides should provide new insights into RNA modifying enzyme activity and enzymatic mechanism. These studies are currently underway in our group and will be reported in due course.

## MATERIALS AND METHODS

### Chemicals

5-FCyd and 5-FUrd were purchased from Carbosynth. All other chemicals were purchased from Sigma-Aldrich or Fisher Scientific unless otherwise specified.

### Bacterial strains

Bacterial strains used in this study are listed in Table S1. *E. coli* K12 BW25113 (83) was used as the *E. coli* WT reference strain. *E. coli* knockout strains, Δ*cdd*, Δ*codA*, Δ*trmA* and Δ*mnmG* were purchased from the *E. coli* Keio Knockout Collection (84). The *E. coli* Δ*cdd*Δ*codA* strain was generated as described below. *B. subtilis* WT168 was used as the *B. subtilis* WT reference strain (71). *B. subtilis* knockout strains from the Gross laboratory collection (70), Δ*yfjO::kan (*BKK08020) and Δ*yfjO::erm (*BKE08020), were obtained from the *Bacillus* Genetic Stock Center. Newly reported *B. subtilis* knockout (Δ*yfjO*) and add-back (Δ*yfjO+yfjO^+^*) strains were generated as described below. *V. harveyi* BB120 (85) was a gift from Bonnie L. Bassler. *K. aerogenes* ATCC 13048, *P. putida* ATCC 12633, *S. mutans* ATCC 25175, *S. epidermidis* ATCC 14990 and *M. smegmatis* ATCC 14468 were purchased from ATCC.

### Culture and medium

Bacterial strains were streaked from -80 ^°^C glycerol stocks onto appropriate agar medium and incubated for the indicated time. The following day, liquid cultures were inoculated from a single colony. Antibiotics for *E. coli* selection were used at the following concentration: 50 µg/µL kanamycin and 25 µg/µL chloramphenicol. Antibiotics for *B. subtilis* selection were used as follows: 7 µg/µL kanamycin, 5 µg/µL chloramphenicol and 1 µg/µL erythromycin. All media used in this study were purchased from Fisher Scientific unless otherwise indicated. Media powders were dissolved in milliQ water and sterilized by autoclaving before use. All bacteria were cultured at the specified temperature with shaking at 200 rpm. *E. coli* strains were cultured in Lysogeny Broth (LB) medium (Miller) at 37 °C for 18 hours. *B. subtilis* strains were cultured in LB medium (Lennox, Sigma-Aldrich) at 37 °C for 18 hours. *V. harveyi* BB120 strain was cultured in LM medium (LB medium (Miller) + 10 g/L NaCl) at 30 °C for 24 hours. *K. aerogenes* ATCC 13048 and *S. epidermidis* ATCC 14990 were cultured in Difco^TM^ Tryptic Soy Broth (TSB) at 37°C for 18 hours. *P. putida* ATCC 12633 was grown in Difco^TM^ Nutrient Broth (NB) at 30 °C for 24 hours. *S. mutans* ATCC 25175 was cultured in Oxoid^TM^ Brain Heart Infusion (BHI, Thermo Scientific) medium at 37 °C for 18 hours. *M. smegmatis* ATCC 14468 was cultured in Middlebrook 7H9 medium supplemented with ADC enrichment [Difco^TM^ Middlebrook 7H9 broth, 2.5 mL/L Tween-80, ADC comprising: 5 g/L bovine serum albumin (Gold Biotechnology), 2 g/L dextrose, 3 mg/L catalase (bovine, Sigma-Aldrich)] at 37 °C for 24 hours.

### Plasmids

For expression of TrmA and MnmG in *E. coli*, *trmA* and *mnmG* coding sequences were PCR amplified from *E. coli* K12 BW25113 genomic DNA. Site-directed mutagenesis of Cys47/Cys277 to Ser variants was generated by overlap extension PCR. PCR products were digested with BamHI (NEB) and NotI (NEB) and ligated into the pCA24N backbone (Addgene, plasmid #87741) using T4 DNA ligase (NEB) with an N-terminal 6xHis under isopropyl β-D-1-thiogalactopyranoside (IPTG)-inducible expression. For expression of YfjO in *E. coli*, *yfjO* was PCR amplified from *B. subtilis* WT 168 genomic DNA, digested with EcoRI (NEB) and XhoI (NEB), and cloned into a modified pET28a vector using T4 DNA ligase (NEB). For construction of *B. subtilis* add-back (Δ*yfjO+yfjO^+^*) strain, *yfjO* was cloned into integrative plasmid pBS1C (72) (Addgene, plasmid #55168) with two recombination sites for insertion at the *amyE* locus. The *yfjO* coding sequence along with 250 bp flanking region (5’ and 3’) was PCR amplified from *B. subtilis* WT 168 genomic DNA, digested with EcoRI (NEB) and PstI (NEB), and ligated into the pBS1C backbone with T4 DNA ligase (NEB).

### *E. coli* strain construction

To generate the *E. coli* double knockout strain Δ*cdd*Δ*codA*, the Δ*cdd::kan* locus from the Δ*cdd* strain was transferred into the clean WT *E. coli* BW25113 strain by P1 transduction, followed by removal of the kanamycin resistance cassette using FLP recombinase (83). The Δ*codA::kan* locus from the Δ*codA* strain was then transferred into the resulting Δ*cdd* strain by P1 transduction, and the kanamycin cassette was removed to yield the final double knockout strain Δ*cdd*Δ*codA*.

### Generation of *B. subtilis* knockout strain

The *yfjO* knockout strain (Δ*yfjO*) was generated using a double-crossover homologous recombination strategy (70). A linear DNA fragment targeting the *yfjO* gene (1 kb 5’ and 1kb 3’ flanking sequences of the *yfjO* locus and kanamycin resistance cassette) was amplified from the Δ*yfjO::kan (*BKK08020) genome using primers P.01 and P.02 (Table S12). The purified PCR product was transformed into *B. subtilis* WT 168 using natural competence (70). Briefly, 2 µg of DNA was added to 300 µL *B. subtilis* WT 168 competent cells and incubated for 2 hours at 37 °C, followed by plating on LB agar supplemented with kanamycin. After overnight incubation, six single colonies were restreaked for purification on a new selection plate and screened for insertion/deletion at the *yfjO* locus by PCR using primers P.03 and P.04 (Table S12). Two clones were further validated by Nanopore whole-genome sequencing (Plasmidsaurus). Sequencing reads were aligned to the *B. subtilis* reference genome (GeneBank assembly GCA_000009045.1). Verified clones were stored in 15% glycerol at -80 °C.

### Generation of *B. subtilis* add-back strain

The add-back strain (Δ*yfjO+yfjO^+^*) was constructed by integration of *yfjO* at the *amyE* locus. The pBS1C-*yfjO* plasmid was linearized with ScaI (NEB) and transformed into *B. subtilis* Δ*yfjO* cells using natural competence (70). Briefly, 1.5 µg of linearized plasmid was added to 300 µL *B. subtilis* Δ*yfjO* competent cells and incubated for 2 hours at 37 °C. The transformation mix was then plated on LB agar supplemented with kanamycin and chloramphenicol and incubated overnight. After ∼18 hours, six single colonies were purified by restreaking on new selection plate and screened for insertion at *amyE* locus by PCR using primers P.05 and P.06 (Table S12). Two confirmed clones were stored in 15% glycerol stock at -80 °C.

### Bacterial growth curves

For toxicity assays with 5-FCyd and 5FUrd, cultures were inoculated and grown in the appropriate medium at the indicated temperature. Upon reaching logarithmic phase, cultures were diluted to optical density (OD) = 0.1 in fresh medium and grown in 96-well plates until reaching mid-logarithmic phase (OD = 0.4). Cultures were then treated with increasing concentrations of 5-FCyd or 5FUrd (1 µM to 500 µM). OD600 measurements were recorded every 15 minutes on a Spectramax iD5 plate reader (Molecular Devices) until early stationary phase. For *B. subtilis* growth curves, cultures were inoculated into LB and grown at 37 °C. Logarithmic phase bacterial cultures were diluted to OD = 0.005 with fresh medium and transferred to 96-well plates for growth at 37 °C, 30 °C. or 25 °C. Cultures were monitored by OD600 measurement every 15 minutes using an Envision plate reader (PerkinElmer) until reaching stationary phase (24 or 30 hours). Two independent biological replicates with three technical replicates each were analyzed.

### Protein expression and purification

For protein purification, proteins were expressed in *E. coli* strain BL21 (DE3). TrmA-6xHis and YfjO-6xHis proteins were expressed overnight at 18 °C with 1mM IPTG. Cells were lysed by sonication, and TrmA-6xHis and YfjO-6xHis were purified using Ni-NTA affinity resin (Thermo Fisher) according to the manufacturer’s recommendations. The resulting proteins were concentrated to 1 mg/ml with an Amicon centrifugal filter (10,000 MWCO, Millipore). For *in cellulo* crosslinking experiments, TrmA-6xHis was expressed in Δ*trmA E. coli* for 6 hours at 37°C with 1mM IPTG. pCA24N-MnmG(WT)-6xHis, pCA24N-MnmG(C47S)-6xHis and pCA24N-MnmG(C277S)-6xHis were expressed in Δ*mnmG E. coli* for 6 hours at 37 °C with 1mM IPTG. For RNA modification analysis in *E. coli*, YfjO-6xHis was expressed in *E. coli* strain BL21 (DE3) for 6 hours at 37 °C with 1mM IPTG.

### RNA extraction and fractionation

For measurement of the metabolic incorporation of 5-fluoropyrimidines, bacterial cultures were treated with 5-FCyd or 5-FUrd at mid-logarithmic phase (OD = 0.5) and incubated until stationary phase (OD ∼2). Cells were harvested and lysed by cryomilling, and total RNA was isolated using TRIzol reagent (Thermo Fisher) following the manufacturer’s instructions. For measurement of native RNA modification in *B. subtilis*, cultures of WT, Δ*yfjO*, and Δ*yfjO+yfjO^+^* strains were grown at 37 °C for 8 hours until stationary phase (OD ∼2). Cells were harvested, resuspended in 50 mM ethylenediaminetetraacetic acid (EDTA), and treated with 2 mg/mL lysozyme (Themo Scientific) for 45 minutes in 37 °C. Cells were pelleted by centrifugation (15 min, 4,000 x g, 4 °C), and total RNA was isolated using TRIzol. RNA was fractionated into small (<200 nt) and large (>200 nt) RNA using the RNA Clean & Concentrator-25 (Zymo Research) following the manufacturer’s protocol. For rRNA separation, 16S and 23S rRNA were isolated by agarose gel purification. Gel slices containing rRNA were excised, mixed with three volumes of RNA agarose dissolving buffer (RAD, Zymo Research), incubated for 5 minutes at 50 °C, and transferred to RNA Clean & Concentrator-5 (Zymo Research) for purification according to the manufacturer’s recommendations.

### Nucleoside LC-QQQ-MS

RNA was digested and dephosphorylated to nucleosides followed by quantification via LC-QQQ-MS following literature precedent (32). Briefly, 1-2 µg of RNA was digested in a 30 µL reaction containing 2 U of nuclease P1 (Wako) at 37 °C for 2 hours (buffer: 7 mM sodium acetate, pH 5.2, 0.4 mM ZnCl_2_). Next, dephosphorylation was performed by adding 1.5 µL (7.5 U) Antarctic phosphatase (NEB) and 3.5 of 10x Antarctic phosphatase buffer, followed by incubation for 2 hours at 37 °C. Nucleosides were analyzed in positive ion dynamic multiple reaction monitoring (DMRM) mode using an Agilent 1290 Infinity II HPLC coupled to an Agilent 6495 triple quadrupole mass spectrometer. Chromatography was performed on a Hypersil GOLD aQ column (Thermo Scientific, 5 μm, 150 x 2.1 mm) at a flow rate of 0.4 mL/min using the following gradient (buffer A = 0.1% formic acid in water, buffer B = 0.1% formic acid in acetonitrile): start and hold at 0% B for 6 min, then increase to 1% B over 1.65 min, increase to 6% B over 1.7 min, hold at 6% for 0.75 min, increase to 50% B over 2 min, increase to 75% B over 2 min, hold at 75% B for 3 min. The following source parameters were used for the mass spectrometry: gas temperature 130 °C, gas flow 11 L/min, nebulizer pressure 25 psi, sheath gas temperature 400 °C, sheath gas flow 12 L/min, and capillary voltage 3.5 kV. Cell accelerator voltage (CAV) was set to 5 V. Transitions, fragmentor voltage and collision energy were optimized for each nucleoside (Table S2). MS1 (parent ion) to MS2 (deribosylated base ion) transition and retention time for canonical and modified nucleoside were set as described in Table S2. Standard curves were generated using commercially available ribonucleosides (Figure S3 and S4). Levels of 5-FCyd, 5-FUrd, m^5^C, m^5^U, and dihydrouridine (DHU) were determined by normalizing the concentration of modified nucleosides to the concentration of the corresponding canonical nucleosides in the sample. Data processing was performed using Agilent MassHunter Workstation Data Acquisition version 10.0. Three independent biological replicates were prepared for each sample.

### *In vitro* crosslinking assay

Δ*trmA E. coli* cultures were treated with 10 µM 5-FUrd (or left untreated) at mid-logarithmic phase (OD = 0.5) and incubated at 37 °C until stationary phase (OD ∼2, 6 hours). Cells were harvested and lysed by cryomilling, and total RNA was isolated using TRIzol reagent. Large and small RNA fractions were obtained using RNA Clean & Concentrator-25 (Zymo Reasearch). *In vitro* crosslinking reaction was performed as described previously with minor modifications (38). Briefly, isolated RNA was refolded by heating at 65 °C for 5 minutes followed by slow cooling to 25 °C at 0.1 °C/sec in RNase-free water. Refolded RNA (9 µM) was incubated with 3 µM purified TrmA protein in 20 µL reaction buffer (20 mM Tris-HCl pH 7.5, 100 mM NaCl, 5 mM MgCl_2_, 2 mM dithiothreitol (DTT), 0.4 U/ µL RNase inhibitor (Murine, NEB), and 100 mM SAM (NEB)) at 37 °C for 1 hour. For RNA digested samples, 1 µL RNase A/T1 mix (Thermo Fisher) was added and reactions were incubated for an additional 1 hour at 37 °C. The crosslinking reaction were analyzed using 10% sodium dodecyl sulfate-polyacrylamide gel electrophoresis (SDS-PAGE) followed by Coomassie staining.

### RNABPP-PS

RNABPP-PS samples were prepared from *E. coli* ΔcddΔcodA and *B. subtilis* WT 168 cultures treated with 50 µM 5-FCyd, 10 µM 5-FUrd, or left untreated at mid-logarithmic phase (OD = 0.5). Cultures (30 mL) were incubated at 37 °C for 6 hours until reaching stationary phase (OD ∼2). Cells were harvested, lysed by cryomilling, and the resulting powder was further lysed using glass beads (0.1 mm, Benchmark) in 600 µL TRIzol reagent (Thermo Fisher, 15596018). Phase separation-based extraction was performed following literature precedent with slight modifications (34, 35). Lysate were fully solubilized in 3 mL TRIzol reagent and centrifuged (5 min, 12,000 x g, 4 °C) to remove insoluble material. 600 µL chloroform was added and samples were vortexed and centrifuged based on the manufacturer’s protocol. After the first centrifugation, the interphase was isolated by discarding the aqueous upper phase and the organic lower phase. The remaining interphase was then resuspended in 1 mL of TRIzol for a second isolation step and repeated for three total interphase isolations. After the third extraction, the interphase was precipitated using 900 µL of methanol and resuspended in 100 µL of 100 mM triethylammonium bicarbonate buffer (TEAB), pH 8.5, containing 1 mM MgCl_2_, and 1% SDS. Samples were digested with 2 µL RNase A/T1 mix for 4 hours at 37 °C, followed by addition of 2 µL RNase Cocktail and overnight incubation to ensure complete RNA digestion. Digested interphase was subjected to 1 mL TRIzol and 200 µL chloroform and the organic layer was collected and precipitated with nine volumes of methanol. Proteins pellets were resuspended in 100 µL of 10 mM 4-(2-hydroxyethyl)-1-piperazineethanesulfonic acid (HEPES) buffer, pH 7.5. Protein concentration was measured by bicinchoninic acid (BCA) assay (Thermo Scientific) and adjusted to a final concentration of 1.0 µg/µL. Three independent biological replicates were prepared for each sample.

### Label-free proteomics sample preparation

Protein samples at a final concentration of 1.0 µg/µL were diluted 1:1 with 100 mM TEAB (pH 8.5), reduced with 5 mM Tris(2-carboxyethyl)phosphine (TCEP, Pierce) for 1 hour at 55 °C and alkylated with 15 mM of iodoacetamide for 30 minutes at ambient temperature in the dark. Samples were digested overnight at 37 °C with MS-grade trypsin gold (Promega) at a ratio of 1 µg enzyme per 50 µg protein. Reactions were quenched by the addition of Optima-grade formic acid to a final concentration of 1%. Peptides were desalted using C18 Stage-Tips (86), eluted in 70% acetonitrile / 0.1% formic acid, and dried to completion. Dried peptides were resuspended in 0.1% formic acid in water for LC-MS/MS analysis.

### Proteomics LC-MS/MS

Peptides were loaded (∼200 ng per injection) onto a trap cartridge (C18 Pepmap, 5 µM particle size, 5 mm X 300 µM) and separated on an analytical column (PepSep C18, 1.5 µm, 10 cm x 150 µm) at 50 °C with a Bruker nanoElute 2 LC system coupled with a Bruker timsTOF fleX mass spectrometer. Peptides were eluted at a flow rate of 500 nL/min using the following gradient (buffer A = 0.1% formic acid in water, buffer B = 0.1% in acetonitrile): start at 4% B, then increase to 35% B over 20 min, increase to 95% B over 0.5 min, hold at 95% for 3.9 min). Data were acquired in positive ion and data-independent acquisition (dia-PASEF) mode over a *m/z* range of 100-1700 with a cycle time of 0.85 s. MS parameters were set as follow: number of MS1 ramps 1, number of MS/MS ramps 7, number of MS/MS windows 25, and collision energy 20-50 eV. Ion mobility parameters were set to 0.7-1.4 V•s/cm2, 100 ms ramp time, 100 ms accu. time, 100% duty cycle, and 9.42 Hz ramp rate.

### Proteomics data analysis

Data-independent acquisition was processed via DIANN 1.8.1 (87). Parameters were set as follows: trypsin/P digestion, three missed cleavages, and up to three variable modifications (N-term M excision, Ox(M), Ac(N-term) and C carbamidomethylation). Peptide length range was 7-30, precursor charge range 1-4, *m/z* range 300-1800, and fragment ion range 200-1800. Mass accuracy and MS1 accuracy were both set to 10. The following settings were enabled: “Use isotopologues”, “MBR”, “No shared spectra”, “Heuristic protein inference”. The precursor FDR threshold was set to 1%. Spectral libraries were generated *in silico* via DIANN from all known *E. coli* and *B. subtilis* proteins database downloaded from UniProt (*E. coli* proteome: UP000000625_2024_07_25; and *B. subtilis* proteome: UP000001570_2024_06_14). The resulting matrix.pg file was imported into Perseus (2.0.11). Data were transformed (log base 2) and annotated as either “5-FCyd”, “5-FUrd”, or “control”. Missing values were imputed from a normal distribution and normalization via median subtraction was applied. Differential protein abundance was assessed using a two-sample t-test for statistical significance, and volcano plots were generated in GraphPad Prism 10.

### *In cellulo* crosslinking assay

Δ*trmA E. coli* strain expressing TrmA and Δ*mnmG* expressing MnmG(WT)-6xHis, MnmG(C47S) -6xHis, or MnmG(C277S)-6xHis, were incubated with either 10 µM or 50 µM 5-FUrd at mid-logarithmic phase (OD = 0.5) for 6 hours or left untreated. Cells were harvested, lysed by cryomilling, and processed by three sequential interphase extractions as described above. Interphase fractions were resuspended in 100 mM TEAB buffer containing 1% SDS and analyzed directly by SDS-PAGE or subjected to RNase digestion for 2 hours at 37 °C prior analysis. Samples were resolved on 10% SDS-PAGE followed by Western blot using an anti-His antibody (1:2500, GenScript).

### *In vitro* methylation reaction

*In vitro* methylation assays were performed following a published protocol with minor modifications (79). Purified rRNAs were refolded by heating at 65 °C for 5 minutes followed by slow cooling to 25 °C at 0.1 °C/sec in RNase-free water. 200 µg of refolded rRNA was incubated with 5 µg (∼100 pmol) of purified YfjO protein in 100 µL reaction buffer (200 mM Tris-HCl, pH 7.5, NH_4_Cl 80 mM, 4 mM Mg(OAc)_2_, 12 mM DTT, 1 U/ µL RNase inhibitor (Murine, NEB), and 50 µM SAM (NEB)) for 2 hour at 37 ^°^C. Reactions were quenched and purified using phenol/chloroform extraction followed by ethanol precipitation. Precipitated rRNA were reconstituted in RNase-free water and analyzed by double-cleavage oligo LC-MS.

### Double-cleavage RNase H assay

The double-cleavage assay was performed following literature precedent with slight modifications (73). Briefly, rRNA purified from *B. subtilis* or *E. coli* cultures, or after *in vitro* methylation reactions, were annealed with a 2’-deoxy-2’-O-methyl chimeric oligonucleotide (Table S12) targeting U620 (*B. subtilis*) or U576 (*E. coli*). 500 pmol of 2’-deoxy-2’-O-methyl chimeric oligonucleotide was annealed with 200 µg rRNA by heating at 95 °C for 2 minutes and slow cooling to 25 °C at 0.1 °C/sec in 100 µL annealing buffer (10 mM Tris-HCl, pH 7.7, 50 mM NaCl, 1 mM EDTA). Cleavage reactions were initiated by adding 4 µL of RNase H (20 U, NEB), 13 µL 100 mM DTT, and 13 µL homemade 10x RNase H reaction buffer (450 mM Tris-HCl, pH 7.7, 500 mM NaCl, 50 mM MgCl_2_), followed by incubation for 16 hours at 37 °C. Upon completion, salt and buffer were depleted using an Amicon centrifugal filter (10 kD MWCO, 0.5 mL, Millipore), and the resulting 5’-phosphate and 3’-hydroxyl oligonucleotide fragments were dried to completion. Dried oligonucleotide fragments were reconstituted in buffer A (10 mM hexafluoroisopropanol (HFIP) and 8.6 mM triethylamine (TEA) in water) and one third of each sample was injected for LC-MS/MS analysis.

### Oligonucleotide LC-MS/MS

Oligonucleotide fragments were analyzed in negative ion mode on an Agilent 1290 Infinity II HPLC coupled to an Agilent 6545XT Q-TOF mass spectrometer. Oligonucleotides were separated on a Poroshell 120 EC-C18 (Agilent, 1.9 µm, 2.1 x 50 mm) at a flow rate of 250 µL/minute using the following gradient: (buffer A = 10 mM HFIP and 8.6 mM TEA in water, buffer B = acetonitrile) start and hold at 0% B for 2 min, then increase to 20% B over 18 min, increase to 95% B over 2 min, and hold at 95% for 1 min. The ionization source working parameters were set as follows: capillary voltage 3.5 kV, gas temperature 275 °C, drying gas flow rate 12 L/min, nebulizer pressure 25 psi, the sheath gas temperature 350 °C and sheath gas flow 12 L/min. Data were acquired in a full scan from *m/z* 120-3200 at a fixed acquisition rate of one spectrum per second. Detected RNA fragments were compared against the theoretical RNase H digestion products and selected for collision-induced dissociation (CID) under 3 collision energy channels (35, 45, 55 V) for fragmentation. Extracted ion chromatograms (EICs), MS spectra, and MS/MS fragmentation spectra are reported. Manual annotation of MS/MS spectra was performed by calculation expected *m/z* value of oligonucleotide fragments using Mongo Oligo Mass Calculator (v2.07).

### Sporulation/germination efficiency assay

*B. subtilis* strains were grown and induced to sporulate following literature procedure (88). Briefly, overnight cultures were diluted to OD = 0.1 in CH medium (89) and grown at 37 °C until OD = 0.6. Sporulation was initiated by medium exchange to sporulation medium as described by Sterlini and Mandelstam (90), followed by incubation for 18 hours at 37 °C with shaking. Sporulated cultures were diluted to OD = 0.6 and split into two 250 µL aliquots. One aliquot remained untreated, while the second was subjected to heat treatment (40 min at 90 °C). Samples were serially diluted to 10^-5^, and 100 µL was plated on LB agar and incubated overnight at 30 °C. Sporulation/germination efficiency was quantified as the ratio of colony forming unit (CFU) following heat treatment relative to untreated control. Sporulation/germination (%) = (CFU heat-treated/CFU untreated) x 100.

### Competitive growth

WT and Δ*yfjO* strains were grown separately to mid-logarithmic phase (OD = 0.6), then each culture was inoculated 1:1,000 into fresh LB medium and combined at a 1:1 ratio to generate the starting co-culture (P_0_). Cocultures were incubated for 18 hours at 30 °C, 37 °C, or 42 °C, followed by serial passaging (1:1000 dilution, ∼10 generations of growth per passage). Genomic DNA was extracted at defined passages 7, 14, or 23 using Wizard genomic DNA isolation kit (Promega) following manufacturer’s protocol. Strain abundance was quantified by PCR amplification of the *yfjO* locus using primers P.03 and P.04 (Table S12). WT strain yielded a 1464 bp amplicon (upper band) band, whereas Δ*yfjO* strain produced a 1135 bp product (lower band) band due to the kanamycin resistance cassette. Band intensities were calculated by quantification of PCR products densitometry using ImageJ, and relative strain abundance was calculated as the percentage of total signal from both PCR products. Calibration samples with defined WT:Δ*yfjO* ratios (0:100 to 100:0) were processed in parallel. Three independent biological replicates were analyzed. Fitness = (strain% at defined passage */* strain% at P_0_)^(1/t) where t = number of passage x generation per passage) (91). Relative fitness = fitness Δ*yfjO/* fitness WT.

### Global protein translation assay

*B. subtilis* cultures at mid-logarithmic phase (OD = 0.6) were treated with 13 µm OP-Puro (MedChemExpress) for 30 minutes at 37 °C with shaking. Cells were harvested, resuspended in phosphate-buffered saline (PBS), and incubated with 2 mg/mL lysozyme (Themo Scientific) for 20 minutes at 37 °C. Cells were pelleted by centrifugation (2 min, 5,000 x g, 4 °C), resuspended in lysis buffer (1% SDS in PBS and cOmplete™ EDTA-free Protease Inhibitor Cocktail) and lysed by sonication. Approximately 100 µg protein lysate was diluted to 80 µL in PBS. Click labeling was performed by adding 10 µL of 10 x CuAAC reagent mix (10 mM CuSO_4_•5H_2_O, 20 mM Tris hydroxypropyltriazolylmethylamine (THPTA), 1 mM Cy5-azide (Lumiprobe)), followed by addition of 10 µL of 100 mM sodium ascorbate (freshly prepared). Reactions were incubated for 40 minutes at 37 °C with shaking (550 rpm). Proteins were precipitated by addition of 1:4 ice-cold acetone, pelleted by centrifugation (5,000 x g, 20 minutes, 4 °C), and analyzed using 10% SDS-PAGE. Fluorescent signal (Cy5 channel) was acquired with a Typhoon scanner (Cytiva), followed by Coomassie staining. Controls included no OP-Puro treatment, pretreatment with puromycin (Pre Puro), pretreatment with chloramphenicol (Pre Cm), and no click. Densitometry of fluorescent and Coomassie bands was performed using ImageJ. Cy5 signal was normalized to protein loading, and values were expressed relative to WT replicate 1. Median ± s.e.m. values from six independent biological replicates are reported.

## ACKNOWLEDGMENTS

We thank John Eng and Venu Vendavasi at the Princeton Chemistry Department Proteomics and Mass Spectrometry Center for assistance with RNA and protein mass spectrometry. R.E.K. acknowledges support from the NSF (MCB-1942565 and MCB-2448119). Q.Z.J. acknowledges support from an EMBO Postdoctoral Fellowship (ALTF 177-2024), the Fulbright Program, and the Israel Council for Higher Education (VATAT). N.J.Y. was supported by a generous gift from the Edward C. Taylor 3rd Year Graduate Fellowship in Chemistry.

