## Supporting Information for "Activity-based profiling of bacterial RNA-modifying enzymes reveals species-specific 5-methyluridine modification in *Bacillus subtilis* 23S rRNA"

Table of Contents

i. Supplementary Tables S1-S12

ii. Supplementary Figures S1-S22

**Table S1.** Bacterial strains information. Kanamycin (kan), chloramphenicol (Cm), and erythromycin (erm).

| # | Strain name | Genotype | Source |
|---|---|---|---|
| 1 | *Escherichia coli* (*E. coli*) K-12 BW25113 | Wildtype (WT) | Ya-Ming Hou lab (1) |
| 2 | *E. coli ΔcddΔcodA* | BW25113 *ΔcddΔcodA* | This study |
| 3 | *E. coli Δcdd* JW2131-1 | BW25113 *Δcdd::kan$^r$* | *E. coli* Keio Knockout Collection (2) |
| 4 | *E. coli ΔcodA* JW0328-1 | BW25113 *ΔcodA::kan$^r$* | *E. coli* Keio Knockout Collection (2) |
| 5 | *E. coli ΔtrmA* JW3937-1 | BW25113 *ΔtrmA::kan$^r$* | *E. coli* Keio Knockout Collection (2) |
| 6 | *E. coli ΔmnmG* JW3719-1 | BW25113 *ΔGidA::kan$^r$* | *E. coli* Keio Knockout Collection (2) |
| 7 | *E. coli* BL21 (DE3) | $F^-$*ompT hsdS*$_B$ ($r_B^-$, $m_B^-$) *gal dcm* (DE3) | |
| 8 | *Bacillus subtilis* (*B. subtilis*) WT168 | WT, *trpC2* | *Bacillus* Genetic Stock Center (BGSC) (3) |
| 9 | *B. subtilis ΔyfjO::kan* BKK08020 | *trpC2 ΔyfjO::kan$^r$* | BGSC (4) |
| 10 | *B. subtilis ΔyfjO::erm* BKE08020 | *trpC2 ΔyfjO::erm$^r$* | BGSC (4) |
| 11 | *B. subtilis ΔyfjO* | *trpC2 ΔyfjO::kan$^r$* | This study |
| 12 | *B. subtilis ΔyfjO+yfjO$^+$* | *trpC2 ΔyfjO::kan$^r$ amyE::(yfjO$^+$ Cm$^r$)* | This study |
| 13 | *Klebsiella aerogenes* (*K. aerogenes*) ATCC 13048™ | | ATCC |
| 14 | *Vibrio harveyi* (*V. harveyi*) BB120 | | Bonnie L. Bassler lab (5) |
| 15 | *Pseudomonas putida* (*P. putida*) ATCC 12633™ | | ATCC |
| 16 | *Streptococcus mutans* (*S. mutans*) ATCC 25175™ | | ATCC |
| 17 | *Staphylococcus epidermidis* (*S. epidermidis*) ATCC 14990™ | | ATCC |
| 18 | *Mycobacterium smegmatis* (*M. smegmatis*) ATCC 14468™ | | ATCC |

**Table S2.** LC-QQQ-MS parameters used for measurement the concentration of modified and canonical nucleosides in RNA. Nucleosides were detected in positive ion dynamic multiple reaction monitoring (DMRM) mode. Cell accelerator voltage (CAV) was set to 5 V for all transitions.

| Compound | Precursor ion (*m/z*) | Product ion (*m/z*) | Fragment or (V) | Collision energy (V) | Retention time (min) | Retention window (min) |
|---|---|---|---|---|---|---|
| **A** | 268.1 | 136 | 166 | 20 | 6.12 | 3 |
| **G** | 284.1 | 152 | 166 | 14 | 8.32 | 3 |
| **C** | 244.1 | 112 | 166 | 20 | 1.98 | 3 |
| **U** | 245.1 | 113 | 166 | 14 | 3.34 | 2.5 |
| **5-FCyd** | 262 | 130 | 166 | 14 | 3.02 | 3 |
| **5-FUrd** | 263 | 131 | 166 | 14 | 4.41 | 2 |
| **DHU** | 247.1 | 115 | 166 | 14 | 1.75 | 3 |
| **$m^5C$** | 258.1 | 126 | 166 | 14 | 3.38 | 3 |
| **$m^5U$** | 259.1 | 127 | 166 | 14 | 7.28 | 3 |

**Table S3.** Total RNA labeling with 5-FUrd across a panel of bacteria. Nucleoside concentrations in total RNA were measured using LC-QQQ-MS after metabolic labeling with 5-FUrd. Nucleoside concentrations are reported as ng/mL. Levels of 5-FCyd or 5-FUrd are indicated as a percentage of total Cyd or Urd, respectively. R, replicate; Con., concentration; N.D., not detected.

| Bacterial strain | 5-FUrd Con. (μM) | A | G | C | U | 5-FCyd | 5-FUrd | 5-FCyd/C (%) | 5-FUrd/U (%) |
|---|---|---|---|---|---|---|---|---|---|
| *E. coli ΔcddΔcodA* R1 | 10 | 352.8 | 605.5 | 319.5 | 268.8 | N.D. | 9.3 | N.D. | 2.9 |
| *E. coli ΔcddΔcodA* R2 | 10 | 439.8 | 750.2 | 392.2 | 328.1 | N.D. | 13.6 | N.D. | 3.5 |
| *E. coli ΔcddΔcodA* R3 | 10 | 472.2 | 810.3 | 430.3 | 354.5 | N.D. | 14.7 | N.D. | 3.4 |
| *K. aerogenes* R1 | 50 | 390.0 | 943.3 | 571.8 | 308.2 | N.D. | 38.7 | N.D. | 12.5 |
| *K. aerogenes* R2 | 50 | 383.4 | 932.0 | 558.6 | 298.4 | N.D. | 42.2 | N.D. | 14.1 |
| *K. aerogenes* R3 | 50 | 377.4 | 921.2 | 551.2 | 299.4 | N.D. | 38.0 | N.D. | 12.7 |
| *V. harveyi* R1 | 100 | 415.1 | 582.2 | 349.9 | 341.8 | 0.17 | 79.9 | 0.05 | 23.4 |
| *V. harveyi* R2 | 100 | 384.3 | 545.3 | 331.9 | 321.5 | 0.04 | 66.8 | 0.01 | 20.8 |
| *V. harveyi* R3 | 100 | 359.3 | 507.9 | 311.1 | 294.7 | 0.07 | 71.1 | 0.02 | 24.1 |
| *P. putida* R1 | 50 | 301.8 | 456.2 | 291.2 | 246.3 | N.D. | 51.7 | N.D. | 21.0 |
| *P. putida* R2 | 50 | 348.1 | 520.2 | 325.6 | 299.9 | N.D. | 48.5 | N.D. | 16.2 |
| *P. putida* R3 | 50 | 394.1 | 592.5 | 386.0 | 358.4 | N.D. | 37.7 | N.D. | 10.5 |
| *B. subtilis* R1 | 10 | 371.6 | 788.5 | 435.9 | 253.4 | 1.46 | 80.7 | 0.33 | 31.9 |
| *B. subtilis* R2 | 10 | 356.6 | 732.8 | 441.6 | 257.6 | 1.17 | 74.8 | 0.26 | 29.1 |
| *B. subtilis* R3 | 10 | 390.2 | 805.3 | 442.6 | 272.0 | 1.02 | 72.7 | 0.23 | 26.7 |
| *S. mutans* R1 | 25 | 401.2 | 583.3 | 359.7 | 293.3 | 0.11 | 116.5 | 0.03 | 39.7 |
| *S. mutans* R2 | 25 | 377.8 | 547.4 | 330.7 | 299.8 | 0.09 | 114.4 | 0.03 | 38.2 |
| *S. mutans* R3 | 25 | 337.9 | 486.3 | 300.6 | 270.7 | N.D. | 105.4 | N.D. | 38.9 |
| *S. epidermidis* R1 | 10 | 603.1 | 1010.2 | 412.0 | 381.3 | 0.37 | 41.7 | 0.09 | 10.9 |
| *S. epidermidis* R2 | 10 | 502.0 | 838.5 | 342.7 | 322.3 | 0.32 | 28.4 | 0.09 | 8.8 |
| *S. epidermidis* R3 | 10 | 472.6 | 781.4 | 321.0 | 300.8 | 0.07 | 26.4 | 0.02 | 8.7 |
| *M. smegmatis* R1 | 100 | 271.6 | 527.5 | 374.8 | 275.3 | N.D. | 19.4 | N.D. | 7.0 |
| *M. smegmatis* R2 | 100 | 247.6 | 488.7 | 358.9 | 257.8 | N.D. | 20.9 | N.D. | 8.1 |
| *M. smegmatis* R3 | 100 | 280.8 | 552.5 | 393.3 | 287.8 | N.D. | 26.3 | N.D. | 9.1 |

**Table S4.** Total RNA labeling with 5-FCyd across a panel of bacteria. Nucleoside concentrations in total RNA were measured using LC-QQQ-MS after metabolic labeling with 5-FCyd. Nucleoside concentrations are reported as ng/mL. Levels of 5-FCyd or 5-FUrd are indicated as a percentage of total Cyd or Urd, respectively. R, replicate; Con., concentration; N.D., not detected.

| Bacterial strain | 5-FCyd Con. (μM) | A | G | C | U | 5-FCyd | 5-FUrd | 5-FCyd/C (%) | 5-FUrd/U (%) |
|---|---|---|---|---|---|---|---|---|---|
| *E. coli ΔcddΔcodA* R1 | 50 | 449.6 | 764.7 | 388.8 | 333.2 | 13.8 | 1.0 | 3.5 | 0.3 |
| *E. coli ΔcddΔcodA* R2 | 50 | 472.1 | 820.9 | 430.4 | 348.9 | 14.6 | 0.7 | 3.4 | 0.2 |
| *E. coli ΔcddΔcodA* R3 | 50 | 501.8 | 884.3 | 466.0 | 372.1 | 16.5 | 0.6 | 3.5 | 0.1 |
| *K. aerogenes* R1 | 50 | 385.8 | 938.5 | 566.6 | 305.3 | 3.9 | 36.3 | 0.7 | 11.9 |
| *K. aerogenes* R2 | 50 | 348.3 | 843.6 | 503.3 | 271.4 | 3.3 | 32.2 | 0.6 | 11.8 |
| *K. aerogenes* R3 | 50 | 413.2 | 1003.8 | 604.1 | 331.8 | 4.3 | 38.3 | 0.7 | 11.5 |
| *V. harveyi* R1 | 100 | 452.7 | 634.0 | 395.2 | 390.1 | 0.17 | 83.5 | 0.04 | 21.4 |
| *V. harveyi* R2 | 100 | 387.5 | 547.4 | 329.6 | 311.5 | N.D. | 77.9 | N.D. | 25.0 |
| *V. harveyi* R3 | 100 | 445.2 | 627.5 | 381.6 | 367.4 | 0.18 | 88.3 | 0.05 | 24.0 |
| *P. putida* R1 | 50 | 349.4 | 522.8 | 355.8 | 308.5 | N.D. | 1.4 | N.D. | 0.5 |
| *P. putida* R2 | 50 | 256.8 | 373.0 | 238.0 | 239.3 | N.D. | 3.0 | N.D. | 1.2 |
| *P. putida* R3 | 50 | 376.3 | 574.0 | 389.9 | 357.7 | N.D. | 1.3 | N.D. | 0.3 |
| *B. subtilis* R1 | 10 | 394.8 | 816.2 | 417.1 | 285.8 | 38.5 | 52.2 | 9.2 | 18.2 |
| *B. subtilis* R2 | 10 | 417.6 | 902.1 | 470.6 | 317.3 | 33.8 | 35.2 | 7.1 | 11.1 |
| *B. subtilis* R3 | 10 | 444.7 | 950.3 | 505.4 | 340.6 | 35.3 | 41.4 | 6.9 | 12.1 |
| *S. mutans* R1 | 10 | 429.0 | 789.1 | 349.2 | 308.1 | 22.1 | 18.8 | 6.3 | 6.1 |
| *S. mutans* R2 | 10 | 257.7 | 469.5 | 202.7 | 179.7 | 12.7 | 13.0 | 6.3 | 7.2 |
| *S. mutans* R3 | 10 | 452.2 | 869.6 | 390.7 | 348.6 | 24.8 | 22.3 | 6.3 | 6.4 |
| *S. epidermidis* R1 | 10 | 441.5 | 729.8 | 297.5 | 280.8 | 5.5 | 24.3 | 1.8 | 8.6 |
| *S. epidermidis* R2 | 10 | 525.8 | 878.5 | 362.1 | 340.4 | 6.4 | 31.7 | 1.7 | 9.3 |
| *S. epidermidis* R3 | 10 | 473.8 | 797.3 | 330.9 | 300.7 | 7.0 | 33.9 | 2.1 | 11.2 |
| *M. smegmatis* R1 | 100 | 282.8 | 563.3 | 413.0 | 290.2 | N.D. | 7.7 | N.D. | 2.6 |
| *M. smegmatis* R2 | 100 | 279.3 | 553.0 | 388.8 | 285.7 | N.D. | 20.5 | N.D. | 7.2 |
| *M. smegmatis* R3 | 100 | 295.3 | 584.3 | 431.2 | 302.1 | N.D. | 8.5 | N.D. | 2.8 |

**Table S5.** Pyrimidine-specific RNA-modifying enzymes in *E. coli* and *B. subtilis* and their identification using 5-Fluoropyrimidnes-based RNABPP-PS. Substrate RNA for each enzyme is indicated. The number of enriched enzymes using RNABPP-PS / the total number of known enzymes is indicated for each enzyme family.

| ***E. coli*** | | | ***B. subtilis*** | | |
|---|---|---|---|---|---|
| **Known enzymes** | **Substrate RNA** | **Enrichment statues with RNABPP-PS** | **Known enzymes** | **Substrate RNA** | **Enrichment statues with RNABPP-PS** |
| **5-methylcytidine methyltransferases ($m^5C$ MTases)** | | | | | |
| RsmF (6) | 16S rRNA | Enriched | RsmB UniProt KB (P94464) | Putative 16S rRNA | Enriched |
| RsmB (6) | 16S rRNA | Enriched | | | |
| RlmI (6) | 23S rRNA | Enriched | | | |
| RNABPP-PS identified / known enzymes: 3/3 | | | RNABPP-PS identified / known enzymes: 1/1 | | |
| **5-methyluridine methyltransferases ($m^5U$ MTases)** | | | | | |
| RlmD (7) | 23S rRNA | Enriched | RlmCD (yefA) (8) | 23S rRNA | Enriched |
| RlmC (7) | 23S rRNA | Enriched | TrmFO (9) | tRNA | Enriched |
| TrmA (10) | tRNA | Enriched | | | |
| RNABPP-PS identified / known enzymes: 3/3 | | | RNABPP-PS identified / known enzymes: 2/2 | | |
| **Dihydrouridine synthases (DUS)** | | | | | |
| DusA (11) | tRNA | Enriched | Dus1 (12) | tRNA | Enriched |
| DusB (11) | tRNA | Low enrichment | Dus2 (12) | tRNA | Enriched |
| DusC (11) | tRNA | Enriched | | | |
| RNABPP-PS identified / known enzymes: 3/3 | | | RNABPP-PS identified / known enzymes: 2/2 | | |
| **Pseudouridine synthases (PUS)** | | | | | |
| TruA (13) | tRNA | Enriched | TruA (14) | tRNA | Enriched |
| TruB (13) | tRNA | Enriched | TruB (14) | tRNA | Enriched |
| TruC (13) | tRNA | N.D. | RluB (15) | 23S rRNA | Enriched |
| TruD (13) | tRNA | No enrichment | YlyB (14) | Putative 23S rRNA | Enriched |
| RsuA (13) | 16S rRNA | Low enrichment | YhcT (14) | 23S rRNA | Enriched |
| RluA (13) | 23S rRNA & tRNA | Enriched | YtzG UniProt KB (O32068) | Putative 16S rRNA | Enriched |
| RluB (13) | 23S rRNA | Enriched | YjbO (14) | Putative tRNA (5) | Enriched |
| RluC (13) | 23S rRNA | Enriched | | | |
| RluD (13) | 23S rRNA | Enriched | | | |
| RluE (13) | 23S rRNA | Enriched | | | |
| RluF (13) | 23S rRNA | Enriched | | | |
| RNABPP-PS identified / known enzymes: 9/11 | | | RNABPP-PS identified / known enzymes: 7/7 | | |

**Table S6.** The differences detected in *ΔyfjO* clone (#1) relative to WT consist of only mutations located exclusively in non-coding regions. Whole genome sequencing alignment of *B. subtilis* WT and newly reported *ΔyfjO* clone (#1). Nanopore sequencing reads were aligned to reference genome (GeneBank assembly GCA_000009045.1).

| # | Genome position | Type of mutation | Reference base | New base | Protein level | Gene annotation |
|---|---|---|---|---|---|---|
| 1 | 556,280 | Insertion +1 | - | C | - | - |
| 2 | 556,477 | Substitution | A | G | - | - |
| 3 | 556,478 | Substitution | A | G | - | - |
| 4 | 873,404-874,799 | Deletion -1395<br>Insertion +1066 | - | - | - | *ΔyfjO::kan$^r$* |
| 5 | 2,061,689 | Deletion -1 | T | - | - | - |
| 6 | 2,061,692 | Insertion +1 | - | G | - | - |
| 7 | 2,061,693 | Substitution | A | C | - | - |
| 8 | 2,061,699 | Substitution | T | C | - | - |
| 9 | 2,061,706 | Substitution | A | C | - | - |
| 10 | 2,061,710 | Substitution | A | G | - | - |

**Table S7.** $m^5C$ and $m^5U$ levels in different RNA species from *B. subtilis* WT, *ΔyfjO*, and *ΔyfjO+yfjO*$^+$ strains measured by nucleoside LC-QQQ-MS. Nucleoside concentrations are reported as ng/mL. Levels of $m^5C$, $m^5U$, and DHU are indicated as a percentage of total Cyd or Urd. R, replicate; N.D., not detected.

| | A | G | C | U | DHU | $m^5C$ | $m^5U$ | DHU /U (%) | $m^5C$ /C (%) | $m^5U$ /U (%) |
|---|---|---|---|---|---|---|---|---|---|---|
| **Total RNA** | | | | | | | | | | |
| WT R1 | 844.2 | 1453.1 | 991.0 | 749.5 | 10.64 | 0.49 | 8.44 | 1.42 | 0.05 | 1.13 |
| WT R2 | 908.7 | 1574.8 | 1065.5 | 785.5 | 11.23 | 0.49 | 8.77 | 1.43 | 0.05 | 1.12 |
| WT R3 | 742.4 | 1242.1 | 1079.7 | 764.9 | 10.18 | 0.46 | 8.05 | 1.33 | 0.04 | 1.05 |
| *ΔyfjO* R1 | 956.5 | 1634.4 | 1101.3 | 862.7 | 11.28 | 0.44 | 7.92 | 1.31 | 0.04 | 0.92 |
| *ΔyfjO* R2 | 890.5 | 1539.2 | 1039.2 | 797.4 | 11.86 | 0.43 | 8.22 | 1.49 | 0.04 | 1.03 |
| *ΔyfjO* R3 | 970.0 | 1694.0 | 1142.1 | 863.3 | 13.01 | 0.45 | 8.93 | 1.51 | 0.04 | 1.03 |
| **Small RNA** | | | | | | | | | | |
| WT R1 | 567.3 | 1280.2 | 730.3 | 451.6 | 34.66 | 0.75 | 27.96 | 7.68 | 0.10 | 6.19 |
| WT R2 | 642.3 | 1454.6 | 855.9 | 510.4 | 42.33 | 0.71 | 34.33 | 8.29 | 0.08 | 6.72 |
| WT R3 | 614.7 | 1329.0 | 764.9 | 480.8 | 34.78 | 0.73 | 28.46 | 7.23 | 0.10 | 5.92 |
| *ΔyfjO* R1 | 718.4 | 1447.9 | 820.7 | 548.8 | 30.62 | 0.80 | 30.00 | 5.58 | 0.10 | 5.47 |
| *ΔyfjO* R2 | 615.9 | 1390.7 | 817.4 | 489.7 | 40.53 | 0.73 | 33.00 | 8.28 | 0.09 | 6.74 |
| *ΔyfjO* R3 | 578.5 | 1298.7 | 767.3 | 458.8 | 39.03 | 0.74 | 31.14 | 8.51 | 0.10 | 6.79 |
| **16S rRNA** | | | | | | | | | | |
| WT R1 | 743.7 | 1301.3 | 624.6 | 535.2 | 0.73 | 1.48 | 0.86 | 0.14 | 0.24 | 0.16 |
| WT R2 | 542.2 | 947.5 | 450.0 | 390.9 | 0.66 | 1.26 | 0.76 | 0.17 | 0.28 | 0.20 |
| WT R3 | 633.6 | 1130.3 | 525.9 | 459.8 | 0.67 | 1.39 | 0.85 | 0.15 | 0.26 | 0.18 |
| *ΔyfjO* R1 | 546.2 | 969.3 | 451.3 | 390.9 | 0.65 | 1.22 | 0.71 | 0.17 | 0.27 | 0.18 |
| *ΔyfjO* R2 | 605.7 | 1106.3 | 506.1 | 439.9 | 0.68 | 1.35 | 0.69 | 0.15 | 0.27 | 0.16 |
| *ΔyfjO* R3 | 444.3 | 805.9 | 365.9 | 316.7 | 0.67 | 1.16 | 0.68 | 0.21 | 0.32 | 0.21 |
| **23S rRNA** | | | | | | | | | | |
| WT R1 | 1050.1 | 1686.9 | 1059.5 | 869.1 | 0.90 | N.D. | 2.76 | 0.10 | N.D. | 0.32 |
| WT R2 | 906.7 | 1449.4 | 912.7 | 741.9 | 0.82 | N.D. | 2.48 | 0.11 | N.D. | 0.33 |
| WT R3 | 993.7 | 1597.5 | 1003.1 | 812.9 | 0.82 | N.D. | 2.49 | 0.10 | N.D. | 0.31 |
| *ΔyfjO* R1 | 382.2 | 626.8 | 298.1 | 252.3 | 0.27 | N.D. | 0.50 | 0.11 | N.D. | 0.20 |
| *ΔyfjO* R2 | 251.9 | 414.6 | 194.9 | 164.1 | 0.14 | N.D. | 0.31 | 0.09 | N.D. | 0.19 |
| *ΔyfjO* R3 | 413.7 | 663.3 | 319.5 | 269.6 | 0.23 | N.D. | 0.57 | 0.09 | N.D. | 0.21 |
| *ΔyfjO+yfjO*$^+$ R1 | 414.0 | 671.8 | 319.0 | 268.8 | 0.22 | N.D. | 0.82 | 0.08 | N.D. | 0.31 |
| *ΔyfjO+yfjO*$^+$ R2 | 497.3 | 877.8 | 382.2 | 323.7 | 0.25 | N.D. | 1.00 | 0.08 | N.D. | 0.31 |
| *ΔyfjO+yfjO*$^+$ R3 | 400.6 | 716.6 | 303.4 | 259.9 | 0.23 | N.D. | 0.85 | 0.09 | N.D. | 0.33 |

**Table S8.** Sporulation/germination efficiency of *B. subtilis* WT and *yfjO* Knockout strains was measured as the ratio of colony forming unit (CFU) from fully sporulated cultures before and after heat treatment (40 min at 90 °C). Cultures were diluted by $10^{-5}$, and 100 µL was plated on LB agar and incubated at 30 °C for overnight. Sporulation/germination (%) = (CFU heat treated/CFU unheated) x 100. R, replicate.

| ***B. subtilis* Strain** | **CFU unheated** | **CFU heated** | **Sporulation/ Germination (%)** |
|---|---|---|---|
| WT 168 R1 | 65 | 48 | 73.85 |
| WT 168 R2 | 115 | 64 | 55.65 |
| WT 168 R3 | 42 | 26 | 61.90 |
| WT 168 R4 | 65 | 35 | 53.85 |
| WT 168 R5 | 57 | 28 | 49.12 |
| *ΔyfjO::kan* BKK08020 R1 | 84 | 24 | 28.57 |
| *ΔyfjO::kan* BKK08020 R2 | 105 | 18 | 17.14 |
| *ΔyfjO::kan* BKK08020 R3 | 87 | 14 | 16.09 |
| *ΔyfjO::kan* BKK08020 R4 | 97 | 16 | 16.49 |
| *ΔyfjO::kan* BKK08020 R5 | 80 | 22 | 27.5 |
| *ΔyfjO* #1 R1 | 68 | 33 | 48.53 |
| *ΔyfjO* #1 R2 | 64 | 43 | 67.19 |
| *ΔyfjO* #2 R1 | 60 | 35 | 58.33 |
| *ΔyfjO* #2 R2 | 35 | 24 | 68.57 |

**Table S9.** Multiple mutations were detected in *ΔyfjO::kan (*BKK08020) relative to WT. Few mutations were identified within coding regions, including changes in sporulation-associated factors (*prpC*, *bmrD*), ribosomal proteins (*rplW*), and motility-related genes (*swrAA*). Whole genome sequencing alignment of *B. subtilis* WT and *ΔyfjO::kan (*BKK08020). Nanopore sequencing reads were aligned to reference genome (GeneBank assembly GCA_000009045.1).

| # | Genome position | Type of mutation | Reference base | New base | Protein level | Gene annotation |
|---|---|---|---|---|---|---|
| 1 | 137,019 | Substitution | C | T | Arg10Cys | *rplW* |
| 2 | 163.950 | Substitution | G | A | - | *rrnI-23S* |
| 3 | 172,772 | Substitution | C | A | - | *rrnG-16S* |
| 4 | 216,044 | Substitution | G | A | Arg214Gln | *skfC* |
| 5 | 531,727 | Substitution | A | G | - | - |
| 6 | 873,404-874,799 | Deletion -1395 bp Insertion +1066 bp | - | - | - | *ΔyfjO::kan*$^r$ |
| 7 | 1,048,569 | Substitution | C | T | Gln500* | *bmrD* |
| 8 | 1,650,847 | Substitution | A | G | Asp155Gly | *prpC* |
| 9 | 2,057,703 | Substitution | C | T | - | - |
| 10 | 2,057,726 | Substitution | T | G | - | - |
| 11 | 2,057,811 | Substitution | A | G | Lys4Arg | *yobE* |
| 12 | 2,057,857 | Substitution | C | T | - | *yobE* |
| 13 | 2,057,947 | Substitution | T | C | - | *yobE* |
| 14 | 2,057,962 | Substitution | T | C | - | *yobE* |
| 15 | 2,057,992 | Substitution | T | G | Cys64Trp | *yobE* |
| 16 | 2,058,013 | Substitution | T | C | - | *yobE* |
| 17 | 2,058,031 | Substitution | T | C | - | *yobE* |
| 18 | 2,058,078 | Substitution | G | A | Gly93Val | *yobE* |
| 19 | 2,058,175 | Substitution | G | C | - | *yobE* |
| 20 | 2,058,184 | Substitution | T | C | - | *yobE* |
| 21 | 2,058,202 | Substitution | T | C | - | *yobE* |
| 22 | 2,058,217 | Substitution | G | A | - | *yobE* |
| 23 | 2,058,219 | Substitution | T | C | Leu140Pro | *yobE* |
| 24 | 2,058,231 | Substitution | T | C | Leu144Pro | *yobE* |
| 25 | 2,058,232 | Substitution | G | T | Leu144Pro | *yobE* |
| 26 | 2,058,253 | Substitution | C | T | - | *yobE* |
| 27 | 2,058,258 | Substitution | T | C | Ile153Thr | *yobE* |
| 28 | 2,058,267 | Substitution | G | A | Ser156Asn | *yobE* |
| 29 | 2,058,286 | Substitution | T | C | - | *yobE* |
| 30 | 2,058,310 | Substitution | A | T | - | *yobE* |
| 31 | 2,058,323 | Substitution | A | G | Lys175Glu | *yobE* |
| 32 | 2,058,337 | Substitution | A | C | - | *yobE* |
| 33 | 2,058,340 | Substitution | T | C | - | *yobE* |

| | | | | | | |
|---|---|---|---|---|---|---|
| 34 | 2,058,376 | Substitution | G | A | - | *yobE* |
| 35 | 2,058,377 | Substitution | T | C | - | *yobE* |
| 36 | 2,058,381 | Substitution | T | A | Leu192Gln | *yobE* |
| 37 | 2,058,385 | Substitution | T | A | - | *yobE* |
| 38 | 2,058,412 | Substitution | T | C | - | *yobE* |
| 39 | 2,058,418 | Substitution | A | T | - | *yobE* |
| 40 | 2,058,421 | Substitution | T | A | - | *yobE* |
| 41 | 2,058,424 | Substitution | A | T | - | *yobE* |
| 42 | 2,058,487 | Substitution | C | T | - | - |
| 43 | 2,058,492 | Substitution | C | T | - | - |
| 44 | 2,058,525 | Substitution | G | A | - | - |
| 45 | 2,434,323 | Substitution | G | A | - | *ypuB* |
| 46 | 2,555,972 | Insertion +1 | - | C | Frame shift | *spoIITA* |
| 47 | 2,708,068 | Insertion +1 | - | C | - | - |
| 48 | 3,289,092 | Substitution | C | T | Arg390Gln | *dhbE* |
| 49 | 3,621,947 | Deletion -1 | T | - | Frame shift | *swrAA* |
| 50 | 3,829,788 | Substitution | G | A | Thr57Ile | *narG* |

**Table S10.** Competitive growth of WT versus *ΔyfjO* strains. Quantification of PCR product densitometry from gels shown in Figure S21. A 1:1 mixed culture ($P_0$) was grown for 18 hours at 30 °C, 37 °C, or 42 °C, followed by serial passaging. The *yfjO* region was amplified by PCR; WT and *ΔyfjO* products are 1464 bp (upper band) and 1135 bp (lower band), respectively. Passage (P) 7, 14, or 23 were analyzed along with cultures of known WT:*ΔyfjO* ratios. WT % and *ΔyfjO* % were calculated as the percentage of total densitometry signal from PCR bands. R, replicate.

| | **WT:*ΔyfjO*** | **WT product densitometry** | ***ΔyfjO* product densitometry** | **WT %** | ***ΔyfjO* %** |
|---|---|---|---|---|---|
| **calibration** | 100:0 | 9581.5 | 0 | 100 | 0 |
| | 90:10 | 7802.6 | 1761.0 | 81.6 | 18.4 |
| | 75:25 | 7010.5 | 3758.3 | 65.1 | 34.9 |
| | 50:50 | 5439.9 | 5559.9 | 49.5 | 50.5 |
| | 25:75 | 3064.5 | 6183.3 | 33.2 | 66.8 |
| | 10:90 | 2017.7 | 8894.7 | 18.5 | 81.5 |
| | 0:100 | 0 | 8919.7 | 0 | 100 |
| **$P_0$** | $P_0$ R1 | 5439.9 | 5559.9 | 49.5 | 50.5 |
| | $P_0$ R2 | 8513.9 | 8848.9 | 49.0 | 51.0 |
| | $P_0$ R3 | 8657.0 | 8592.8 | 50.2 | 49.8 |
| **30 °C** | $P_7$ R1 | 8019.0 | 2739.5 | 74.5 | 25.5 |
| | $P_7$ R2 | 7073.8 | 2430.0 | 74.4 | 25.6 |
| | $P_7$ R3 | 7243.5 | 2593.3 | 73.6 | 26.4 |
| | $P_{14}$ R1 | 9454.5 | 1335.4 | 87.6 | 12.4 |
| | $P_{14}$ R2 | 9710.6 | 1088.8 | 89.9 | 10.1 |
| | $P_{14}$ R3 | 7721.6 | 963.4 | 88.9 | 11.1 |
| | $P_{23}$ R1 | 9714.4 | 215.0 | 97.8 | 2.2 |
| | $P_{23}$ R2 | 11875.1 | 346.4 | 97.2 | 2.8 |
| | $P_{23}$ R3 | 10339.8 | 2120.3 | 83.0 | 17.0 |
| **37 °C** | $P_7$ R1 | 5229.9 | 2049.8 | 71.8 | 28.2 |
| | $P_7$ R2 | 5814.2 | 1998.3 | 74.4 | 25.6 |
| | $P_7$ R3 | 7740.1 | 2570.5 | 75.1 | 24.9 |
| | $P_{14}$ R1 | 10414.9 | 795.7 | 92.9 | 7.1 |
| | $P_{14}$ R2 | 11142.7 | 0 | 100 | 0 |
| | $P_{14}$ R3 | 9965.4 | 851.8 | 92.1 | 7.9 |
| | $P_{23}$ R1 | 10581.9 | 88.4 | 99.2 | 0.8 |
| | $P_{23}$ R2 | 9784.8 | 0 | 100 | 0 |
| | $P_{23}$ R3 | 11560.9 | 209.2 | 98.2 | 1.8 |
| **42 °C** | $P_7$ R1 | 7279.0 | 4301.1 | 62.9 | 37.1 |
| | $P_7$ R2 | 8611.5 | 4849.7 | 64.0 | 36.0 |
| | $P_7$ R3 | 8272.3 | 5468.8 | 60.2 | 39.8 |
| | $P_{14}$ R1 | 9233.8 | 3276.3 | 73.8 | 26.2 |
| | $P_{14}$ R2 | 9701.2 | 2880.6 | 77.1 | 22.9 |
| | $P_{14}$ R3 | 10031.2 | 3626.9 | 73.4 | 26.6 |
| | $P_{23}$ R1 | 11607.9 | 1363.0 | 89.5 | 10.5 |
| | $P_{23}$ R2 | 12915.9 | 1006.8 | 92.8 | 7.2 |
| | $P_{23}$ R3 | 11792.9 | 2149.0 | 84.6 | 15.4 |

**Table S11.** Global protein translation quantification in *B. subtilis* WT, *ΔyfjO*, and *ΔyfjO+yfjO*$^+$ strains. Protein synthesis was measured using O-propargyl-puromycin (OP-Puro) labelling. Quantification of relative Cy5 intensity from fluorescent gel imaging and Coomassie-stained gels appears in Figure S22 are shown below. Cultures were labeled with OP-Puro at 37 °C for 30 minutes, followed by cell lysis and CuAAC click chemistry with Cy5-azide. Signals were normalized to the WT R1 signal. Relative Cy5 intensity = Normalized Cy5 signal/ Normalized Coomassie signal. Pre Puro, pretreatment with puromycin; Pre Cm, pretreatment with chloramphenicol. R, replicate.

| | Sample | Coomassie signal | Cy5 signal | Normalized Coomassie signal | Normalized Cy5 signal | Relative Cy5 intensity |
|---|---|---|---|---|---|---|
| **Gel 1** | No OP-Puro | 4271913 | 36159 | 1.00751786 | 0.03412778 | 0.03387313 |
| | WT R1 | 4240037 | 1059518 | 1 | 1 | 1 |
| | WT R2 | 4107884 | 928983 | 0.96883211 | 0.87679775 | 0.90500484 |
| | WT R3 | 4275810 | 1033194 | 1.00843695 | 0.97515474 | 0.96699624 |
| | No OP-Puro | 4786103 | 35001 | 1.12878803 | 0.03303483 | 0.02926575 |
| | *ΔyfjO* R1 | 4104636 | 582732 | 0.96806608 | 0.54999726 | 0.56814021 |
| | *ΔyfjO* R2 | 4958147 | 1032593 | 1.16936409 | 0.9745875 | 0.83343375 |
| | *ΔyfjO* R3 | 5198662 | 916463 | 1.22608883 | 0.86498106 | 0.70547993 |
| | WT Pre Puro | 6165116 | 130170 | 1.4540241 | 0.12285775 | 0.08449499 |
| | WT Pre Cm | 4693093 | 1386912 | 1.1068519 | 1.30900277 | 1.18263588 |
| | No click | 5461503 | 231 | 1.28807909 | 0.00021802 | 0.00016926 |
| **Gel 2** | WT R1 | 5107302 | 5414982 | 1 | 1 | 1 |
| | WT R4 | 5210925 | 5746522 | 1.02028919 | 1.06122643 | 1.04012317 |
| | WT R5 | 5382602 | 5324626 | 1.05390322 | 0.9833137 | 0.93302088 |
| | WT R6 | 5151323 | 5745064 | 1.00861923 | 1.06095717 | 1.05189069 |
| | *ΔyfjO* R4 | 5461503 | 4812010 | 1.06935188 | 0.88864746 | 0.83101501 |
| | *ΔyfjO* R5 | 5370131 | 4736502 | 1.05146142 | 0.87470318 | 0.8318928 |
| | *ΔyfjO* R6 | 5655148 | 5380668 | 1.1072672 | 0.99366314 | 0.8974014 |
| | *ΔyfjO+yfjO*$^+$ R1 | 5159311 | 5287803 | 1.01018326 | 0.9765135 | 0.96666964 |
| | *ΔyfjO+yfjO*$^+$ R2 | 4322024 | 4070198 | 0.84624406 | 0.75165495 | 0.88822478 |
| | *ΔyfjO+yfjO*$^+$ R3 | 5187283 | 5265402 | 1.01566013 | 0.97237664 | 0.95738389 |
| | *ΔyfjO+yfjO*$^+$ R4 | 5338602 | 5650360 | 1.0452881 | 1.04346792 | 0.99825868 |
| | *ΔyfjO+yfjO*$^+$ R5 | 4804800 | 6046653 | 0.94077068 | 1.11665247 | 1.186955 |
| | *ΔyfjO+yfjO*$^+$ R6 | 5142307 | 5910130 | 1.00685391 | 1.09144038 | 1.08401066 |
| | WT Pre Puro | 5914184 | 858601 | 1.15798596 | 0.15856027 | 0.13692763 |

**Table S12.** Oligonucleotides used in this study.

| # | Name | Description | Sequence |
|---|---|---|---|
| Primers for PCR amplification | | | |
| 1 | P.01 (5pL) | Double recombination construct | CTAATTCCAGCGGCATGTTATAC |
| 2 | P.02 (3pR) | Double recombination construct | AAATAACCAACGCAGCAAGTG |
| 3 | P.03 | Confirm *yfjO* deletion and quantify WT:*ΔyfjO* ratio | AAAGCTTCATATAGAAACGGAG |
| 4 | P.04 | Confirm *yfjO* deletion and quantify WT:*ΔyfjO* ratio | GGAAAACTATTTTATGACTGGGA |
| 5 | P.05 | Confirm *amyE* insertion | TTCTTCGCTTGGCTGAAAAT |
| 6 | P.06 | Confirm *amyE* insertion | CACCAGGTTTTTGGTTTGCT |
| **2’-deoxy-2’-O-methyl Chimeric oligonucleotides** | | | |
| 7 | CO–616-623 | 23S-rRNA–*B. subtilis* | CmCmGmGmTTCAUmUmCmUmACAAAmAm |
| 8 | CO–566-583 | 23S-rRNA–*E. coli* | CmUmGmAmCCCAUmUmAmUmACAAAmAm |

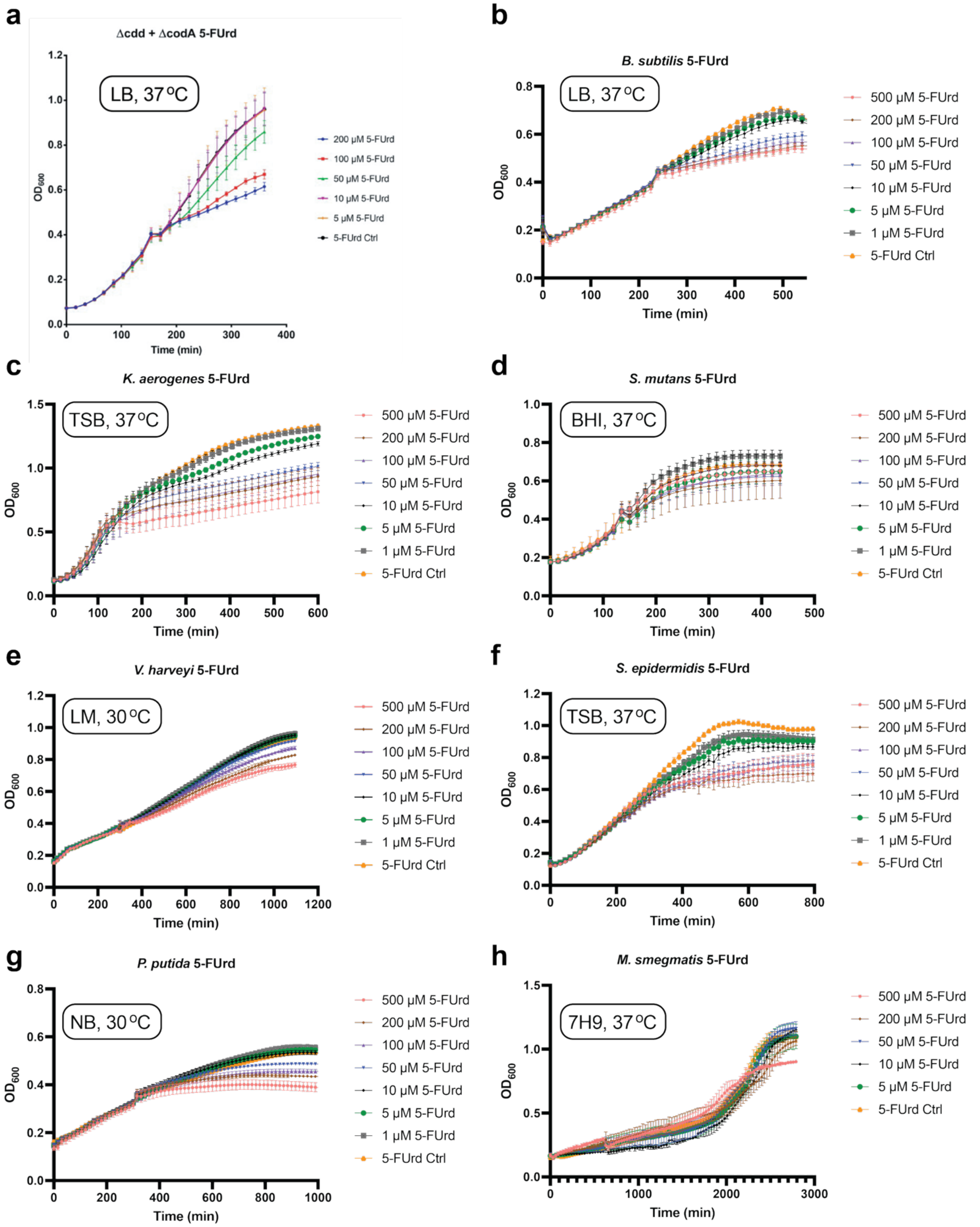


**Figure S1.** Bacterial growth curves in the presence of 5-FUrd. Cultures of individual bacterial strains **(a)**-**(h)** were inoculated and grown in the indicated medium and temperature, followed by addition of 5-FUrd (1-500 µM) when the culture reached OD ~0.4. OD600 measurements were recorded every 15 minutes using a plate reader until early stationary phase. Two independent biological replicates with three technical replicates each were analyzed; data represent mean values ± s.e.m.

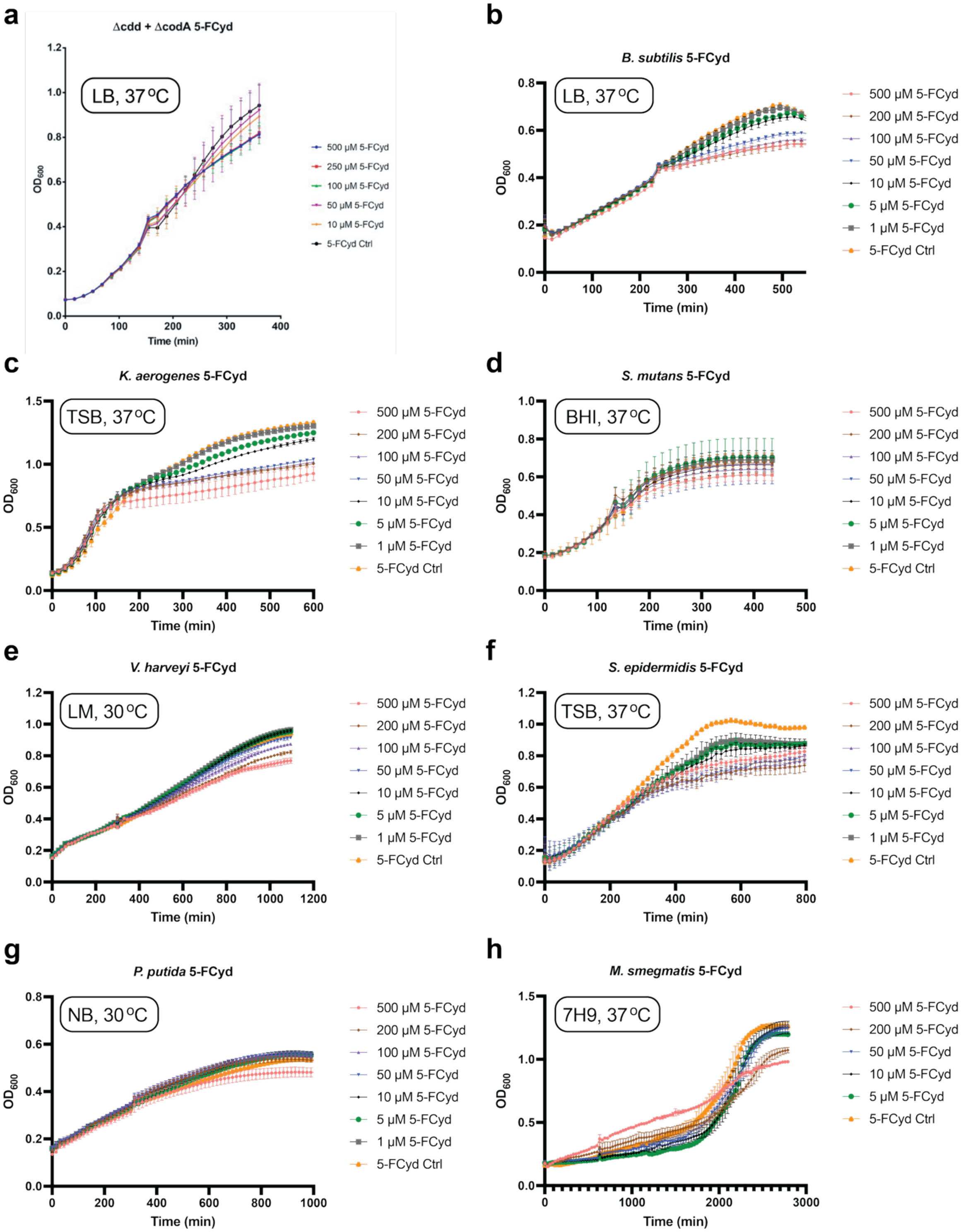


**Figure S2.** Bacterial growth curves in the presence of 5-FCyd. Cultures of individual bacterial strains **(a)**-**(h)** were inoculated and grown in the indicated medium and temperature, followed by addition of 5-FCyd (1-500 µM) when the culture reached OD ~0.4. OD600 measurements were recorded every 15 minutes using a plate reader until early stationary phase. Two independent biological replicates with three technical replicates each were analyzed; data represent mean values ± s.e.m.

A: 500 - 10 ng/mL

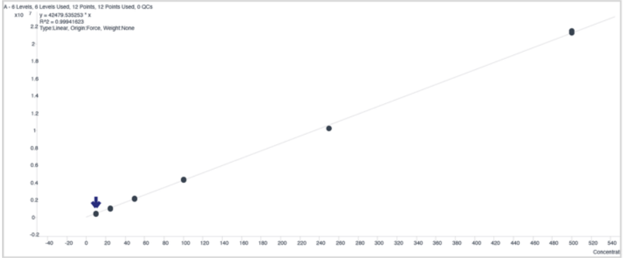

G: 500 - 10 ng/mL

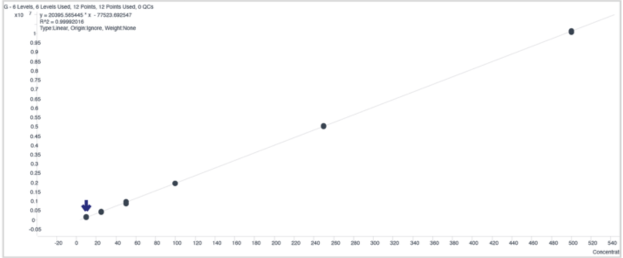

C: 500 - 10 ng/mL

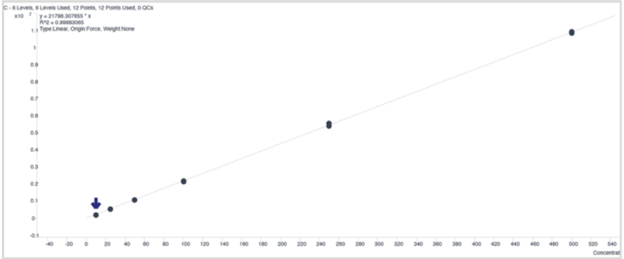

U: 500 - 10 ng/mL

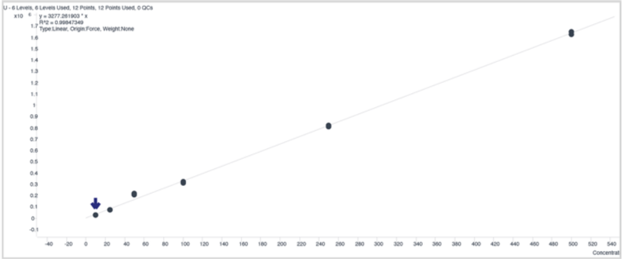

5-FCyd: 50 - 0.5 ng/mL

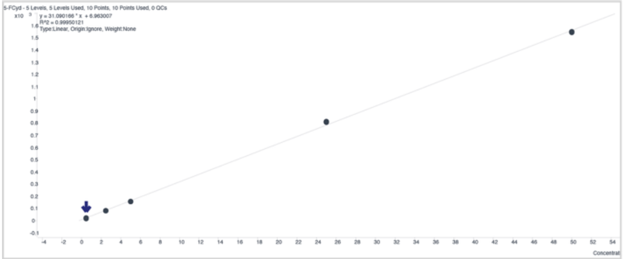

5-FUrd: 50 - 0.5 ng/mL

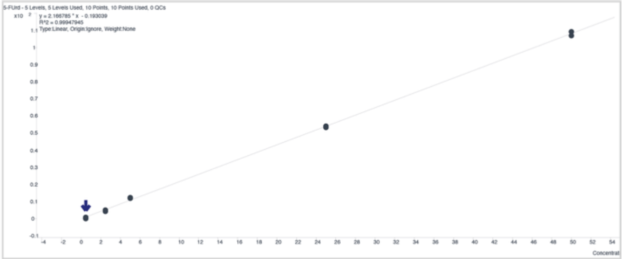

$m^5C$: 50 - 1 ng/mL

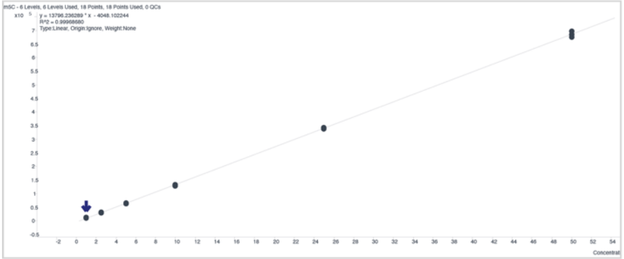

$m^5U$: 50 -1 ng/mL

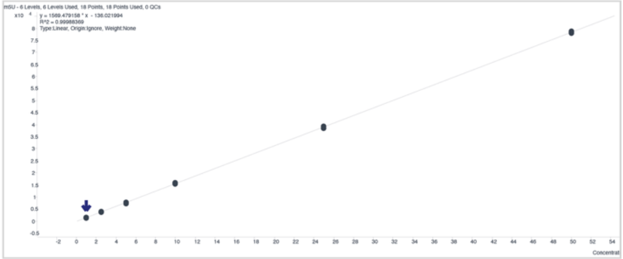

DHU: 50 - 1 ng/mL

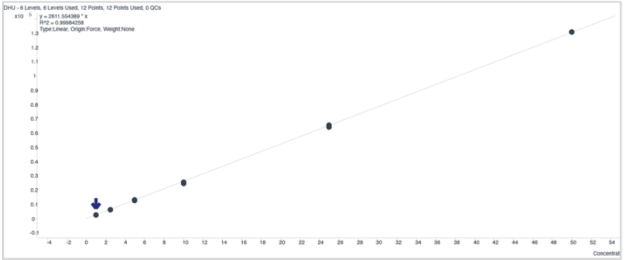

**Figure S3.** Representative calibration curves for canonical and modified nucleosides analyzed by LC-QQQ-MS.

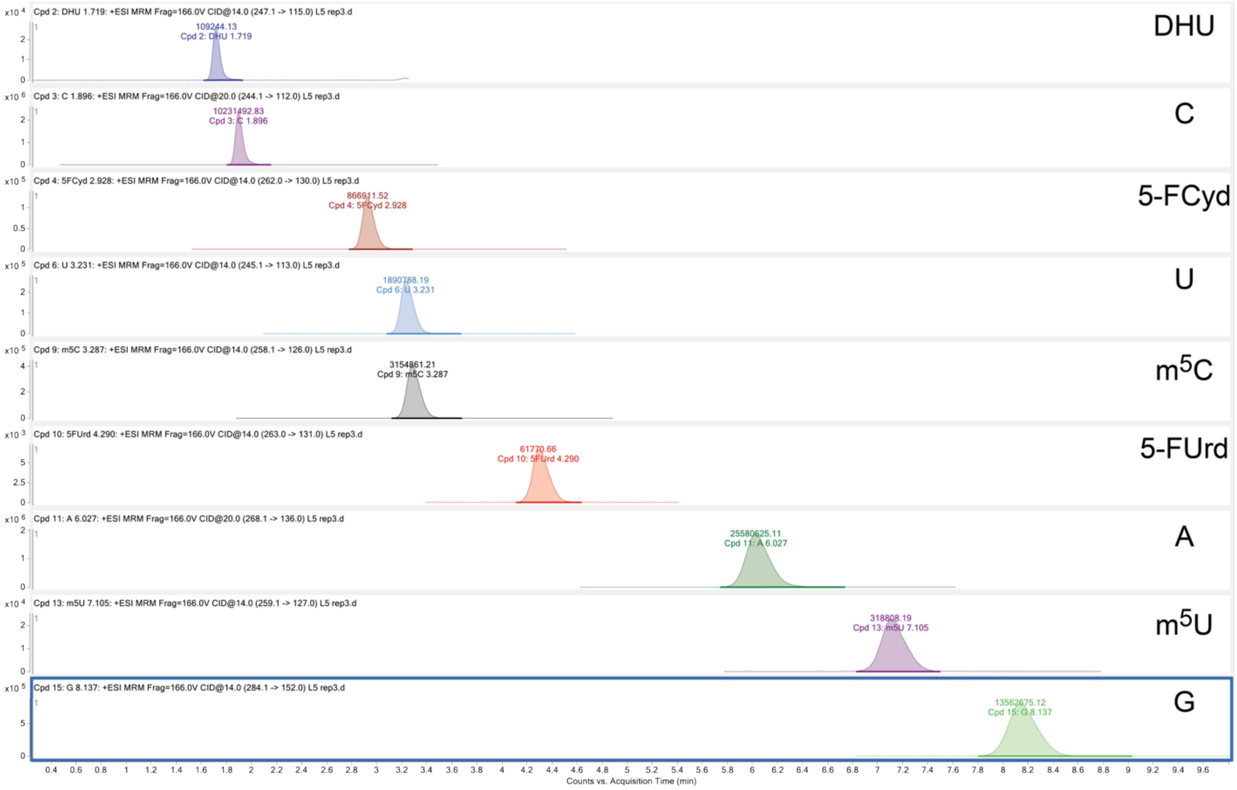


**Figure S4.** Representative chromatogram from the nucleoside LC-QQQ-MS analysis.

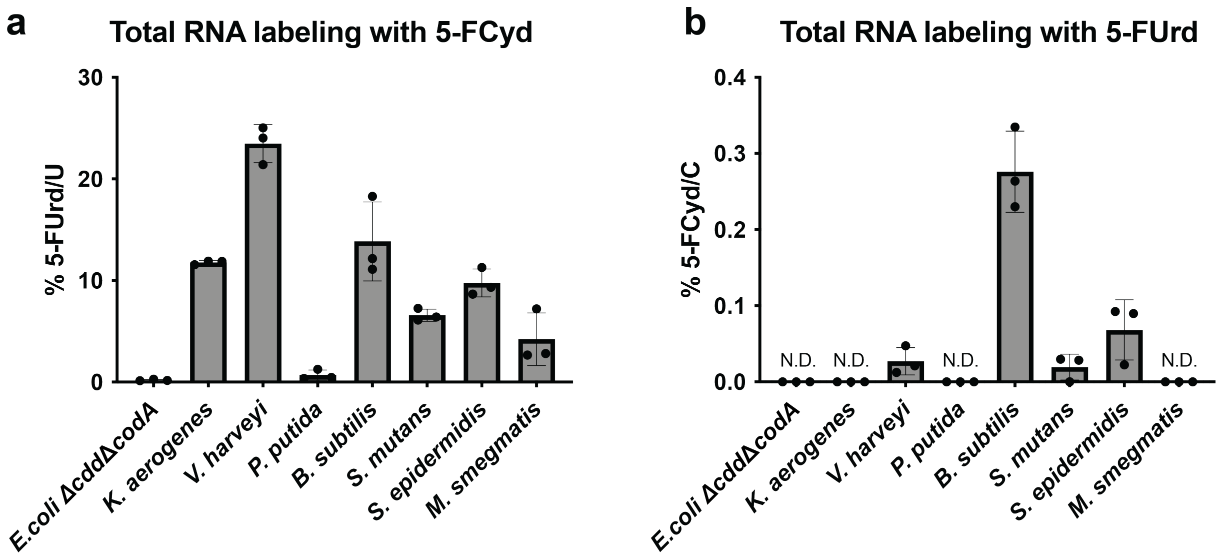


**Figure S5.** Metabolic conversion between 5-FCyd and 5-FUrd in total RNA measured by LC-QQQ-MS after metabolic labelling of panel of bacteria. **(a)** 5-FUrd levels following treatment with 5-FCyd. **(b)** 5-FCyd levels following treatment with 5-FUrd. Three independent biological replicates were analyzed. Data are representative of mean values ± s.e.m. Levels of 5-FCyd or 5-FUrd are indicated as a percentage of total Cyd or Urd, respectively. N.D., not detected.

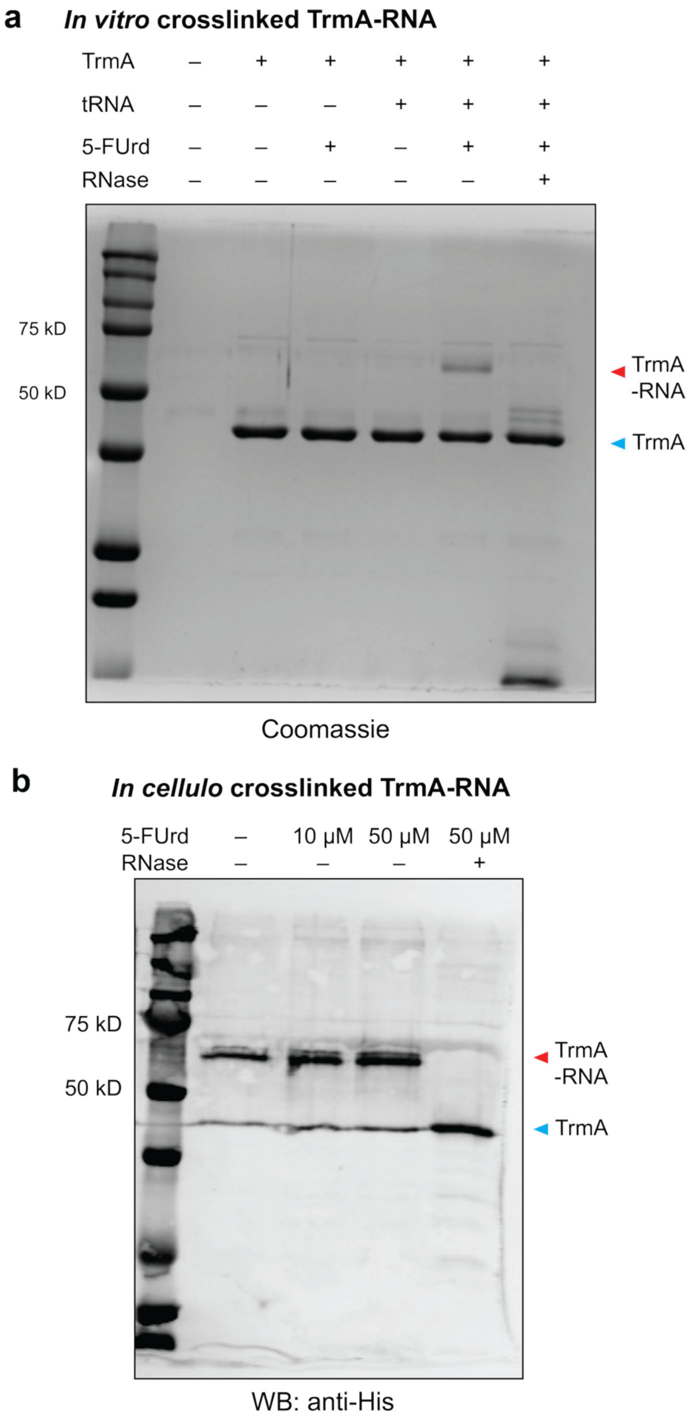


**Figure S6.** TrmA crosslinking with 5-FUrd-modified RNA. **(a)** Full Coomassie-stained gel showing RNA-protein crosslinking after *in vitro* methylation reaction between recombinant TrmA and small RNA isolated from 5-FUrd treated *E. coli ΔtrmA* cells. **(b)** Western blot analysis of TrmA-RNA complex present in interphase samples following metabolic labeling with 5-FUrd. *E. coli ΔtrmA* cells expressing TrmA-6xHis were treated with 10 μM or 50 μM 5-FUrd for 6 hours or left untreated, followed by three rounds of interphase extractions. Samples were analyzed by Western blot using anti-His antibody.

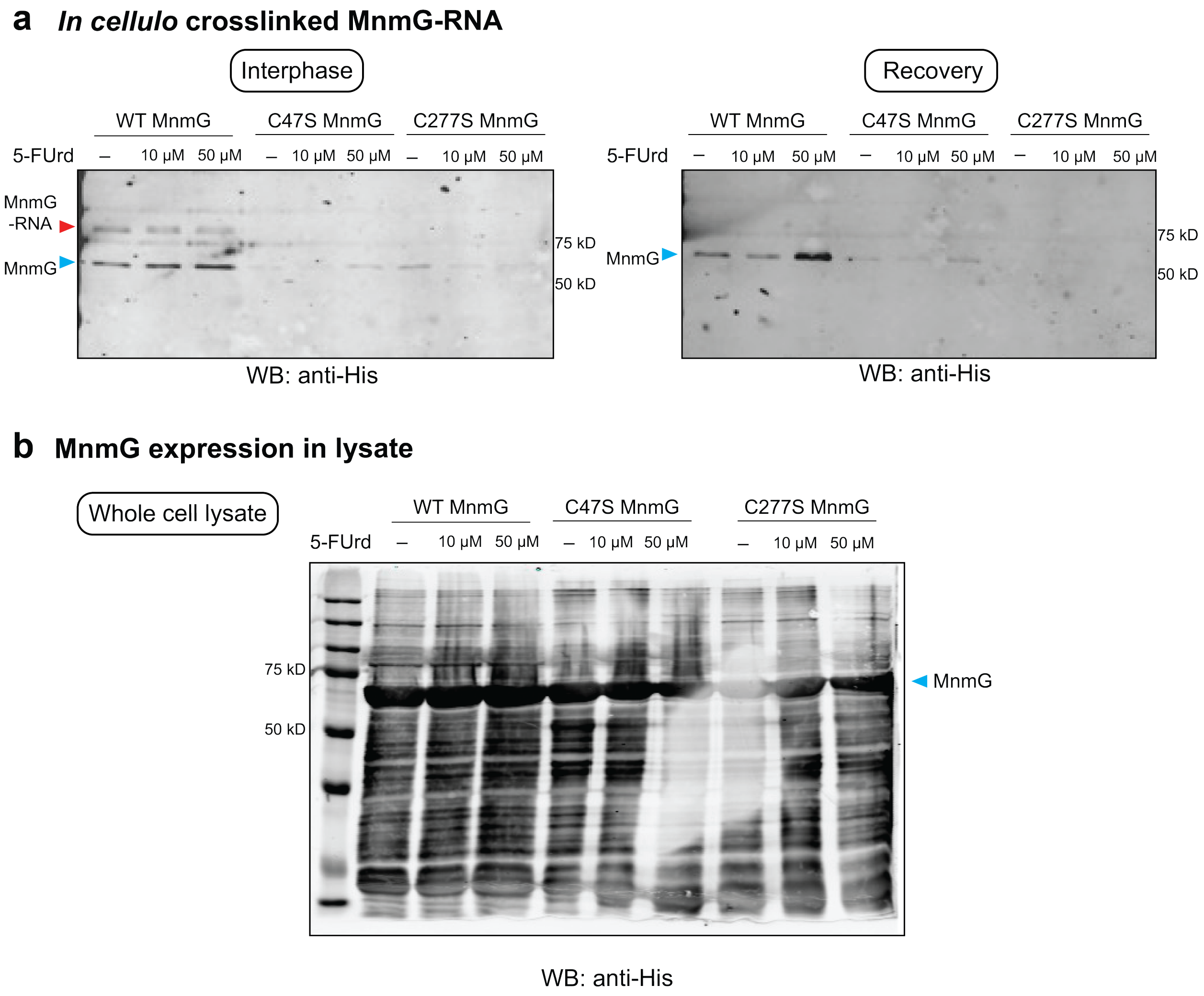


**Figure S7.** Analysis of MnmG-RNA crosslinking and protein expression. **(a)** Western blot analysis of MnmG-RNA complex in interphase samples following metabolic labeling with 5-FUrd (second replicate). Left: after three rounds of interphase extractions; right: after MnmG recovery following RNase digestion. **(b)** Western blot analysis of MnmG expression in whole cell lysates. *E. coli ΔmnmG* cells expressing MnmG(WT)-6xHis, MnmG(C47S)-6xHis or MnmG(C277S)-6xHis were treated with 10 μM or 50 μM 5-FUrd for 6 hours or left untreated. Samples were analyzed by Western blot using anti-His antibody.

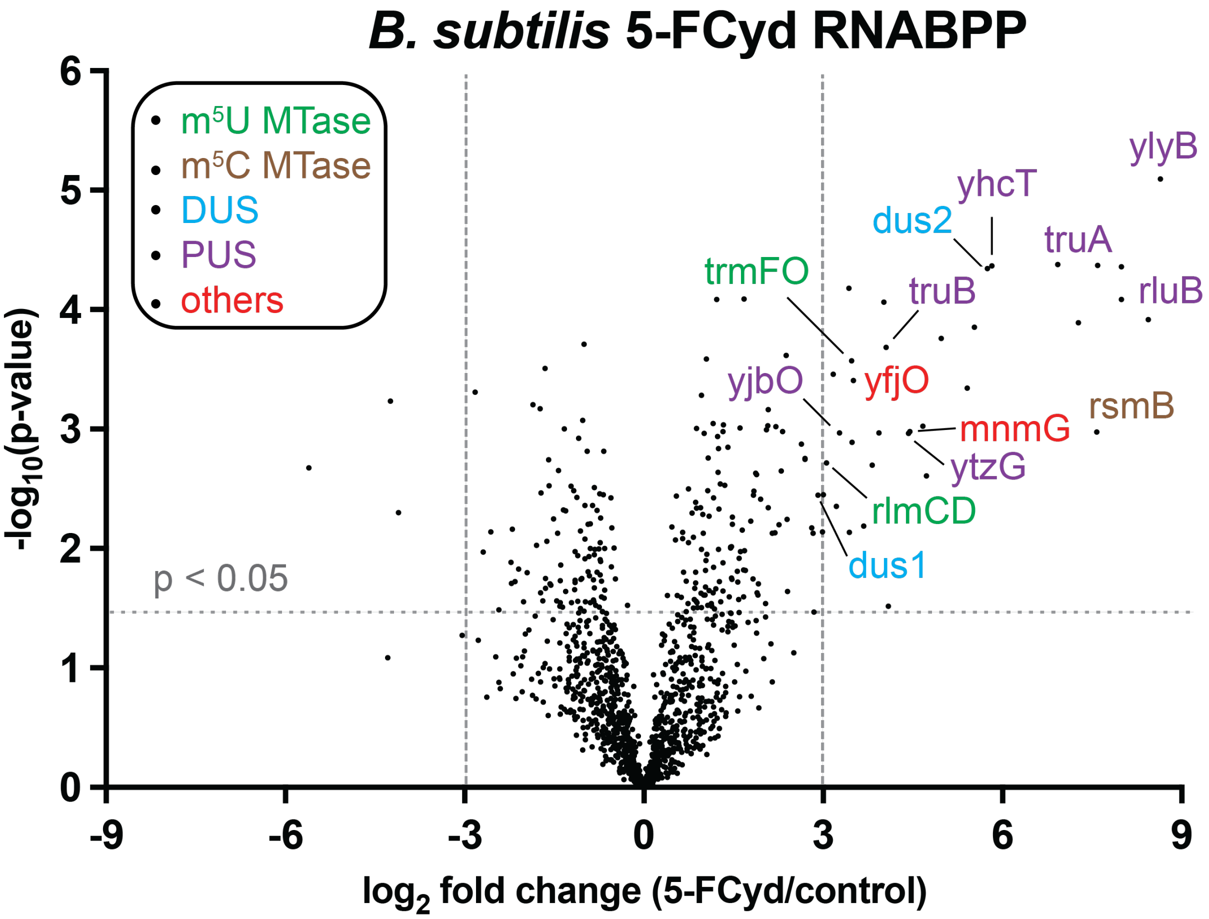


**Figure S8.** Proteomic profiling of 5-FCyd-reactive proteins using RNABPP-PS in *B. subtilis*. Volcano plot is presented; expected and unexpected RNA-modifying enzymes are annotated and classified.

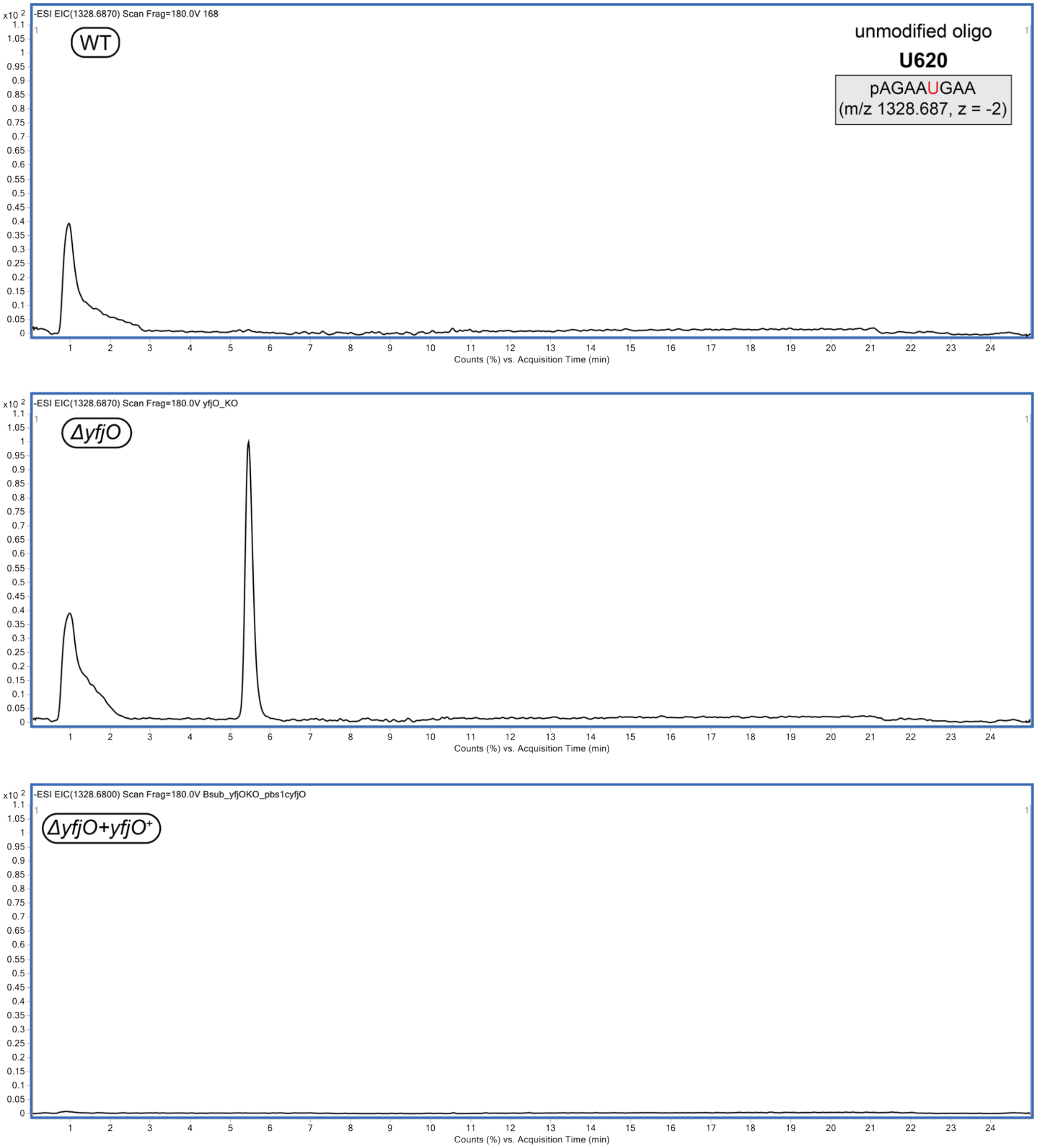


**Figure S9.** Double-cleavage oligonucleotide LC-MS analysis of *B. subtilis* 23S rRNA. Extracted ion chromatograms of unmodified oligonucleotide fragment generated from sequence-guided RNase H cleavage of 23S rRNA isolated from *B. subtilis* WT, *ΔyfjO,* or *ΔyfjO+yfjO*$^{+}$.

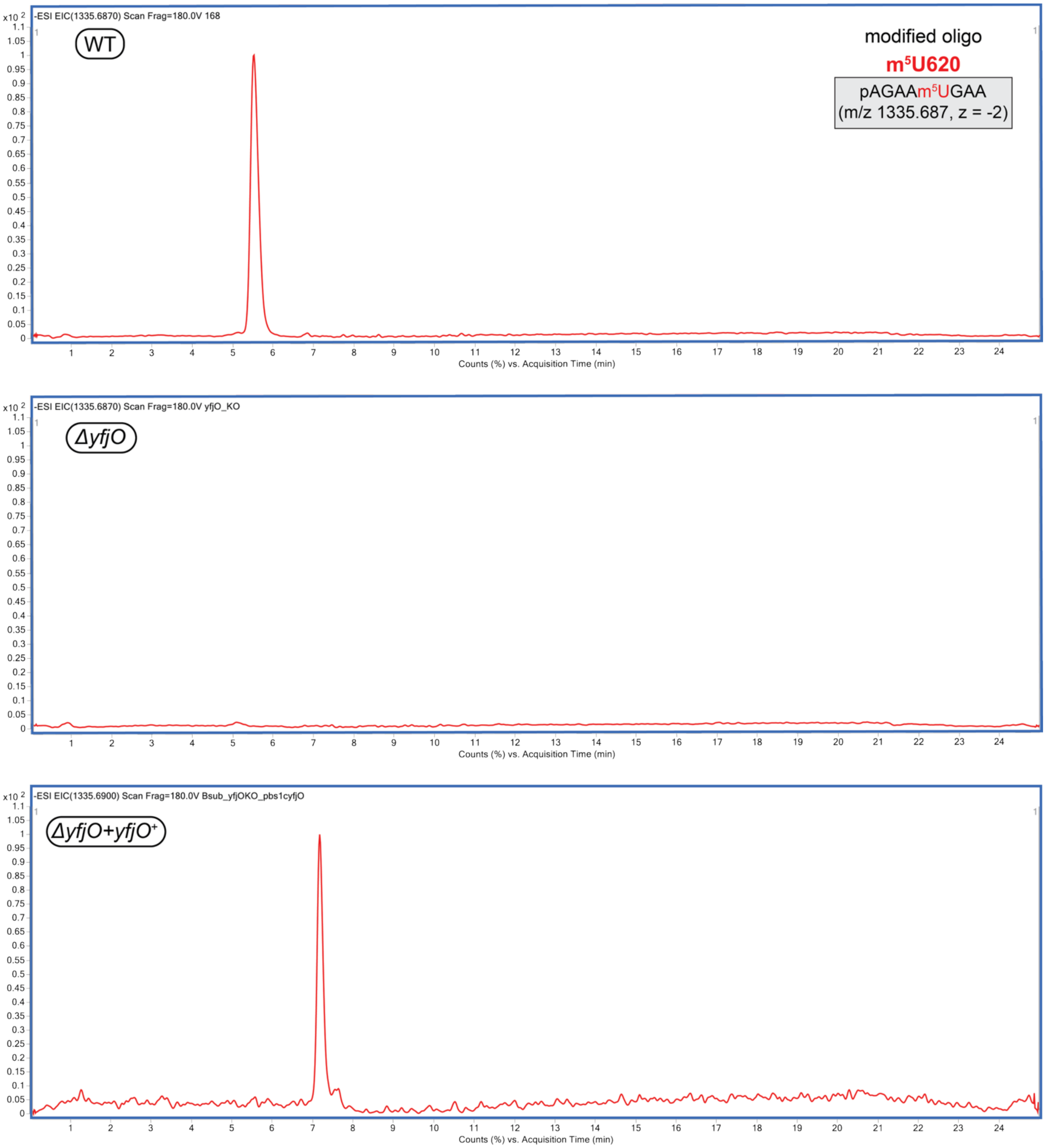


**Figure S10.** Double-cleavage oligonucleotide LC-MS analysis of *B. subtilis* 23S rRNA. Extracted ion chromatograms of $m^5U$-modified oligonucleotide fragment generated from sequence-guided RNase H cleavage of 23S rRNA isolated from *B. subtilis* WT, *ΔyfjO,* or *ΔyfjO+yfjO*$^+$.

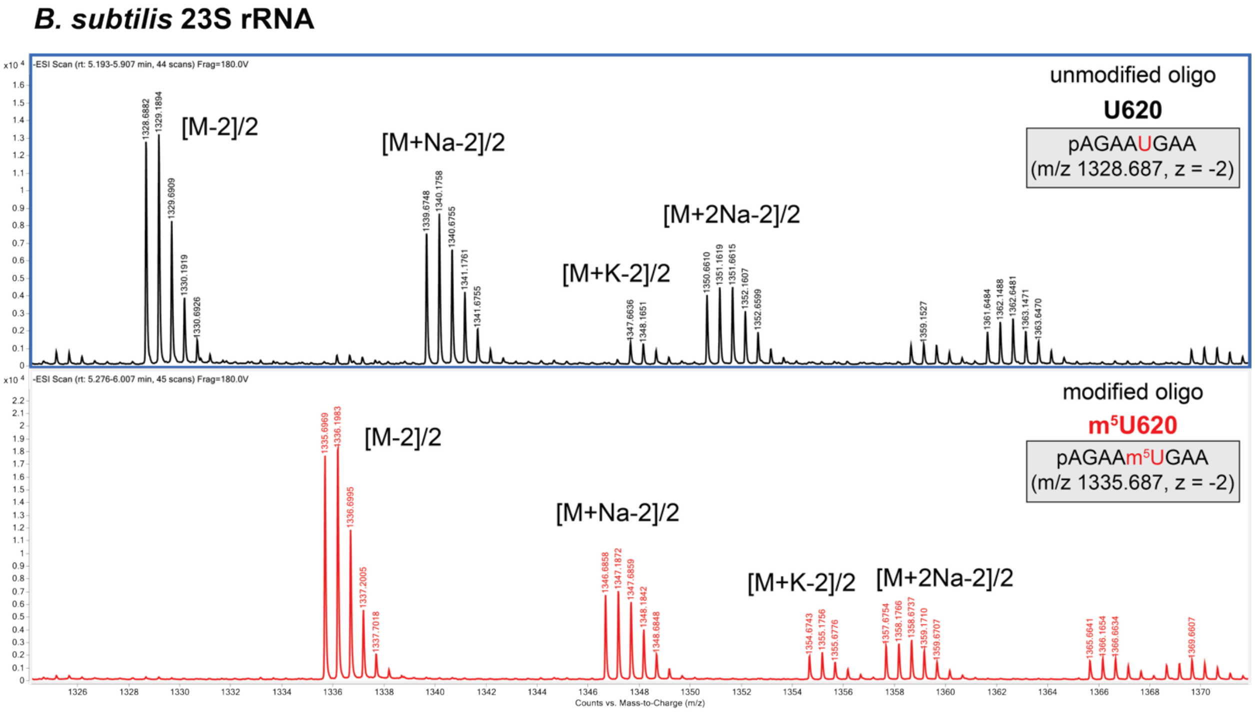


**Figure S11.** Representative MS spectra of oligonucleotide fragments produced from sequence-guided RNase H cleavage. Unmodified (*ΔyfjO*) and corresponding $m^5$U-modified (WT) oligonucleotide from *B. subtilis* 23S rRNA.

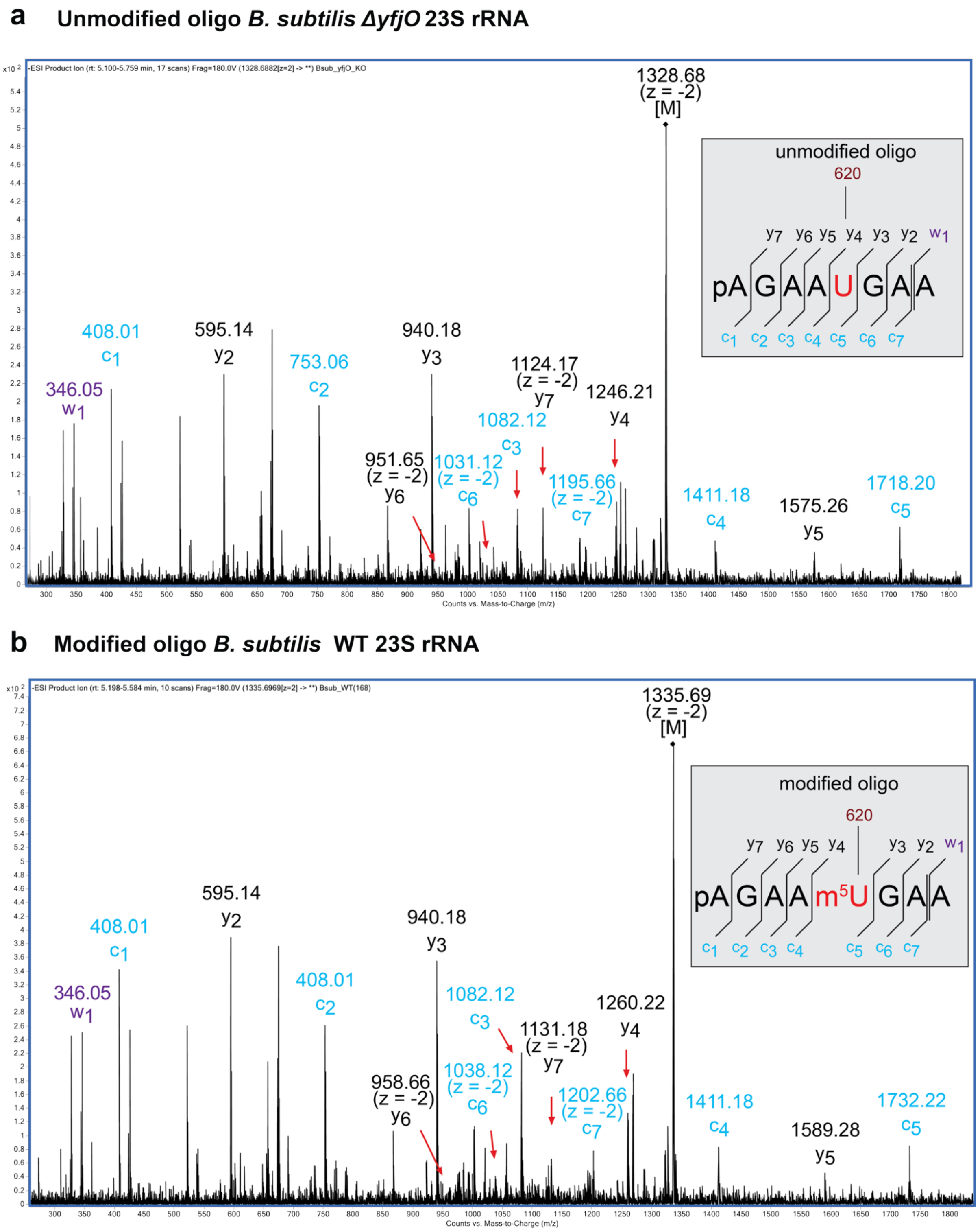


**Figure S12.** Representative MS/MS spectra of oligonucleotide fragments produced from sequence-guided RNase H cleavage. **(a)** Unmodified oligonucleotide fragment isolated from *B. subtilis ΔyfjO* 23S rRNA. Precursor ion is 1328.68 (z = -2). **(b)** Corresponding m$^5$U-modified oligonucleotide fragment isolated from *B. subtilis* WT 23S rRNA. Precursor ion is 1335.69 (z = -2).

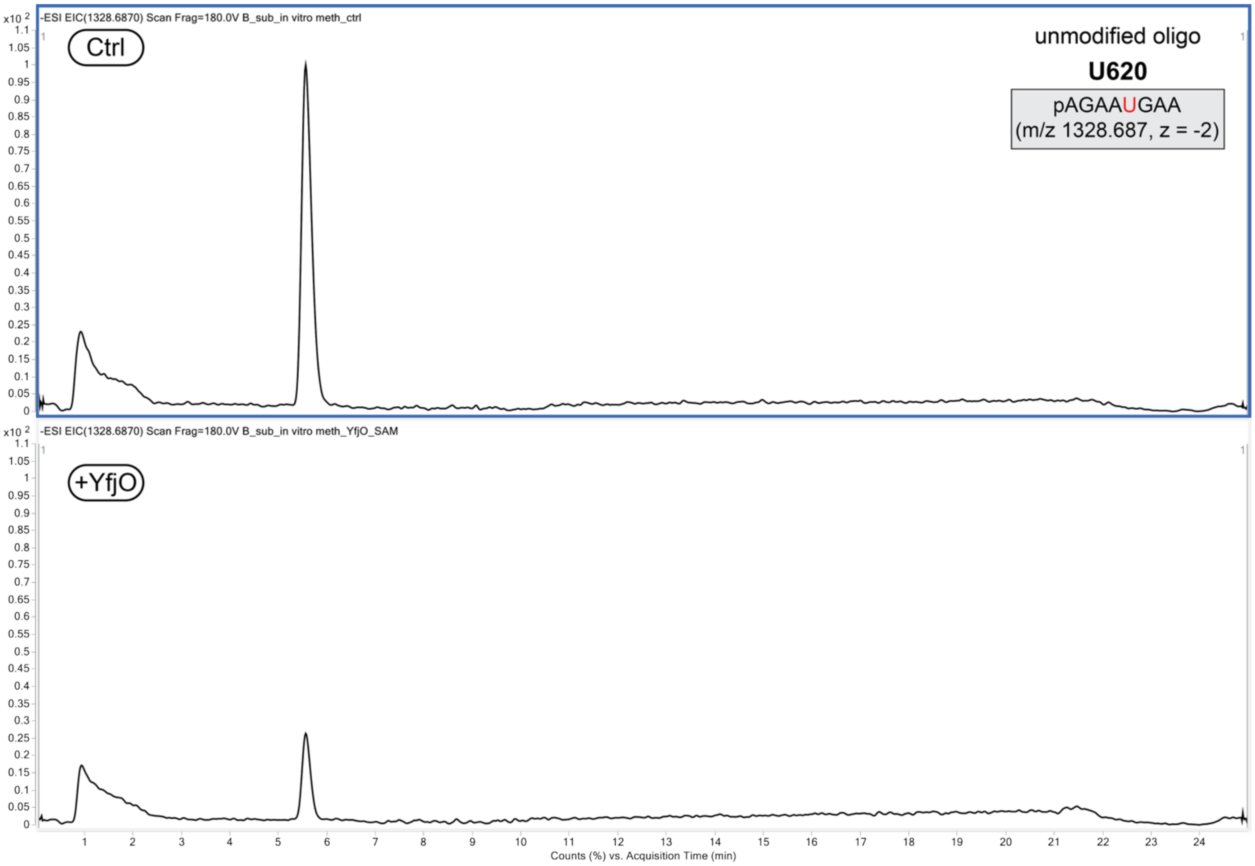


**Figure S13.** Double-cleavage LC-MS analysis of 23S rRNA after *in vitro* methylation reaction using recombinant YfjO and isolated 23S rRNA from *B. subtilis ΔyfjO*. Extracted ion chromatograms of unmodified oligonucleotide fragment generated from sequence-guided RNase H cleavage of -YfjO and +YfjO samples. Peak area: -YfjO: 2029350.9 +YfjO: 428835.9.

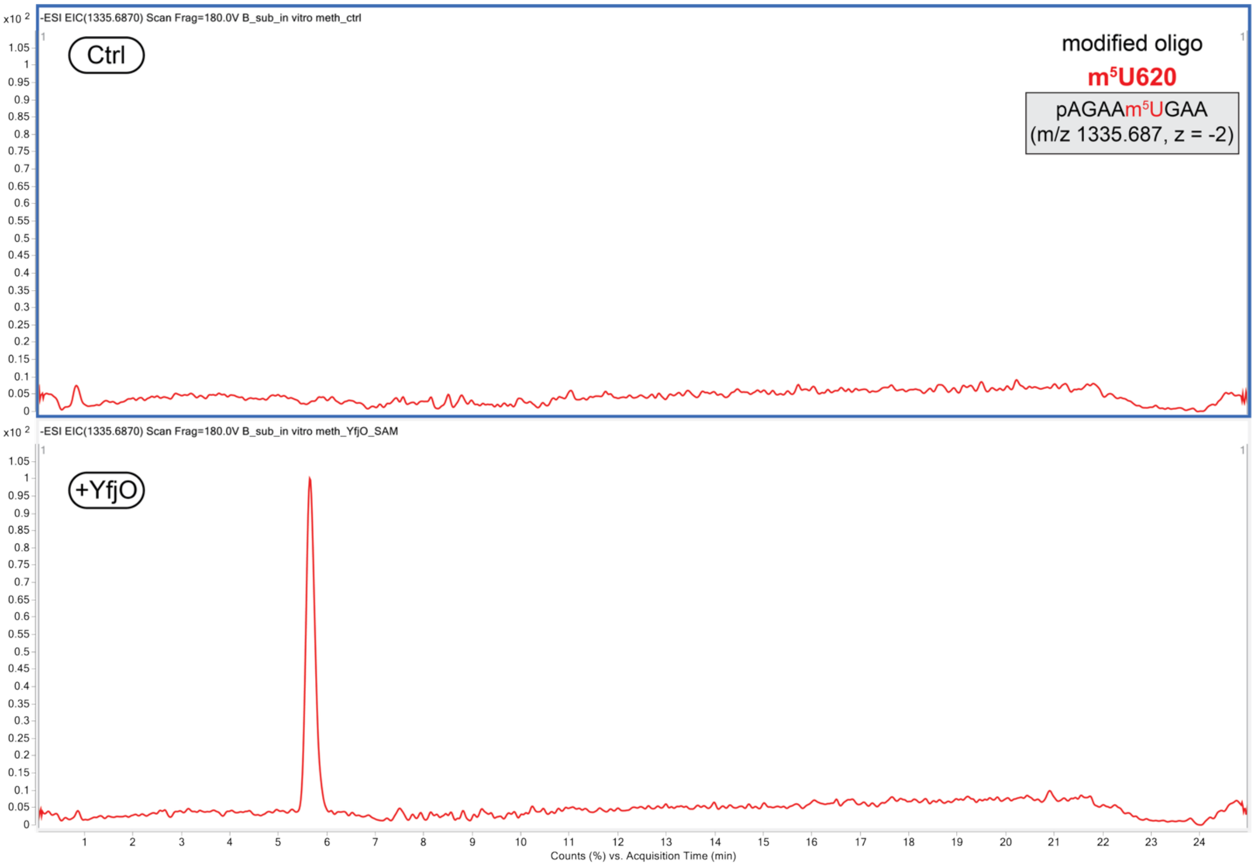


**Figure S14.** Double-cleavage LC-MS analysis of 23S rRNA after *in vitro* methylation reaction using recombinant YfjO and isolated 23S rRNA from *B. subtilis ΔyfjO*. Extracted ion chromatograms of m$^5$U-modified oligonucleotide fragment generated from sequence-guided RNase H cleavage of -YfjO and +YfjO samples.

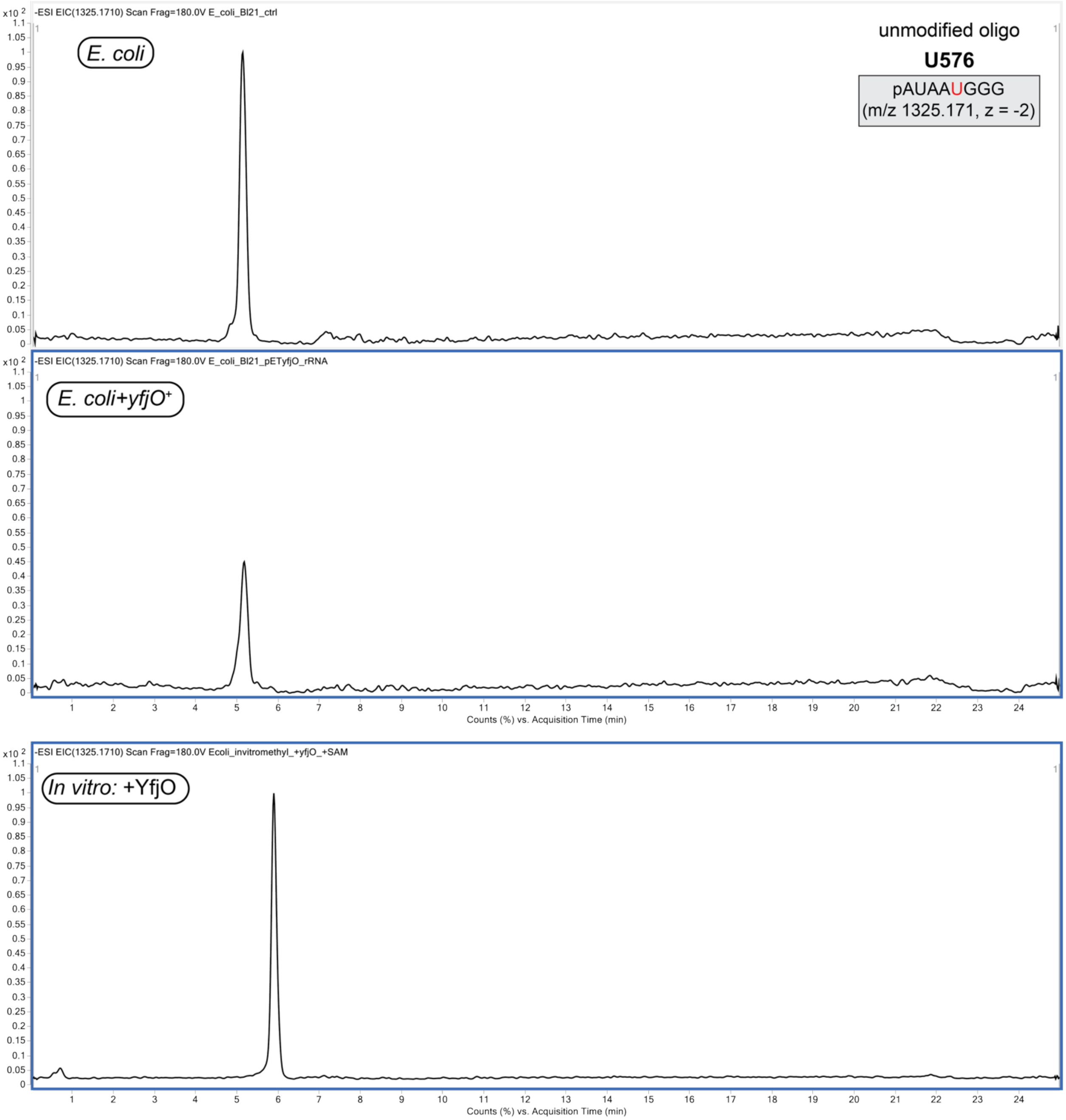


**Figure S15.** Double-cleavage oligonucleotide LC-MS analysis of *E. coli* 23S rRNA. Extracted ion chromatograms of unmodified oligonucleotide fragment produced from sequence-guided RNase H cleavage of 23S rRNA isolated from control *E. coli* BL21, YfjO-expressing *E. coli* (*E. coli +yfjO*$^{+}$), and after *in vitro* methylation reaction between recombinant YfjO and isolated *E. coli* 23S rRNA.

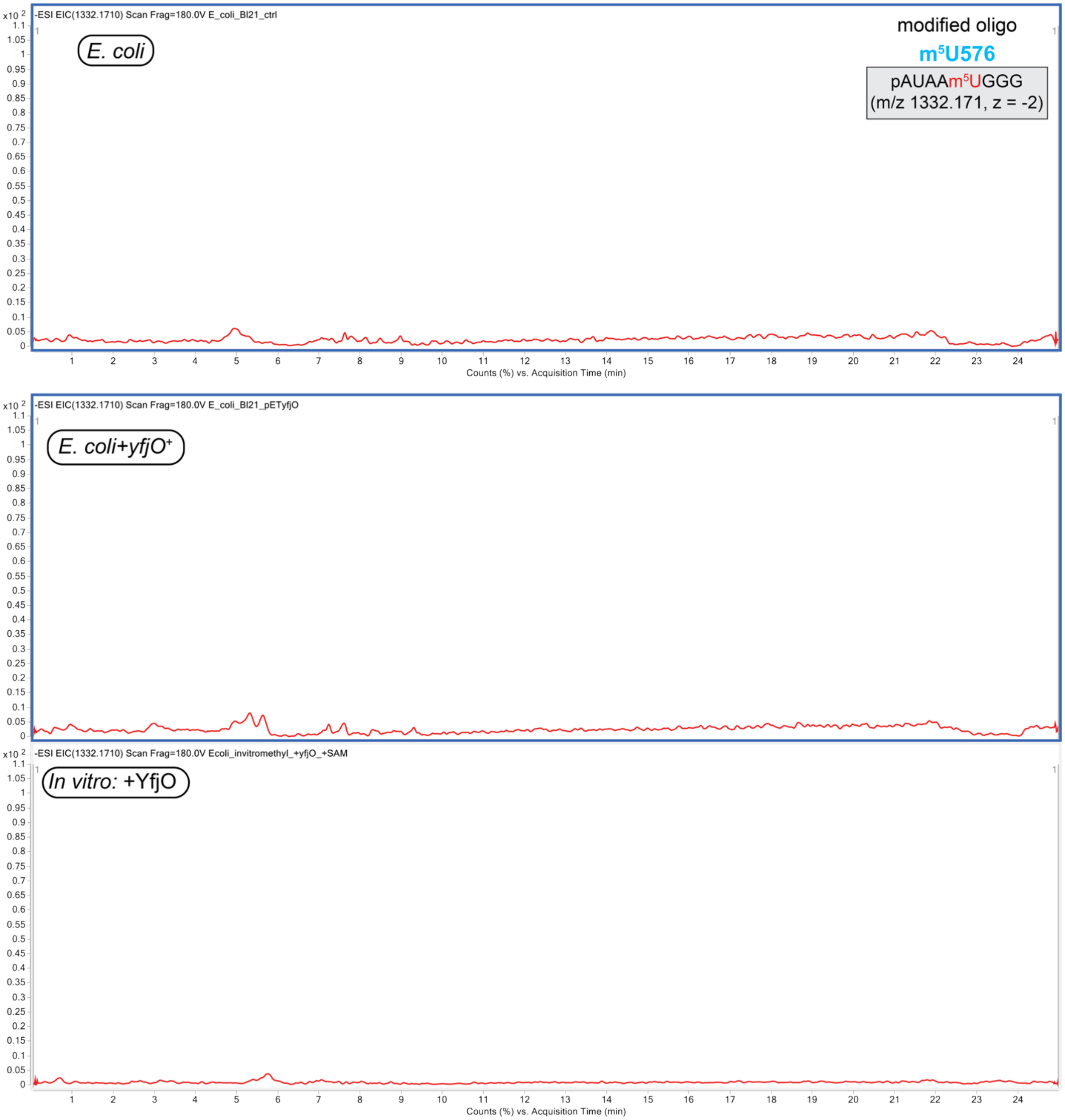


**Figure S16.** Double-cleavage oligonucleotide LC-MS analysis of *E. coli* 23S rRNA. Extracted ion chromatograms of m$^5$U-modified oligonucleotide fragment produced from sequence-guided RNase H cleavage of 23S rRNA isolated from control *E. coli* BL21, YfjO-expressing *E. coli* (*E. coli +yfjO*$^+$), and after *in vitro* methylation reaction between recombinant YfjO and isolated *E. coli* 23S rRNA.

### *E. coli* 23S rRNA

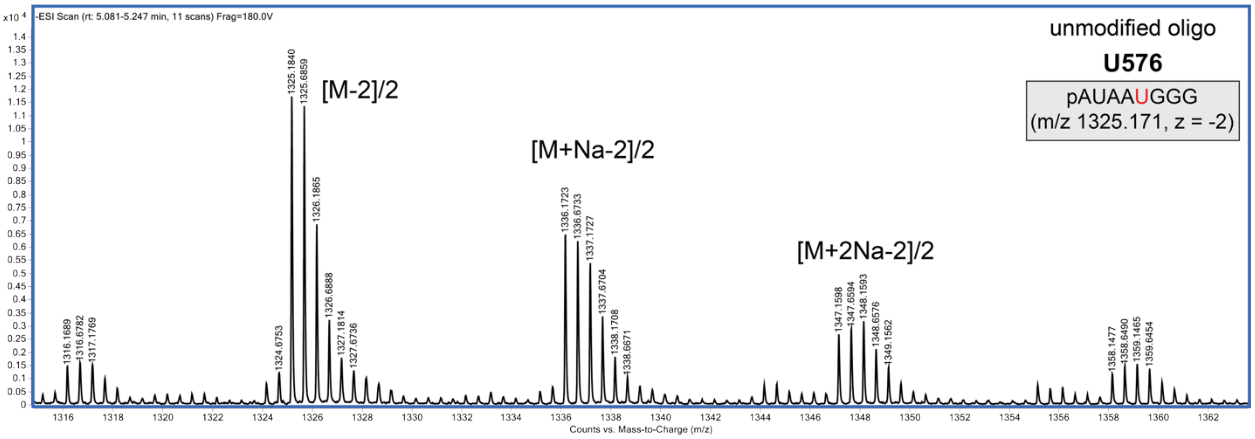


**Figure S17.** Representative MS spectra of oligonucleotide fragments produced from sequence-guided RNase H cleavage. Unmodified oligonucleotide from *E. coli* 23S rRNA.

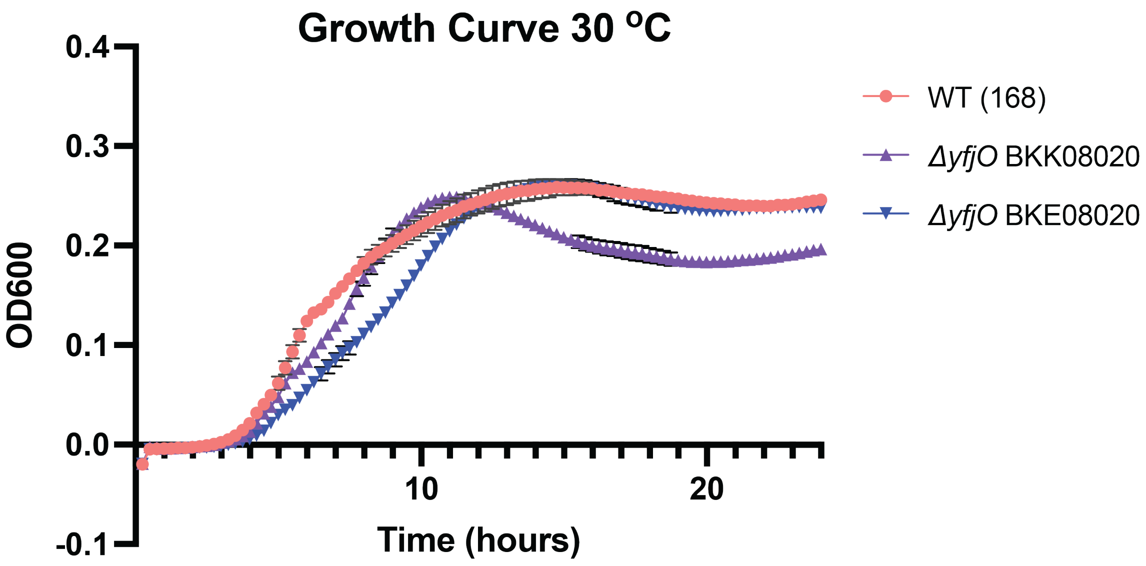


**Figure S18.** Growth curves for *B. subtilis* WT and two *ΔyfjO* strains from Gross laboratory collection; *ΔyfjO::kan* (BKK08020) and *ΔyfjO::erm* (BKE08020) at 30 °C for 24 hours. Cultures were monitored by OD600 measurement every 15 minutes for 24 hours using plate reader. Two independent biological replicates with three technical replicates each were analyzed; data represent mean ± s.e.m.

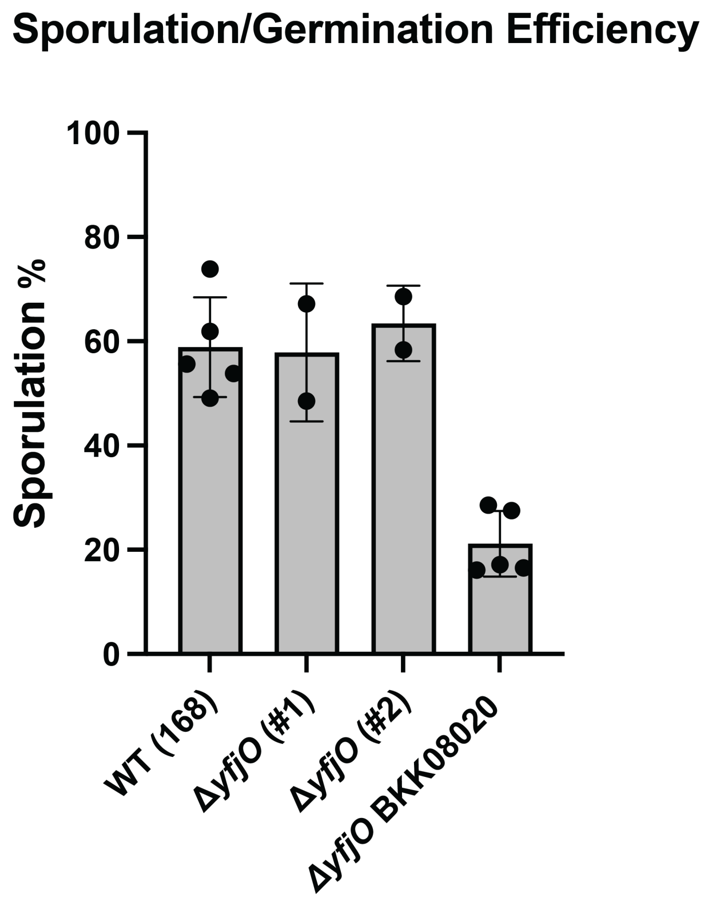


**Figure S19.** Sporulation/germination efficiency of *B. subtilis* WT, *ΔyfjO::kan* (BKK08020), and two newly reported independent clones of *ΔyfjO* was measured as the ratio of CFU from fully sporulated cultures before and after heat treatment (40 min at 90 °C). Sporulation/germination % = (CFU heat treated/CFU unheated) x 100.

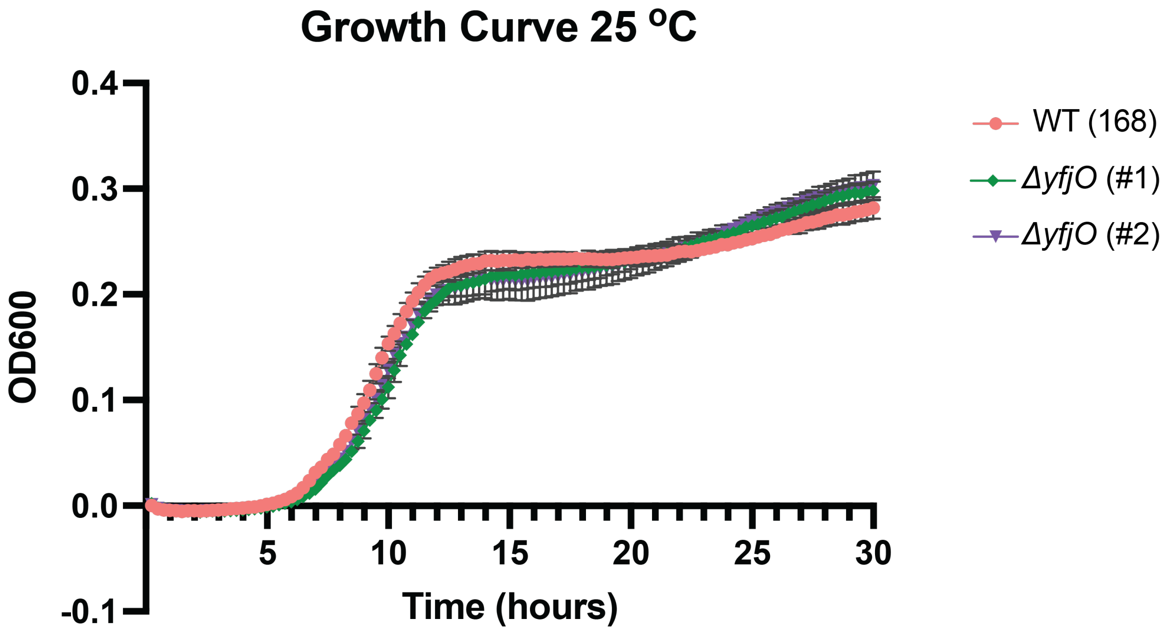


**Figure S20.** Growth curves of *B. subtilis* WT and two newly reported independent clones of *ΔyfjO* at 25 °C for 30 hours. Cultures were monitored by OD600 measurement every 15 minutes for 30 hours using plate reader. Two independent biological replicates with three technical replicates each were analyzed; data represent mean ± s.e.m.

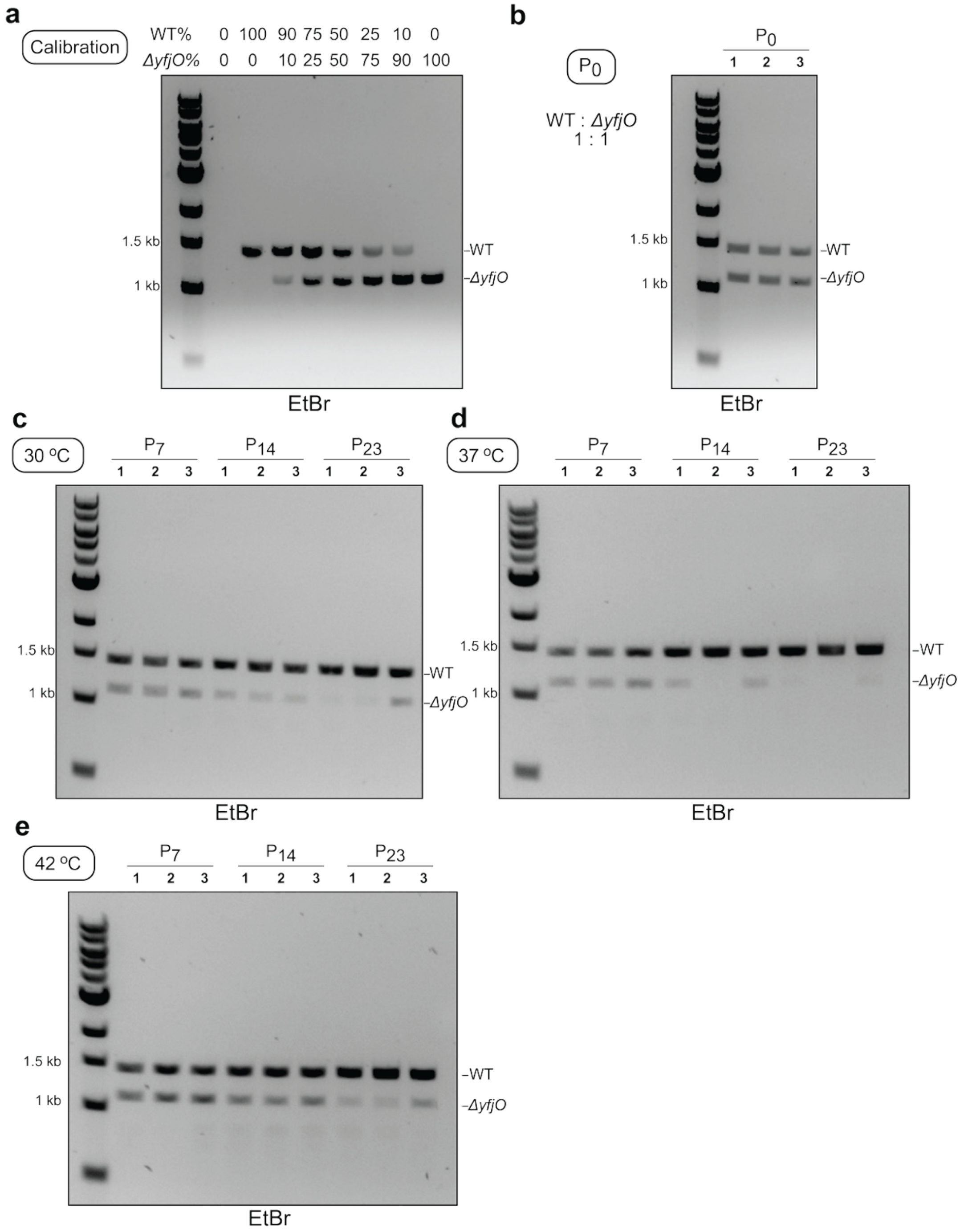


**Figure S21.** Gel analysis of competitive growth of WT versus *ΔyfjO* strains. A 1:1 mixed culture ($P_0$) was grown for 18 hours at 30 °C, 37 °C, or 42 °C, followed by serial passaging. The *yfjO* region was amplified by PCR; WT and *ΔyfjO* products are 1464 bp (upper band) and 1135 bp (lower band), respectively. **(a)** Calibration samples with defined WT:*ΔyfjO* ratios (0:100 to 100:0). **(b)** 1:1 mixed WT and *ΔyfjO* culture ($P_0$). **(c)-(e)** Passages 7, 14, or 23 were analyzed for cultures grown at 30 °C, 37 °C, and 42 °C. WT % and *ΔyfjO* % were calculated as the percentage of total densitometry signal from PCR bands.

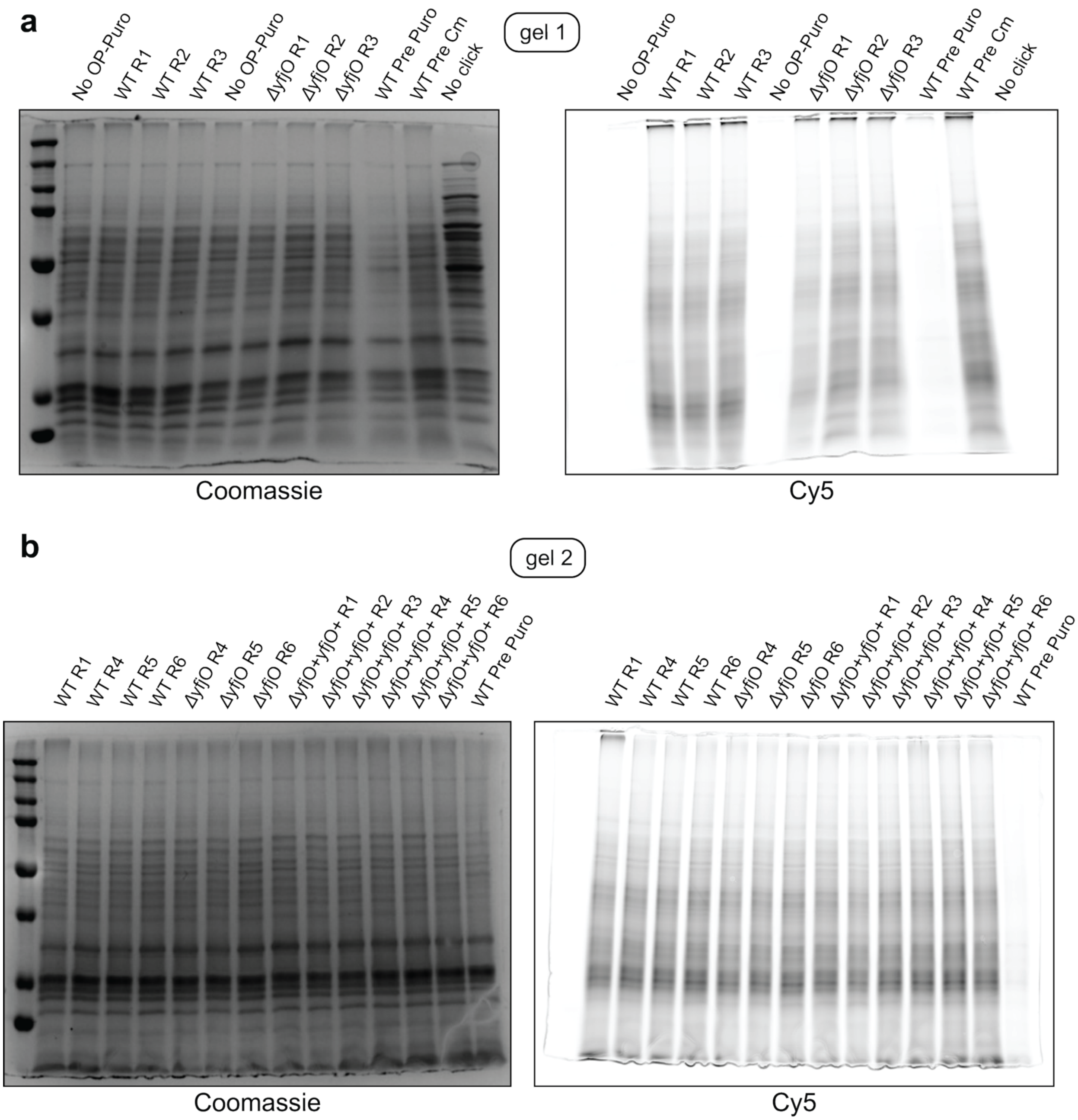


**Figure S22.** Global protein translation in *B. subtilis* WT, *ΔyfjO*, and *ΔyfjO+yfjO*$^{+}$ strains. Protein synthesis was measured using OP-Puro labelling. Representative gels are shown: Coomassie-stained (left) and fluorescent imaging (Cy5 channel, right). **(a)** First gel. **(b)** Second gel. Cultures were labeled with OP-Puro at 37 °C for 30 minutes, followed by cell lysis and CuAAC click chemistry with Cy5-azide. Pre Puro, pretreatment with puromycin; Pre Cm, pretreatment with chloramphenicol. R, replicate.
